# An autonomous system for multi-objective continuous evolution at scale

**DOI:** 10.64898/2026.03.02.709196

**Authors:** Ryan M Boileau, Stefan M Golas, Qianni Ma, Bowen Jiang, Aradhana, Mary Jia, Nika Ilieva, Aidan Baydush, Hanyu Fu, Emma J Chory

**Affiliations:** Department of Biomedical Engineering, Duke University, Durham, NC, USA; Center for Advanced Genomic Technologies, Duke University, Durham, NC, USA; Center for Biomolecular and Tissue Engineering, Duke University, Durham, NC, USA; Center for Quantitative Biodesign, Duke University, Durham, NC, USA

**Keywords:** Directed evolution, protein engineering, phage-assisted continuous evolution, synthetic biology, lab automation

## Abstract

Natural evolution is high-dimensional; organisms adapt to many pressures at once, across substrates, environments, and genetic backgrounds. Yet most directed evolution methods flatten this landscape to a single selection axis, hiding tradeoffs, and limiting what can be learned. Phage-assisted continuous evolution (PACE) is uniquely suited for multivariate selection because horizontal gene transfer couples genotype to propagation and allows the same phage lineage to traverse different selection environments. In practice, implementing this at scale has been prohibitive because each selection demands its own host culture, and every culture must be held for days to weeks within a narrow, infectable density window using continuously responsive bioreactors. In this work, TurboPRANCE is presented as an open-source, queueable robotic platform that integrates ∼200 independently controlled turbidostats with 96 parallel PACE lagoons under closed-loop control. Each turbidostat operates as a fully separate unit that can be equilibrated and initiated on its own schedule, enabling asynchronous starts and sustained operation without intervention. Automated media formulation, programmable dosing, on-deck sterilization, and adaptive scheduling coordinate growth control with the changing needs of the robotic workflow, dynamically adjusting dilution and transfer timing around formulation, sampling, and handling steps to keep each culture at consistent infectable densities despite unpredictable method demands. Cultures can be multiplexed and titrated into lagoons at defined ratios, swapped in and out on a schedule, or kept fully separate across experiments, creating a combinatorial space of selection pressures and programs that is effectively unbounded. Additionally, to enable high-throughput evolutionary tracking that scales with TurboPRANCE, Nanopore long-read sequencing was combined with DeepVariant, a deep learning-based variant caller, enabling population-level tracking of evolving variants. The result is a system that generates high-resolution time-resolvable evolutionary trajectories and large parallel datasets spanning diverse selection regimes, yielding dense, multivariate training data to map and engineer complex fitness landscapes at scale.

Turbidostat, Phage, and Robotics-Assisted Near Continuous Evolution (TurboPRANCE)
In phage-assisted continuous evolution (PACE), biomolecular activity is coupled to pIII expression, linking function to phage propagation. By altering the host strain, distinct selection pressures can be imposed on the same evolving phage population. In TurboPRANCE, ∼200 selection programs can vary over time, including periodic “drift” (mutagenesis), alternation between pressures, or rotational reassignment of host sources, enabling a combinatorial space of selection pressures.

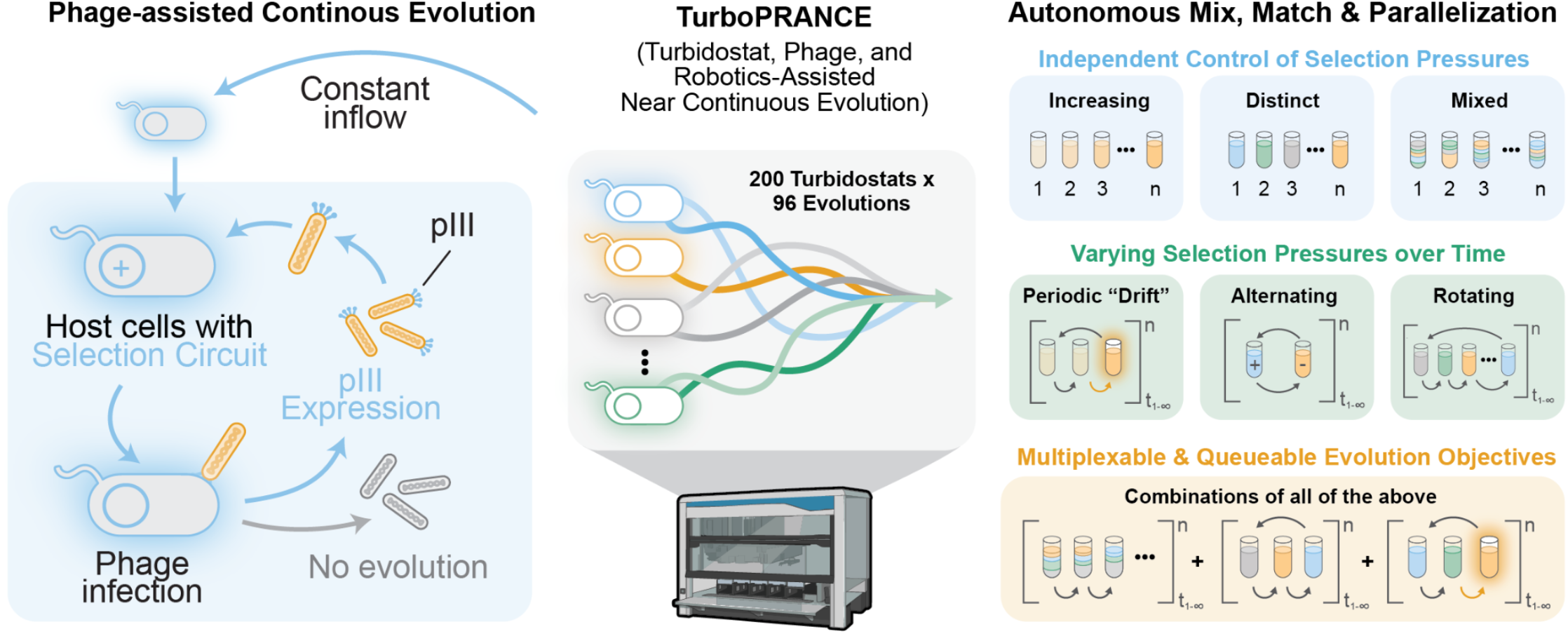

## Introduction

For over a billion years, evolution has performed a vast, ongoing experiment in multi-objective optimization, navigating hundreds of competing selection pressures at once. Each new biomolecule must simultaneously satisfy multiple performance criteria ranging from catalytic activity, affinity, specificity, stability, host burden, toxicity, and environmental robustness. As a result, most evolutionary solutions reflect local optima in a high-dimensional fitness landscape— not because Darwinian selection is incapable of finding stronger activities, rather, because being “too good” at one property can compromise fitness when making strategic trade-offs in others. In the lab, directed evolution navigates this multi-dimensional challenge by imposing an explicit selection that links molecular function to enrichment, thereby enabling optimization of a desired trait over short experimental timescales. As such, directed evolution has transformed protein engineering, yielding industrial enzymes tuned for harsh process conditions, catalysts with altered substrate scope or selectivity, protein-based probes and reporters with improved performance, and binding proteins with tailored affinity and specificity. In most implementations of directed evolution, mutant libraries are enriched for the “best” variants after iterative cycles of mutagenesis and screening^1^. These methods are configured as goal-oriented processes, where each discrete enrichment round is evaluated against an engineered readout, reducing selection to a single experimental axis. Even though natural systems such as immune defenses evolve under overlapping pressures across many substrates and contexts, most experimental evolution strategies compress this multivariate landscape into a single readout, yielding sparse, one-dimensional data that can obscure the underlying fitness surface.

One route to higher-dimensional information is through continuous evolution, where many rapid successive evolution rounds add a time axis: replication rate becomes a readout and adaptation can be tracked through kinetics and trajectories, rather than collapsed into a single endpoint. These approaches couple a molecular activity to the replication rate of a genetic carrier, such that beneficial variants can rise in frequency over time. In Phage-Assisted Continuous Evolution (PACE), the activity of a biomolecule of interest is coupled to expression of a phage gene essential for infectivity (e.g. pIII)^2,3^. Diversity is supplied by engineered host bacteria with extremely high mutation rates (∼2 bases/kb per replication)^4^ enabling the system to explore vast regions of sequence space over many generations^5^ (producing between 10^8^-10^14^ unique variants every 10 minutes). Continuous diversification ensures that only functional variants trigger phage replication, allowing beneficial mutations to rapidly emerge and out-compete non-functional sequences. Unsurprisingly, phage-assisted evolution has produced some of the field’s most impactful molecular tools, including evolved proteases^6,7^, polymerases^2,8^, high-affinity protein interactors^9–11^, DNA-protein interactors^12,13^, a diversity of metabolic and biocatalytic enzymes^14–16^, small molecules^17^ and antibiotics^18,19^, base and prime editors^20–23^, PAM-flexible Cas nucleases^24,25^, integrases^26,27^, and CRISPR-associated transposases^28^. PACE has also served as a robust experimental system to quantify deterministic and contingent outcomes^29,30^, map how path-dependent trajectories^31^ are shaped by mutation rate and selection stringency^32^, and can expose broad structure-function constraints, ranging from solubility limits^10^ to the emergence of drug resistance^6^.

The viral-host replication framework of PACE also offers unique advantages over earlier evolution approaches such as phage display^33^, yeast display^34^ (which require intermittent sorting of “winners” and “losers” each round^35^), and higher-order continuous evolution methods such as OrthoRep^36^ (a yeast-based platform that mutates the target gene on an engineered orthogonal chromosome). A key advantage is that in display-based techniques, each new round of screening requires rebuilding a library (via epPCR, recloning, etc.), or each type of selection pressure (via cloning and retransformation). In contrast, in PACE, enriched phage can be used to initiate a subsequent experiment without reconstructing the variant pool. In prior studies, the power of this feature has been demonstrated by sequentially enriching and re-infecting host strains encoding progressively stronger promoters^6,31^, more divergent substrates^6^, or by alternating between positive and negative selection pressures^16,37^. However, PACE requires a dedicated continuous-flow turbidostat to keep host bacteria at a steady, infectable density, since phage propagation^2^ and sustained mutagenesis^4^ rely on maintaining a narrow OD_600_ window (typically 0.4 to 0.6^4^). Thus, multiplexing strains, circuits, and selection regimes across parallel bioreactors remains technically limiting in practice.

Previously, the system PANCE (phage-assisted non-continuous evolution) enabled microplate-format evolution^3,14^ and replaced continuous flow with repeated, manual re-infection cycles. However, keeping cultures within the narrow infectable window to sustain mutagenesis at each daily transfer is labor-intensive and can be difficult to reproduce. As a result, PANCE yields fewer effective viral generations, which can be limiting for complex evolutions or those that require tight control of component concentrations, kinetics, or metabolite availability. These technical limitations motivate the use of PANCS (phage-assisted non-continuous selection)^11,17^, a more user-friendly method which efficiently enriches variants from highly diverse starting pools (including de novo libraries)^38^. In a complementary approach, ePACE^24^ (PACE paired with eVOLVER^39^) parallelizes multiple PACE experiments using arrays of chemostats (i.e. fixed flow rate reactors), where each chemostat supplies one culture to a paired evolution lagoon. However, because PACE strains’ growth rate can vary widely depending on the circuit burden, chemostat flow conditions must be pre-tuned for each strain to prevent overgrowth or dilution at steady state, and these strains cannot be readily re-directed to different lagoons without implementing complex fluidics systems. Building on the need for scalable continuous evolution, Phage- and- Robotics- Assisted Near-Continuous Evolution (PRANCE) demonstrated that automated liquid handling could sustain near-continuous selection and mutagenesis in high-throughput^29^. In PRANCE, each host strain is maintained in an off-deck turbidostat or pre-prepared as a separate stock culture, and these cultures are pumped onto a liquid handler to supply the continuous plate-based transfers in high-throughput.

In principle, horizontal gene transfer inherent to PACE enables sampling many selection pressures simultaneously, resulting in evolution that generates tailor-made molecules with multiple desirable properties. However, to harness the full potential of PACE, new platforms are still needed that can stably maintain many distinct host strains in parallel and rapidly re-route evolving phage between them. To address this need, the work developed herein presents TurboPRANCE, a high-throughput turbidostat-enabled evolution platform that maintains up to 200 independent, density-responsive flow reactors and uses them as a reusable palette to program 96 parallel PACE lagoons with distinct, user-defined selection pressures. Because each turbidostat can encode a different strain background, circuit, inducer state, or media formulation, pressures can be composed as precise mixtures, reassigned over time, and updated without interrupting the run, enabling systematic exploration of trade-offs such as activity across many substrates, performance across a ladder of selection strengths, and outcomes under defined combinations of constraints. Irrespective of circuit burden, TurboPRANCE keeps every culture in an infectable regime despite widely varying growth rates, tracks evolution progress in real time by on-deck plate-reader measurements, and can autonomously deliver supporting inputs (including helper strains, when needed) to stabilize trajectories and prevent washout. Critically, the robot is not merely a transfer device: it functions as a parallelized process controller and programmable mixing layer, routing defined culture streams into selected lagoons on a per-cycle basis to implement time-varying selection programs. The platform is designed for non-overlapping operation, allowing experiments to be staggered and new turbidostats and lagoons to be queued while other evolutions remain underway. Together, TurboPRANCE turns PACE into a mix-and-match, multi-objective evolution engine and makes it practical to map many overlapping selection landscapes at a scale that has not been previously accessible.

## Results

### System architecture & method optimization for programmable multi-pressure continuous evolution

TurboPRANCE combines 200 independent turbidostats with 96 parallel phage evolution lagoons, each supported by automated media formulation, on-deck sterilization, and adaptive control software on a Hamilton STARlet liquid-handling robot **(Fig. 1A)**. A peristaltic pump network **(Supp Fig. 1)** routes bleach, water, media, and waste between off-deck reservoirs and on-deck components **(Supp Fig. 2)**, through custom 3D-printed washer modules that segregate bleach, water, media, and waste into independent fluid channels **(Supp Fig. 3)**. The full system uses a real-time feedback layer written in Python through PyHamilton, an open-source python framework for controlling Hamitlon liquid handlers^40^. Unlike PRANCE, which relied on pre-defined timing intervals and bacteria supplied from an off-deck turbidostat^29^, TurboPRANCE maintains and controls each continuous bioreactor directly on deck. TurboPRANCE operates in two phases within a continuous loop that can be initiated independently for each culture **(Fig. 1B)**: (i) a turbidostat equilibration phase, in which individual wells are inoculated, tracked through early to mid-exponential growth, and then held at any pre-defined OD600 setpoint, and (ii) a lagoon-transfer phase, in which one or more equilibrated turbidostats are routed to specified phage lagoons at defined flow rates to initiate and sustain continuous evolution **(Supp Fig. 4)**. Because each turbidostat well transitions on its own schedule, experiments can be staggered and run side-by-side, with some wells equilibrating while others are already supplying lagoons. To support this, a predictive growth model^40^ continuously updates strain-specific growth parameters from ongoing OD measurements and adjusts transfer volumes to maintain infectable densities despite widely varying growth rates across strains and circuit architectures. Automated media formulation, fluid handling, and waste management enable sustained, unattended operation, while plate-reader measurements provide real-time monitoring of culture density and reporter output **(Supp Fig. 4)**. An interactive manifest file specifies strain identity, setpoints, flow rates, sampling, and lagoon assignments, and can be updated during operation to reconfigure selection pressures without interrupting the run **(Supp Fig. 5)**. Achieving robust multi-day sterility required iterative optimization of both tip handling and deck layout. Early configurations suffered from stochastic turbidostat death caused by bleach residue transferring between tip columns during high-force ejection cycles; this was resolved by implementing a segregated dirty-tip workflow that physically isolates recently bleached tips from clean staging areas **(Supp Fig. 6)**. Separately, phage cross-contamination from lagoon sampling flyover into turbidostat tip racks was eliminated by repositioning clean tips away from the lagoon transfer corridor **(Supp Fig. 7)**, which required designing a custom single-channel washer module to handle 96-head and 8-channel flyover differences **(Supp Fig. 3)**.

**Figure 1.**
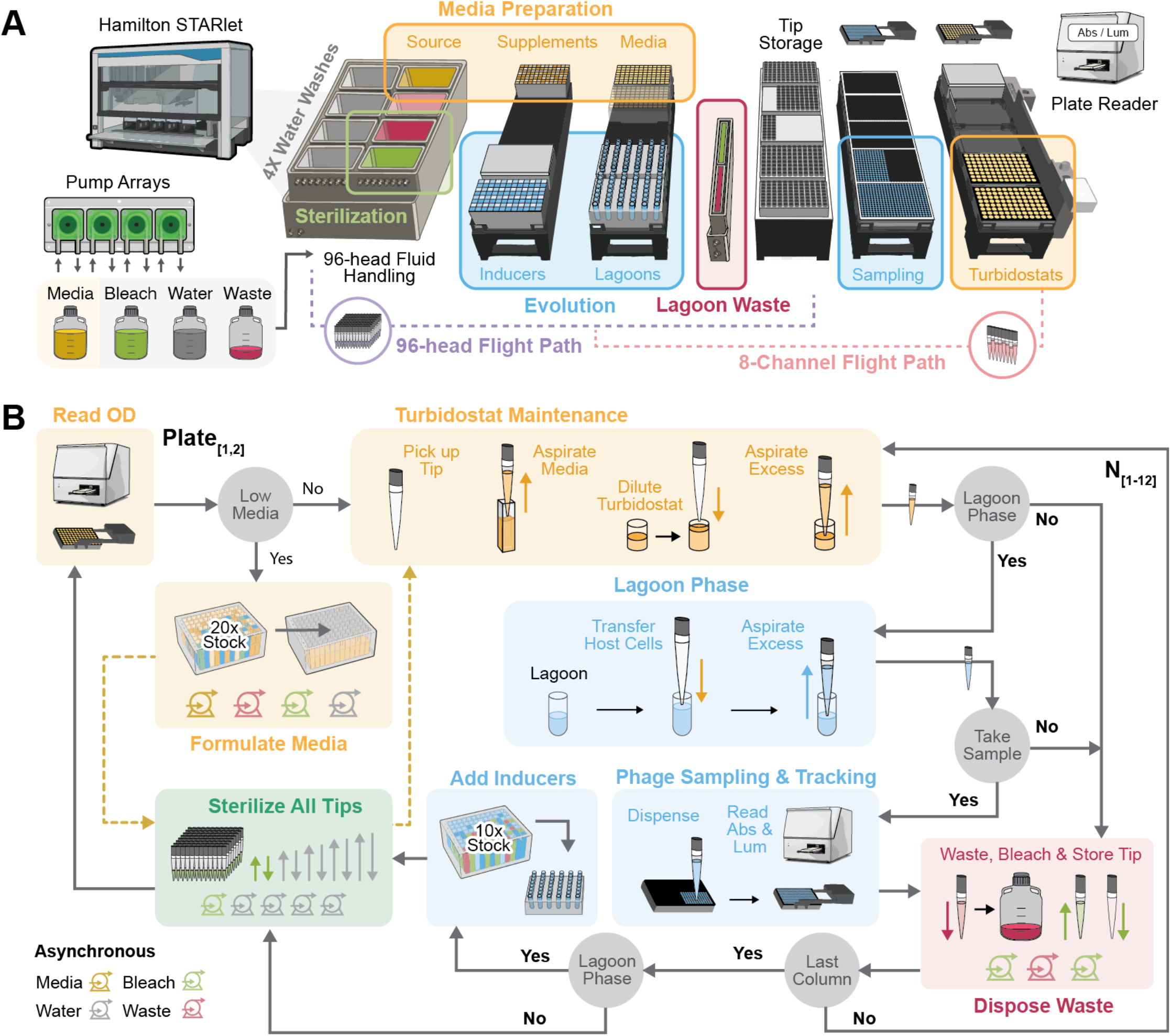
TurboPRANCE Overview and Method Architecture. **(A)** A liquid handler integrates ∼200 independently controlled turbidostats with 96 parallel evolution lagoons. Bulk media, water, bleach, and waste reservoirs are connected through peristaltic pump arrays to on-deck washer modules and a media reservoir. The method performs media formulation, turbidostat dilution, lagoon transfers, inducer addition, sampling into 384-well plates, and plate-reader measurements. Separate 96-channel and 8-channel workflows, together with spatial separation of sterile and contaminated components, enable continuous multi-day operation. **(B)** Each cycle starts by reading the next turbidostat plate. Media is refilled if needed. Turbidostats are then serviced column-wise (N = 1-12): pick up tips, aspirate media, dilute, and remove excess. If lagoon phase is active, a defined fraction of excess is routed to mapped lagoons (with volume rebalanced), and lagoons can be sampled into 384-well plates before inducer addition. Tips are bleached and returned, waste is disposed, and the cycle repeats.

During development of the multi-day evolution campaigns, a number of challenges were observed not previously encountered in conventional turbidostats, which typically run for only 24-48 hours and grow bacteria alone. In TurboPRANCE, cultures must be sustained for days, and despite the deletion of multiple bio-film associated genes in S2060^12^, biofilm accumulation is regularly observed in PACE experiments **(Supp Fig. 8A)**, which can be mitigated by adjusting the height of mixing **(Supp Fig. 8B)**. However, overly aggressive pipette mixing shears the F-pilus required for M13 phage infection. It was observed that no mixing (shaking only) causes rapid biofilm accumulation and culture overgrowth **(Supp Fig. 9A)**. However, it was also observed that mixing at regular intervals is prohibitive to phage propagation, so the method was carefully tuned to allow for regular turbidostat maintenance without compromising phage propagation **(Supp Fig. 9B)**. A further constraint arises from slow-growing or burdened strains that cannot individually sustain the volumetric outflow required for continuous lagoon transfer. To address this, TurboPRANCE supports column batching, in which multiple turbidostat wells carrying the same strain are serviced as a single unit and their excess culture is pooled before delivery to the lagoon. This increases the total available host volume per cycle without changing per-well setpoints or control logic, enabling higher lagoon flow rates even when individual cultures grow too slowly to meet the demand from a single well **(Supp Fig. 10)**. The tradeoff is reduced strain capacity per plate, but batching provides a practical route to robust operation for burdensome strains when high continuous flow rate is required for sufficient selective pressure.

### Adaptive cycle-time projections are required for simultaneous turbidostat and lagoon operations

At its core, a fundamental challenge of TurboPRANCE is stable turbidostat maintenance. Maintaining each lagoon within an operational density range to support continuous infection requires keeping each turbidostat at a defined OD setpoint while simultaneously supplying a steady outflow of uninfected bacteria to the lagoons at the flow rates required for continuous evolution (typically on the order of 1 vol/hr). This creates a timing constraint: each culture must be revisited for dilution before growth drives it beyond the allowable density band. Unlike a classical chemostat, where dilution occurs at a fixed volumetric rate and density drifts if growth rate deviates from the tuned flow condition **(Fig. 2A) (Supp Fig. 11A)**, TurboPRANCE must repeatedly return to each culture at the correct moment to maintain a defined OD setpoint. In PANCE, long discrete intervals inherently permit repeated overshoot between rounds reducing mutagenesis capability and requiring a fresh culture. **(Fig. 2A) (Supp Fig. 11B)**. In original PACE, this is circumvented using an idealized continuous turbidostat which converges smoothly to the setpoint under instantaneous measurement and feedback **(Fig. 2A) (Supp Fig. 11C)**. However, on a liquid handler turbidostat control is necessarily discrete, and fixed-interval servicing produces growth-rate-dependent sawtooth oscillations around the setpoint **(Fig. 2A) (Supp Fig. 11D)**. When service intervals vary, the controller underestimates the time until the next dilution (even by a few seconds), cultures will thus overshoot the setpoint resulting in periodic over and under-dilution **(Supp Fig. 11E)**. Continually overshooting an OD setpoint not only reduces infectability but, in severe cases, requires dilution volumes that exceed the robot’s pipetting capacity **(Fig. 2A)**. A key challenge is that unlike most automation protocols executed on a liquid handler, Δt is not a fixed property of the method **(Fig. 2B)**. Instead, it is an emergent property of how long the robot takes to complete the set of operations demanded by a future state of the system before it can return to a given well, which is highly variable across strains, the number of active turbidostats and lagoons, and changes over time **(Fig. 2C)**. Different strains also grow at different rates and impose different operational loads (pipetting time, media refill requirements), and these demands change as cultures equilibrate, as lagoon transfer begins, and as sampling schedules shift. Control therefore requires accurate prediction of the elapsed wall-clock time between current and future dilution events (Δt_est_), as Δt cannot be reliably hard-coded from a list of planned submethods. To overcome this, the platform adapts to operational demands that change over time, including pipetting volumes (early low-volume dilutions versus later high-volume maintenance), media refill timing driven by the fastest-growing wells, and optional sampling and reporter readouts that can vary from well to well.

**Figure 2.**
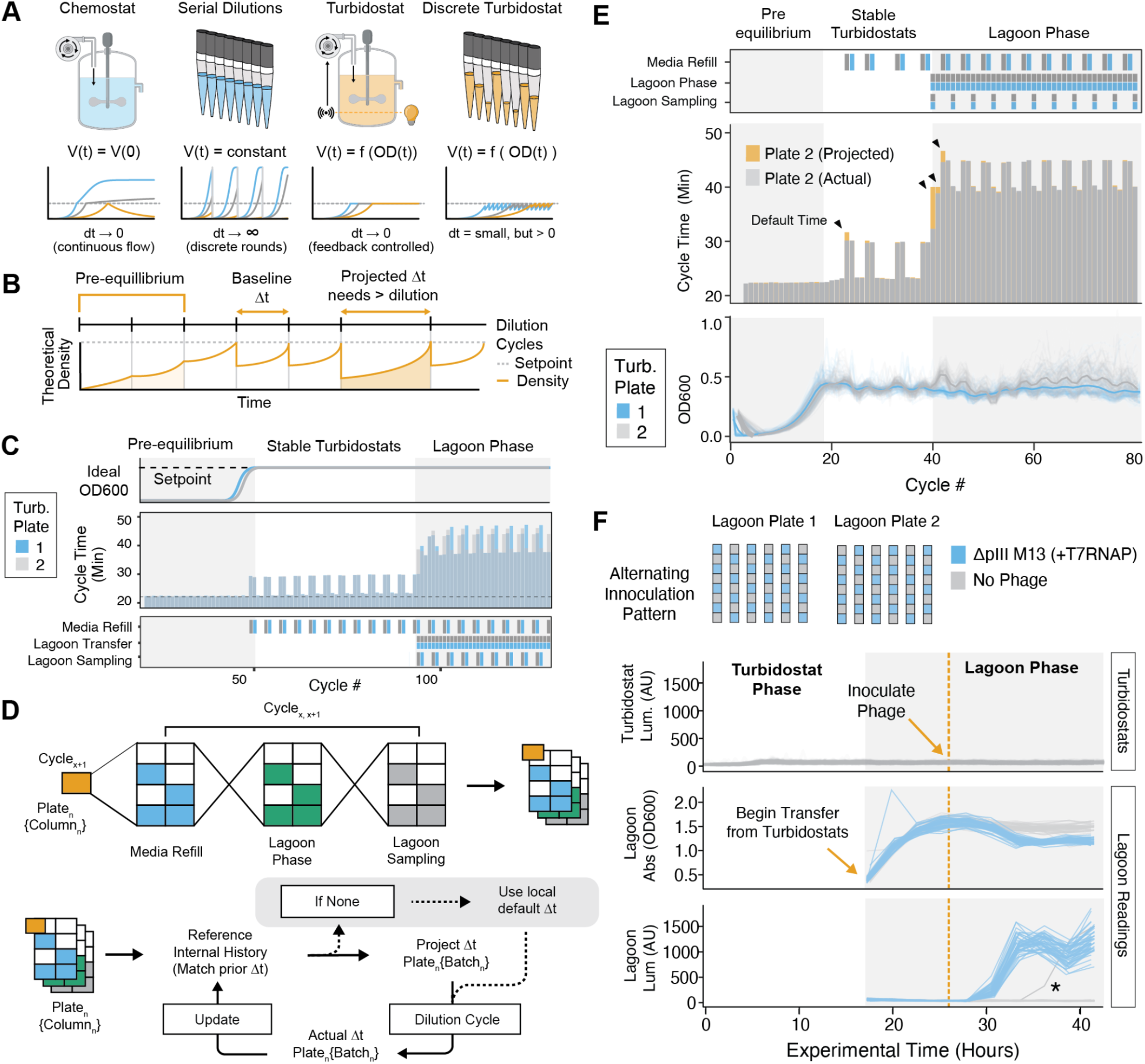
Adaptive cycle-time projections are required for simultaneous turbidostat and lagoon operations. **(A)** Comparison of chemostat, turbidostat, serial dilution, and fixed-interval discrete turbidostat culture control. **(B)** Accurate Δt (time between dilutions) is required to maintain turbidostat OD setpoints; underestimation causes OD overshoot and can require dilution volumes beyond pipetting capacity. **(C)** Different method requirements change cycle spacing, so the controller must forward-project how long the next interval will be in order to compute the correct dilution volume to return cultures to their setpoints as lagoon transfer, sampling, and media refill turn on and off. **(D)** TurboPRANCE estimates Δt per plate and batch, then adapts the estimate using the most recent observed robot execution times under matching conditions. **(E)** Yellow indicates projected cycle times that converge to measured cycle times as the robot workload changes; turbidostat OD measurements stabilize across phase transitions under adaptive timing. **(F)** Validation run propagating T7RNAP phage in pT7-pIII-LuxAB host lagoons: no luminescence is detected in the upstream turbidostats (no back-contamination), while inoculated lagoons show increased luminescence and a depression in lagoon density (absorbance) consistent with infection. One control lagoon shows delayed luminescence (*), consistent with a manual tip contact event during inoculation.

It was observed that three key sources of variability make fixed cycle timing particularly unreliable **(Supp Fig. 12)**. First, pipetting time depends on volume, and the required dilution volumes shift substantially between early growth and steady-state maintenance, introducing minute-scale changes in cycle duration **(Supp Fig. 13)**. Second, unpredictable operations such as media re-formulation needs, transition into lagoon phase (which can be autonomously initiated upon turbidostat set-point acquisition), and lagoon sampling frequency occur intermittently and add variable overhead, (Cycle_X+1_ and Cycle_X+2_) **(Fig. 2B-C) (Supp Fig. 13)**. Third, media refilling is triggered by consumption, which is most impacted by the fastest-growing wells, so refill events occur at irregular times that differ between runs and strain compositions **(Supp Fig. 13B-D)**. To address this, TurboPRANCE uses a fully adaptive cycle-time projection strategy specifically to preserve stable turbidostat maintenance under these variable conditions, rather than a semi-adaptive approach in which only media refill timing is updated dynamically while lagoon and sampling increments remain fixed **(Supp Fig. 14)**. Users specify initial default times for possible operation combinations, but the system then records observed cycle durations and updates Δt estimates in real time based on the most recent measured execution times, accounting for the operations performed in adjacent cycles. This adaptive timing is tracked at the level of individual batches (for example, plate columns during sampling), since different batches can diverge by several minutes in effective cycle time even within the same plate **(Supp Fig. 15)**. Because these timing fluctuations arise from both task composition and hardware performance, their magnitude and structure can differ across robotic platforms, further undermining any assumption of a fixed, globally valid cycle time.

### Localized software configurations facilitate varying TurboPRANCE setups

Due to observed variability between hardware configurations, TurboPRANCE was also designed to be system-agnostic and portable across physical setups by separating hardware-specific parameters from the core control logic. Robustness was evaluated by deploying the same codebase on two distinct TurboPRANCE systems which were assembled with different hardware configurations **(Supp Fig. 2)**, including a new-model and a refurbished Hamilton STARlet 2020 liquid handler with different channel types (CORE I vs. CORE II), and different plate readers (OMEGAstar versus of VANTAstar) **(Supp Fig. 16)**. Operation across these platforms required accommodating several platform-specific differences while preserving identical cycle logic and controller behavior. First, the two liquid handlers differed in internal X/Y/Z calibrations and offsets. This necessitated separate deck layout files (plate locations, tip washer positions, and access points) and platform-specific calibration of pipetting Z-heights, which were required to maintain constant working volumes in both turbidostat wells and lagoons. Second, the refurbished liquid handler exhibited minutely, but quantifiably slower mechanical throughput (pipetting, gripper movements, and overall command execution), reducing achievable cycle speed. After initial trial runs, default projection times were derived and used as starting values for adaptive projection timing on each setup. Third, plate reader integration can differ in both software and geometry: older readers can require compatibility with legacy control software, can have different plate orientations, or require different calibration curves to convert absorbance to turbidostat and lagoon density estimates. Finally, the peristaltic pump arrays differ between setups (tubing lengths and pump speeds), requiring recalibration of delivered volumes for fluid handling steps. Across these variables, TurboPRANCE was built to load robot-, reader-, and pump-specific parameters from local configuration files, enabling cross-platform operation without modifying core method logic. Both hardware configurations were used for TurboPRANCE experiments presented throughout the remainder of this study.

To quantify TurboPRANCE’s operational envelope for fast-growing strains, we cultured 96 different strains from the Keio Knockout Collection in DRM medium without antibiotics, a condition that maximizes growth rate by eliminating plasmid burden and selection pressure. Steady-state growth rate estimates (k) ranged from 0.95 to 1.6 hr^-1^ (doubling times of 25-45 min), with a mean of 1.3 +/-0.1 hr ^-1^ **(Supp Fig. 17)**. These growth rates overlap with but greatly exceed those of typical PACE host strains, which carry accessory plasmids and are maintained under antibiotic selection (∼0.4-1.0 hr ^-1^, doubling time ∼ 40-100 min). The maximum growth rate that TurboPRANCE can stably maintain is set by the constraint that bacteria must not outgrow the maximum pipettable dilution during a single service cycle. In turbidostat-only mode, cycle times alternate between approximately 20 and 30 min, yielding a maximum supportable growth rate of k = 1.7 hr ^-1^. When operating with a full complement of 96 lagoons, cycle times increase to approximately 35-45 min due to regular media prep, sampling, and additional fluid-handling steps required for lagoon transfers. Under these conditions, the maximum supportable growth rate drops to k = 1.13 hr ^-1^, and 93 of 96 Keio strains exceed this growth rate limit. Importantly, this throughput constraint can be managed by reducing the number of active lagoons. Because the robot processes wells in batches of 8, removing either turbidostats or lagoon columns directly shortens the cycle, so the method is readily adaptable to rapidly growing strains, including those that support phage selection, rather than evolution. For typical PACE applications, where host strains grow at k∼0.4-1.0 hr ^-1^ due to plasmid maintenance and antibiotic selection, these growth rates are comfortably supported even at high lagoon counts. Growth rate simulations establish that TurboPRANCE can operate with fast-growing, low-burden strains by reducing lagoon count, expanding the method’s applicability beyond standard PACE host backgrounds.

### Compartmentalization of turbidostat and phage propagation

Following system buildout and validation, an enrichment experiment (without mutagenesis) was performed using a published high-activity circuit in which phage-encoded T7RNAP drives transcription from a host accessory plasmid containing promoter for T7 (pT7) upstream of gIII and LuxAB. Freshly inoculated turbidostat wells were equilibrated to OD 0.4 within 7 h and were maintained for an additional ∼10 h **(Fig. 2E)** before initiating continuous transfer of host bacteria to lagoons at 0.5 vols/hr. After 10 h of lagoon transfer, lagoon absorbance stabilized, and M13-T7RNAP phage was manually introduced at a 1:10 multiplicity of infection (MOI). Phage were added to half of the lagoons in a checkered pattern to assay cross-contamination between neighboring conditions **(Fig. 2F)**. Luminescence increases were observed in phage-spiked lagoons by the second sampling timepoint (3 h) and continued rising over multiple timepoints before peaking and modestly decreasing **(Fig. 2F)**, as previously observed by DeBendedictis & Chory et al ^29^ bult without a clear mechanistic explanation. A stochastic Monte Carlo model of infection in a continuously diluted lagoon recapitulated the transient luminescence peak and identified its origin as a balance between phage spread and continuous dilution with uninfectable host bacteria **(Supp Fig. 18)**. When effective MOI is high relative to host inflow, infection transiently saturates the lagoon before continuous replenishment of uninfected cells restores a lower steady state. At higher flow relative to MOI, saturation does not occur and infection approaches a reduced plateau. These dynamics explain the observed luminescence overshoot as a transient imbalance between propagation and dilution. Additionally, although the absorbances of lagoons become nonlinear above ∼1.5 OD600, infected lagoons exhibited a pronounced absorbance depression during infection, as previously observed^29^. No luminescence increase was detected in unspiked control lagoons or in the turbidostats throughout the experiment, with one exception: a single control lagoon showed delayed luminescence **(Fig. 2F)**. This event coincided with an observed manual handling error in which a multichannel tip contacted the rim of that control well during phage addition to adjacent lagoons, underscoring the contamination risk between neighboring lagoons. Despite this, luminescence remained absent from all other control lagoons and from the turbidostat array, supporting effective containment during routine operation and the efficacy of the tip-washing and layout features intended to prevent carryover and flyover of phage-contaminated liquid **(Supp Fig. 19)**. Together, these results demonstrate stable, self-sustained, and compartmentalized parallel operation of turbidostats and lagoons at high throughput.

### Evaluating multivariate evolution parameters

Following validation of faithful phage propagation across lagoons, TurboPRANCE was applied to evolution campaigns. Because PACE experiments have historically been lower throughput, published results rarely deviate from default conditions, including the turbidostat set points (OD600 0.4-0.6), lagoon flowrate (1 vols/hr), and the multiplicity of infection used to seed evolution (1 phage to 10 bacteria). Each parameter was systematically varied using a well-characterized selection that evolves T7RNAP for T3 promoter (pT3) activity **(Fig. 3A)**. For these experiments and all subsequent experiments shown, phage were added to lagoons prior to initiating bacterial transfer, resulting in infection immediately following the first transfer event. To enable direct comparisons across parameter sets the initial transfer volume into lagoons was also standardized across conditions. Additionally, to reduce any impacts from biofilm accumulation in turbidostat wells, approximately 90% of each turbidostat culture was periodically transferred by multichannel pipette into a fresh plate without interrupting the run. Increases in luminescence are reported here as a proxy for evolutionary progress, and evolution outcomes are saved and sampled for downstream processing.

**Figure 3.**
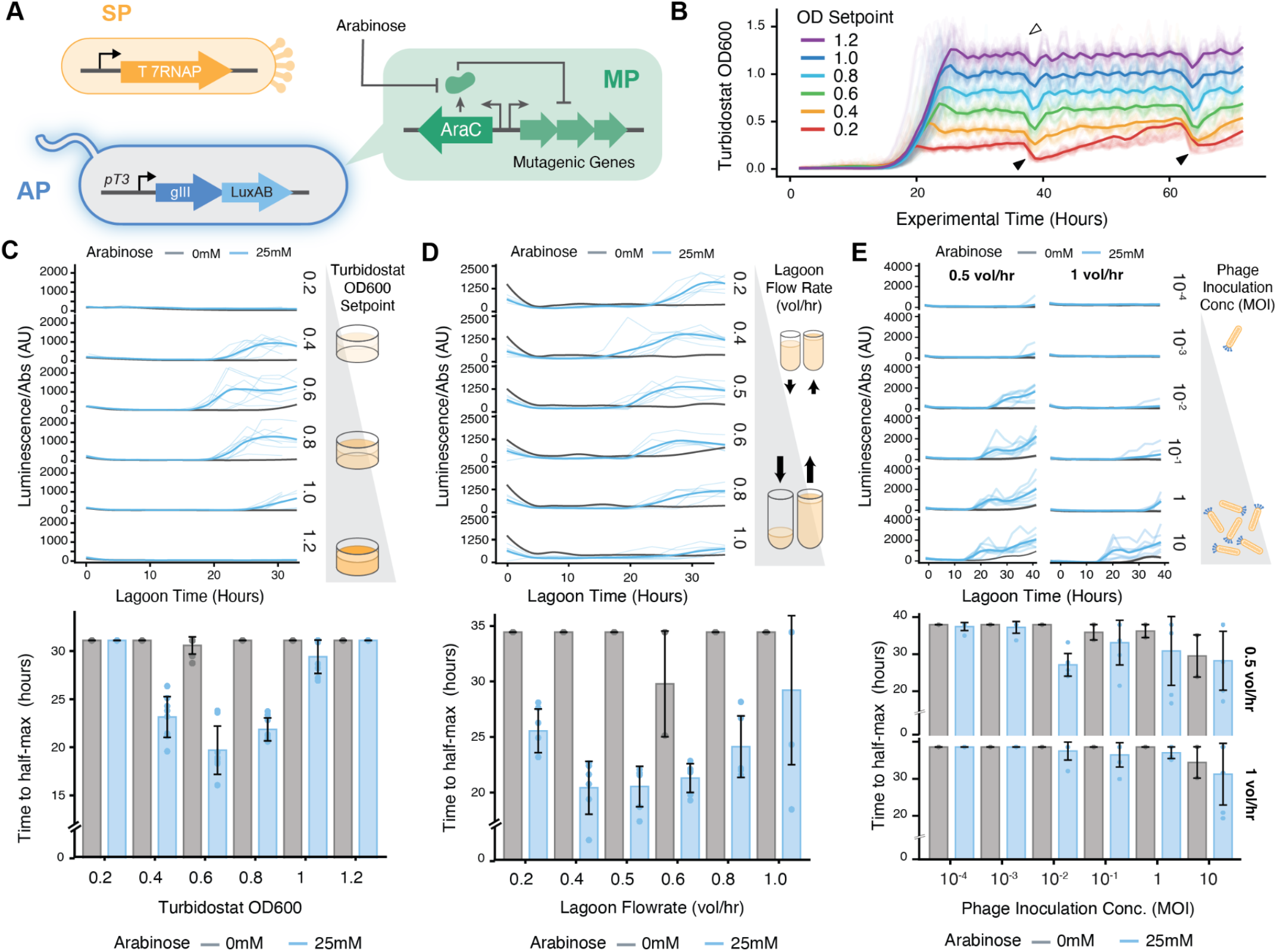
Multivariate parameter exploration in TurboPRANCE. **A)** Circuit schematic linking phage-encoded activity to pIII-LuxAB expression from a pT3-driven accessory plasmid, with an arabinose-inducible mutagenesis plasmid (MP). **(B)** Turbidostat OD600 trajectories across setpoints (0.2 to 1.2) during equilibration and maintenance (32 replicates per setpoint). **(C)** Top: Turbidostat OD setpoint sweep (0.2 to 1.2) showing lagoon luminescence normalized to absorbance over time under 0 mM or 25 mM arabinose. Bottom: Corresponding time to half-max luminescence (t50) summarized versus setpoint (8 evolutions, 8 controls per setpoint). **(D)** Top: Lagoon flow-rate sweep (0.2 to 1.0 vol/hr) showing lagoon luminescence/absorbance over time under 0 mM or 25 mM arabinose. Bottom: Corresponding t50 summarized versus flow rate (6 evolutions, 2 controls per flow rate). (E) Top: Phage inoculation concentration (MOI) sweep (10^-4 to 10) showing lagoon luminescence/absorbance over time at flow rates 0.5 and 1 vol/hr. Bottom: t50 summarized versus MOI for each flow rate (6 evolutions, 2 controls per MOI). Error bars represent SD.

First, the effect of turbidostat OD setpoint on T7RNAP evolution was evaluated by running turbidostats at OD setpoints spanning 0.2 to 1.2 and initiating evolutions at 0.5 vol/hr with a 1:10 MOI. TurboPRANCE maintained most cultures near their assigned setpoints throughout the experiment, including during the transition into lagoon phase **(Fig. 3B) (Supp Fig. 20A)**. Notably, turbidostats maintained at OD 0.2 exhibited faster growth than higher setpoints, consistent with shorter doubling times in early exponential phase^41^, which reduced controllability during entry into lagoon phase under longer cycle times. Lagoons were supplemented with 25 mM arabinose to induce mutagenesis and were compared to matched control lagoons without induction (0 mM arabinose). Luminescence was first detected in lagoons supplied by OD 0.6 turbidostats at approximately 16 h, followed by OD 0.8, 0.4, and 1.0 conditions **(Fig. 3C)**. By 33h, luminescence was only observed in a single lagoon supplied by OD 0.2 and 1.2 turbidostats. In the no-arabinose controls, signal was detected in three of 48 controls with delayed onset. Differences across conditions were quantified by fitting binomial regressions to the first luminescence peak to estimate time to half-max luminescence (t50) values and by calculating mean luminescence normalized to absorbance for timepoints up to 12 hours after t50. These analyses confirmed OD 0.6 as the condition with the shortest t50 and highest mean signal, followed by OD 0.8 and OD 0.4, whereas other setpoints showed further delays or no evidence of evolution **(Fig. 3D), (Supp Fig. 20A)**. Together, these data indicate that the turbidostat OD used to seed lagoons can be tuned to accelerate measurable evolutionary progression.

The impact of lagoon flowrate on T7RNAP evolution was also evaluated by running evolutions at flowrates of 0.2 to 1.0 vols/hr. The lowest t50 values were observed at intermediate flowrates (0.4 to 0.6 vols/hr). Flowrates outside this range still produced luminescence increases consistent with evolution, but with moderately increased t50 values. Mean peak luminescence was broadly similar across conditions, with reduced signal at 1.0 vol/hr, consistent with delayed trajectories relative to the experiment duration **(Fig. 3D) (Supp Fig. 20B)**. Additionally, the effect of initial MOI on the rate of evolution was tested using a 10-fold dilution series spanning 10:1 to 1:10,000 phage-to-bacteria, evaluated at OD 0.6 under two lagoon flowrates (0.5 and 1.0 vols/hr) **(Fig. 3E) (Supp Fig. 20C)**. t50 values could not be estimated for MOIs below 1:1,000 due to the absence of an early luminescence transition within the experiment window, although late timepoint trends suggested delayed evolution. In contrast, higher MOIs were associated with faster luminescence onset and lower t50 values, most prominently at 0.5 vol/hr.

Finally, to decouple parameter effects on PACE operation from evolutionary outcomes, analogous experiments were performed using T7RNAP enrichments on pT7, without evolution **(Supp Fig. 21)** Consistent with prior reports, host cultures maintained at higher OD limited phage propagation in the T7RNAP-pT7 enrichment assay **(Supp Fig. 21A-B)**. Enrichments showed a sharp reduction in phage propagation capacity when turbidostat OD exceeded 0.6 **(Supp Fig. 21C,G-H)**. In contrast, higher lagoon flowrates produced substantially higher luminescence, consistent with increased phage propagation capacity under these conditions **(Supp Fig. 21C,G-H)**. Across MOI conditions, the fraction of luminescent lagoons decreased sharply with decreasing MOI and was absent at 1:10^7^ and below for both 0.5 and 1.0 vol/hr flowrates, indicating that a minimum infected host fraction is required to sustain infection and prevent phage washout, even with strong circuits **(Supp Fig. 21D,I-J)**. Together, these results demonstrate that TurboPRANCE enables rapid and parallel comparative PACE experiments that identify parameter regimes accelerating the accumulation of improved variants.

### Stability of turbidostats and staggered culture initiation

Complex PACE campaigns can run for more than 200 hours and are often executed as a series of discrete runs, each requiring its own turbidostat culture to be started, maintained, and then re-initiated for subsequent rounds, during which mutagenesis strains or plasmids can lose stability over time. To test whether TurboPRANCE could support staggered experiment starts from long-maintained cultures, T7RNAP-pT3 evolutions were initiated from staggered turbidostats that had been autonomously maintained on-deck for up to six days prior to starting evolutions **(Fig. 4A)**, using fresh MP6-transformants to initiate turbidostats approximately every 24 hours. Notably, growth rates within individual turbidostat cultures increased after 36-48 hr during continuous maintenance, consistent with progressive instability or corruption of the MP6 plasmid **(Supp Fig. 22A)**. MP6 is genetically unstable at high ODs, motivating the expectation that evolutionary kinetics would be delayed or absent in aged cultures. Despite this expectation, turbidostats aged six and seven days produced lagoons with luminescence increases on time scales comparable to younger turbidostats and exhibited higher mean luminescence **(Fig. 4B) (Supp Fig. 22B)**. These data indicate that turbidostats maintained for up to six days can support subsequent evolution, enabling delayed or staggered experiment initiation when required. For mutagenesis-plasmid sensitive hosts such as MP6, staggered operation also provides a route to periodically re-inoculate fresh turbidostats while maintaining ongoing lagoon experiments, enabling runs spanning days to weeks. In addition, because standard PACE campaigns are often executed as discrete ∼48 h runs and turbidostat sensor biofilming is a common operational constraint, this staggered re-inoculation strategy provides a practical mechanism to circumvent biofilm-driven drift during extended experiments. These results demonstrate that with TurboPRANCE turbidostats can be treated as modular inputs to the lagoon layer. When a culture has been maintained for an extended period or begins to drift in growth rate, it can be replaced with a freshly inoculated turbidostat without interrupting ongoing lagoon experiments or dismantling the system. This queueable exchange enables long-duration campaigns in which host strains are swapped in and out as needed, preserving lagoon continuity even when upstream cultures become unstable.

**Figure 4.**
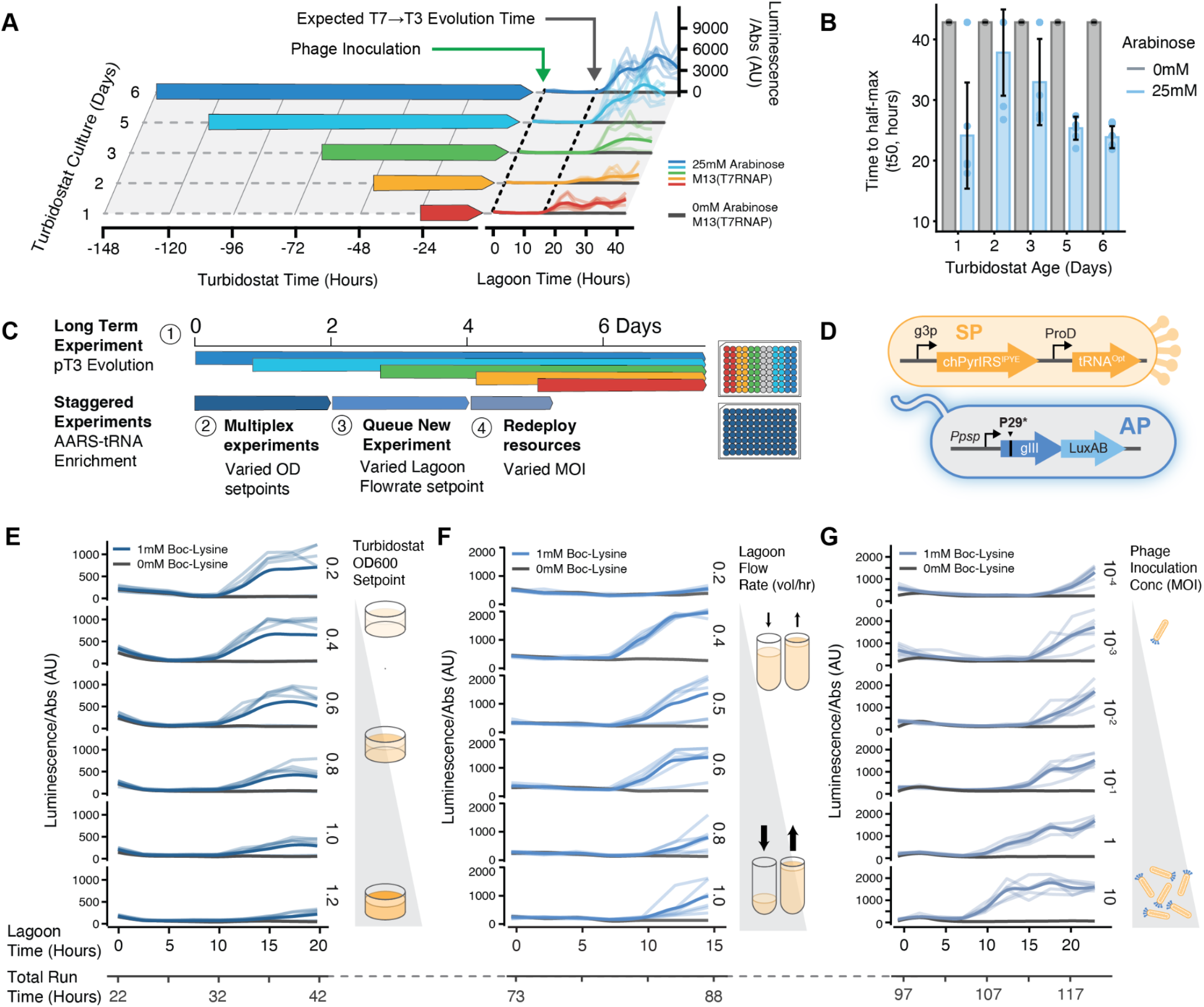
Parallelized and time-shared evolution campaigns. **(A)** T7RNAP pT3 evolution initiated from turbidostats aged 0 to 6 days prior to lagoon transfer. **(B)** Time to half-max luminescence (t50) across turbidostat ages under mutagenic (25 mM arabinose) and non-mutagenic (0 mM) conditions (2 controls and 6 evolutions per turbidostat age). **(C)** Long-term run architecture showing staggered turbidostat initiation and queued experiments. Independent parameter sweeps are multiplexed, new experiments are initiated while others are ongoing, and resources are redeployed without interrupting the run. **(D)** PACE Circuit for ncAA enrichments, coupling orthogonal tRNA synthetase and tRNA expression to a pIII–LuxAB accessory plasmid readout. **(E)** Turbidostat OD600 setpoint sweep (0.2 to 1.2) with lagoon luminescence and absorbance tracked over evolution time. **(F)** Lagoon flow-rate sweep (0.2 to 1.0 vol/hr) with lagoon luminescence and absorbance tracked over evolution time. **(G)** Initial phage inoculum (MOI) sweep with lagoon luminescence and absorbance tracked over evolution time. All ncAA experiments were conducted with 2 controls and 6 enrichments per condition. Error bars represent SD.

### Parallelized and time-shared evolution campaigns

To demonstrate that TurboPRANCE can support concurrent but distinct, and/or staggered experimental programs, three additional experiments were conducted during the same seven-day run **(Fig. 4C) (Supp Fig. 22C)**. To enable multi-experimental workflows, additional features were added to support multi-user operation and repeated experiment initiation within extended runs. Unique experiment identifiers were incorporated into metadata collection to support structured parsing and active run monitoring. Due to the observed accumulation of biofilm **(Supp Fig. 8)**, automated plate switching was also added to transfer turbidostats to a fresh 96-well plate without manual multichannel intervention. This capability permits queuing of new experiments by placing freshly inoculated turbidostats in designated wells of a new plate while omitting selected wells from the current plate during transfer, after which newly inoculated cultures are automatically maintained by the platform **(Supp Fig. 22D)**. Together, these developments supported the staggered initiation, parallel parameter exploration across distinct circuit architectures, and queueable multi-experiment operation within extended continuous runs.

To showcase this capability, phage were propagated using an orthogonal circuit architecture designed to evolve non-canonical amino acid (ncAA) incorporating orthogonal tRNAs and cognate tRNA synthetases **(Fig. 4D)**, with different media and supplement needs (i.e. addition of a non-canonical amino acid). Briefly, phage encoded a previously evolved tRNA-synthetase pair (chPylRS IPYE, tRNAopt^16,29^) that incorporates Boc-Lysine into amber codons, with pIII production gated by Boc-Lysine incorporation. Using this ncAA circuit, three parameter sweeps were carried out as independent experiments, performed in series: (i) a turbidostat OD setpoint optimization **(Fig. 4E)**, (ii) a lagoon flowrate sweep **(Fig. 4F)**, and (iii) an MOI scan **(Fig. 4F)**, as described above. These experiments were executed sequentially within the same continuous run with two different reset modes between experiments. Between the first (OD) and second (flowrate) experiments, turbidostats were fully re-inoculated alongside new lagoons. Between the second (flowrate) and third (MOI) experiments, the established turbidostats were retained and new lagoons were initiated, isolating lagoon-level parameters while reusing the same host cultures. Across the three ncAA experiments, results were consistent with trends observed in T7 RNAP enrichments. Lagoons supplied by high-OD turbidostats failed to propagate phage **(Fig. 4E)**, and an intermediate flowrate produced the strongest luminescence signal **(Fig. 4F)**. Even at lower MOIs, no washout was observed **(Fig. 4G)**, but signal was observed at substantially delayed rates compared to comparable T7RNAP selections **(Fig. 2F) (Supp Fig. 21D)**. This is consistent with the slower kinetics of aaRS driven UAG-pIII circuits compared to direct activation of the pT7 promoter by T7RNAP. Together, these data demonstrate rapid, iterative, and asynchronous experimentation on multiple circuit types within a single long run by either re-inoculating turbidostats between experiments or restarting lagoons while retaining established host populations.

Beyond running independent experiments in parallel, TurboPRANCE can also compose multiple selection pressures within a single lagoon by partitioning total inflow across independently maintained turbidostat sources within each dilution cycle. To validate this capability, T7RNAP-to-pT3 evolutions were performed using either one or two independent turbidostat sources at matched total dilution rates (0.5 vol/hr). Evolution trajectories were comparable between programs, and t50 values were not significantly different, confirming that per-cycle pressure composition changes are readily implementable **(Supp Fig. 23)**.

### High-throughput long-read sequencing evolution tracking

Building time-resolved trajectories of individual genotypes in PACE populations (including which mutations co-occur on the same molecule, and how specific haplotypes rise and fall over time) remains an outstanding practical challenge. Common approaches to track continuous evolution campaigns include Sanger sequencing of clonal plaques, short-read Illumina amplicon sequencing of mixed populations (for variants that fit within a paired-end read)^29^, or Illumina-based analysis coupled to ancestral reconstruction for longer variants^42^. Long-read Nanopore sequencing nominally sidesteps reconstruction by directly reading full-length molecules, but its per-read error rate is sufficiently high and structured that apparent read-level haplotypes cannot be interpreted as true genotypes^43^. Thus, to match the scale of TurboPRANCE, we developed a semi-automated Nanopore long-read workflow, paired with neural network trained variant calling^44^ as a downstream extension of the automation pipeline, enabling routine processing of evolution trajectories at throughput more commensurate with that of TurboPRANCE.

To validate this approach, terminal evolution samples were filtered to isolate phage from bacteria, amplified with multiplexed forward and reverse barcodes (up to 384 index pairs), and quantified in 384-well format for pooling. Following phage amplification and barcoding, initial attempts at quantification revealed that there was substantial background signal in the DNA quantification assay (Qubit^45^), despite the dsDNA-selective chemistry.. As a result, Qubit-based normalization did not reflect true differences in phage-derived amplicon abundance, and instead resulted in massive library pooling biases, further amplified by vastly different phage supernatant concentrations following evolution. Prior to optimization, DNA concentrations exhibited limited dispersion (CV = 0.30), while the resulting read fractions varied 6.3-fold more than DNA concentrations (F-test, p < 1e-49), indicating that normalization based on these measurements failed to control sequencing representation. Quantitative fidelity of phage library preparation was substantially improved through filtering phage supernatant and from adding a bead-based PCR cleanup step to remove residual primers, short fragments, and background absorbance from PCR mix chemical composition. Following purification, concentrations closely mirrored the distribution of maximum luminescence across the 96 lagoon evolutions, with comparable variability (CV = 1.16 vs. 1.26; F-test, p = 0.43) and a significant well-by-well correlation (Spearman rho = 0.64, p < 1e-11) **(Supp Fig. 24)**. These results confirm that quantification measurements accurately captured the skewed distribution of phage population sizes arising from differential evolution outcomes, enabling effective library normalization prior to pooling and sequencing. Samples were subsequently normalized per well using concentration-derived dilution factors and equimolar pooling was executed by a PyLabRobot^46^ script on an OT-2 **(Fig. 5A)** before being prepared using Nanopore ligation chemistry for Flongle sequencing.

**Figure 5.**
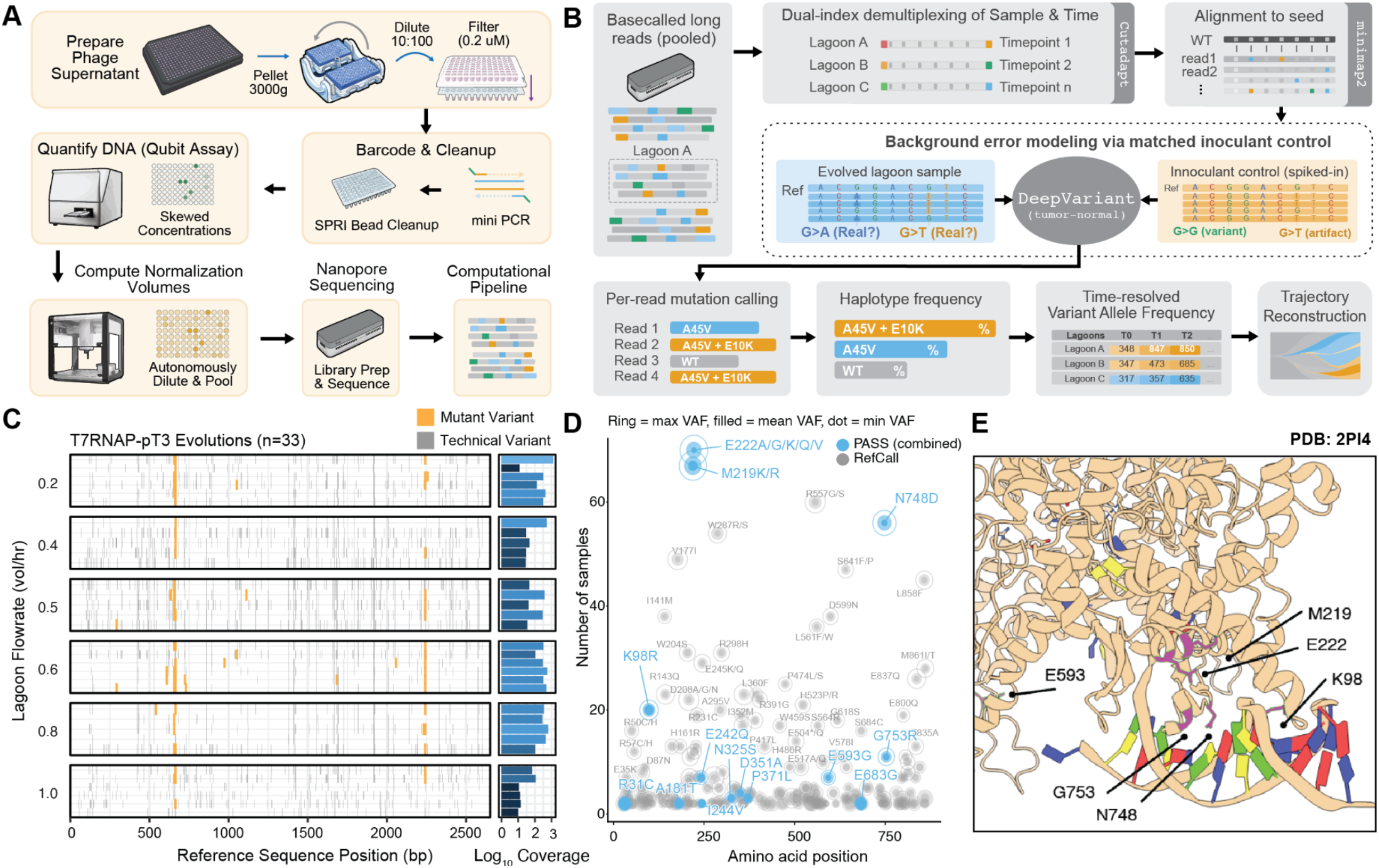
TurboPRANCE sequencing pipeline and variant landscape across evolution conditions. **(A)** Wet-lab workflow for preparing pooled nanopore sequencing libraries from PACE lagoon samples. Phage DNA is extracted, quantified, barcoded, and pooled at normalized concentrations for Oxford Nanopore sequencing. **(B)** Computational pipeline for extracting per-lagoon haplotype frequencies. Reads are demultiplexed by sample and timepoint (Cutadapt), aligned to the seed (Minimap2), and filtered for background sequencing errors using a tumor-normal-style approach: DeepVariant calls variants in both evolved samples and a spiked-in inoculant control, and shared sites are removed as systematic artifacts. Cleaned reads undergo per-read mutation calling to determine haplotype frequencies across lagoons and timepoints. **(C)** Per-read haplotype maps across 33 T7RNAP-pT3 evolution samples, faceted by lagoon flowrate. Each line is a sequenced read; amber = mutant variant, gray = technical variant (DeepSomatic). Right sidebar: per-sample coverage (log10). **(D)** Nonsynonymous variant landscape after rescue of missed mutations. Each point is a codon position detected in more than one sample; mutations at the same codon are combined. Open rings = max VAF, filled dots = min VAF. Blue = PASS (including rescued calls validated by at least one high-confidence PASS call), gray = remaining RefCall. **(E)** Crystal structure of T7RNAP in complex with pT7 and established and candidate evolved resides highlighted.

After sequencing, to identify the mutations associated with nanopore, reads were aligned to the expected parental sequence using Minimap2^47^ **(Fig. 5B)**. Nucleotide variants were then identified using DeepSomatic^48^, a deep neural network trained on noisy long-read data to enrich for true variants and flag technical artifacts. This is accomplished, in part, by comparing sequencing samples with suspected mutations with a “normal” control sample. Therefore, as a comparator with evolved samples when using DeepSomatic, the parental phage was deeply sequenced using the same library preparation strategy. DeepSomatic classifies candidate sites as PASS (supported variant), GERMLINE (technical artifact), or RefCall (no variant called). Using this variant calling strategy, mutations were assigned across all sequencing libraries to generate per-read haplotypes for 33 T7RNAP-pT3 evolution samples spanning a flow-rate series. This analysis captured both true biological variants and recurrent Nanopore-associated artifacts across the full amplicon **(Fig. 5C)**.

Artifact calls occurred at similar frequencies across all flow-rate conditions, consistent with a technical error profile rather than flow-rate dependent biology. This approach recovered canonical T7 RNAP adaptation mutations previously reported from Sanger sequencing of clonal phage in independent PACE campaigns^2,29^, despite the high amount of reproducible technical noise observed. Indeed, greater than 50 technical mutations occurred in > 10% of samples, with some exhibiting population level prevalence of >30% Variant Allele Frequency (VAF). Nonetheless, expected mutations M219R and N748D were detected with high VAF throughout in more than 60 evolutions, and mutations at E222 were also frequent (most commonly E222K, with lower-frequency E222A/G/Q/V), indicating that a DeepVariant-supported nanopore sequencing strategy can faithfully identify mutations in evolving populations with read-level resolution **(Fig. 5D)**. Additionally, variant maps were generated across all 83 sequenced evolutions, sorted by flow rate, MOI, and turbidostat age, which showed that these hotspots recur at consistent positions regardless of the evolution parameter **(Supp Fig. 25)**. Haplotype phasing was evaluated at the read level for residues M219, E222, and N748 across all evolution samples. Most populations were dominated by single substitutions at one of these sites (219, 222, or 748), rather than by multi-mutation haplotypes. However, a subset of samples contained detectable double mutants, including E222Q,N748D and E222K,N748D **(Supp Fig. 25D-E)**.

Across 77 pooled samples, 182 nonsynonymous PASS calls were identified, corresponding to 35 unique amino acid changes. Because calls are evaluated independently per sample, low-frequency variants in low-coverage samples can fall below the PASS threshold even when confidently detected elsewhere. Stratification by mapped read count confirmed this coverage dependence **(Supp Fig. 26A)**. Of the 35 PASS amino acid changes, 19 also appeared as GERMLINE or RefCall in other samples, representing 118 additional low-confidence observations in low coverage samples (i.e. in 0mM Arabinose controls). To address this, a targeted reassignment was applied: for any amino acid change classified as PASS in at least one sample (indicating confidence of absence from the parent lineages), the matching GERMLINE or RefCall observations in other samples were reclassified as “rescued” variants. This conservative strategy limits rescue to mutations supported high-confidence calls over background. Support for separation of real mutations from artifacts is further evident in recurrence patterns **(Supp Fig. 26C)**. Nonsynonymous changes never classified as PASS recurred broadly at similar low frequencies, consistent with systematic sequencing errors. In contrast, PASS-supported mutations showed heterogeneous recurrence across samples, consistent with biological variation accumulated during continuous evolution.

Using this strategy, lower-frequency variants were detected that, to our knowledge, have not yet been associated with T7RNAP-pT3 function. Several additional mutations stand out as low frequency relative to 219/222/748 but present across evolution conditions and experiments. This includes K98R, G753S, and E593G. Evaluating all mutant residues on an available structure of pT7-T7RNAP showed that, remarkably, while E593 was physically distant from the promoter interface both K98 and G753 are within the same distance, or closer, to the pT7 promoter as commonly mutated residues 219/222/748 **(Fig. 5E)**. These results suggest K98R and G753R as promising candidate mutations modifying pT3-T7RNAP activity. Showcased by the detection of both high-frequency mutation hotspots and low-frequency variants across a range of coverage depths, and resolution of haplotypes, Nanopore long-read sequencing can robustly resolve population-level full-length haplotypes in evolving phage populations when empowered by deep learning variant calling.

## Discussion

Highly multiplexed turbidostat maintenance, when coupled with single read sequencing resolution of evolving populations expands the conceptual scope of continuous evolution from single-objective optimization to systematic exploration of multidimensional fitness landscapes. A longstanding challenge in directed evolution is not simply identifying improved variants, but understanding how biomolecules navigate competing and shifting selection pressures to arrive at functional solutions. By enabling large-scale multiplexing across strains, selection circuits, and environmental variables, and by permitting reassignment or pooling of selective pressures over time, TurboPRANCE allows many dimensions of selection to be interrogated within a unified experimental framework. Rather than optimizing one property in isolation, the platform supports coordinated evolution under overlapping and dynamically structured constraints, more closely approximating the multi-objective nature of natural molecular evolution.

Leveraging horizontal gene transfer between many distinct bacterial circuits is a powerful, but still uncommon strategy in PACE. TurboPRANCE is designed to readily enable multi-strain evolution, allowing a population of phage to interface with dozens of distinct circuit strains and thereby support higher-dimensional selection architectures. Multi-strain configurations also create a straightforward route to combining positive selection with distributed negative selections. Prior PACE implementations of negative selection have typically co-localized positive and negative circuits within the same cell^37,38^. In contrast, distributing positive and negative pressures across strains could yield distinct outcomes by altering how variants experience and integrate selective constraints over time. Much like the immune system, where evolving repertoires are shaped by sequential sampling of different contexts^49,50^, staged exposure to separate pressures may favor different trajectories through sequence space and enrich epistatic intermediates that are difficult to access under a single, fixed configuration. This TurboPRANCE feature is particularly well suited to molecule classes where relevant objectives scale to the hundreds, including proteases^6,7^ tuned across substrate libraries, DNA-binding domains optimized across target and decoy sequences^12,21^, or protein-protein interaction modules engineered within large interaction families^11^. For example, an antibody could be evolved toward a desired target while selecting against dozens of off-target interactions in parallel, or an enzyme could be optimized across panels of substrates while simultaneously constraining unwanted side activities.

The system also enables a single biomolecule to be evolved towards multiple independent biomolecular properties. For example, a single population could be selected simultaneously for improved solubility^10^, catalytic efficiency, reduced off-target activity, or folding, with each property encoded in a different selection strain. Beyond the circuit itself, strains could be further engineered to modulate how variants experience selection, for example by enabling selective transport or efflux of small molecules, tuning intracellular cofactor availability, or producing (or withholding) specific substrates and intermediates, thereby expanding the range of chemically and metabolically defined environments that can be imposed through genetics rather than manual supplementation.

From an engineering perspective, this platform is fundamentally designed so that robotic complexity does not translate into user burden. In contrast to PRANCE^29^ (which required constant off-deck bacteria preparation), initiating a TurboPRANCE experiment requires only inoculating a single colony into a designated well of a 96-well plate, after which equilibration, density control, and lagoon transfer proceed autonomously. This substantially reduces experimental burden and eliminates the need for constant culture babysitting during multi-day campaigns. As such, the platform can run continuously for several days with minimal intervention, typically requiring only periodic refills of media and wash fluids in bulk storage vessels, and collecting sample plates. Users can also time-share and queue non-overlapping experiments asynchronously while other evolutions are underway by partitioning turbidostat plates into independent user spaces, conceptually analogous to time-sharing operating systems such as MULTICS^51^. TurboPRANCE seamlessly tracks metadata and exposes fine-grained experimental control through a programmable manifest **(Supp Fig. 5)** that defines turbidostat-to-lagoon mappings and transfer fractions, enabling manual or algorithmic target reassignment and real-time stringency modulation without interrupting operation. Given prior results showing that stringency schedules can strongly influence evolutionary success^29^, this flexibility should enable systematic discovery of effective stringency programs and target assignment patterns that elicit distinct trajectories through sequence space.

A practical limitation encountered during extended TurboPRANCE operation is the intrinsic instability of mutagenesis strains (such as MP6^4^). Though evolution was observed in our T7RNAP variants with aged turbidostats (e.g. likely destabilized MP6), the original T7RNAP to TP3 evolution is a fairly accessible target, as it was originally obtained using weak mutagenesis^2,4^, suggesting that strongly beneficial mutations can be recovered under relatively modest mutational pressure. More restrictive evolutionary targets may not exhibit similar tolerance to extended pre-evolution maintenance. Nonetheless, automated plate switching can recover turbidostats that drift from their setpoint, but mutagenesis plasmid loss introduces additional variability: as the burden decays, host growth accelerates and faster-growing cells can rapidly dominate a turbidostat, complicating long-duration runs. A staggered architecture provides a practical workaround by initiating lagoons from temporally offset turbidostats, allowing fresh mutagenesis-competent cultures to enter while ongoing lagoons continue uninterrupted. Staggering requires queueing replacement cultures and therefore trades capacity for robustness, but the reduction is bounded, at most two-fold in the limiting case of one queued replacement per active turbidostat, and is primarily relevant for campaigns exceeding 4 days. Prior robotics-enabled evolution workflows typically required frequent culture preparation and hands-on intervention to sustain multi-day operation^29^. Further, classic turbidostats are commonly disrupted on approximately 48 hour timescales when biofilms interfere with turbidity sensing and trigger erroneous overdilution. In this context, queueable, staggered turbidostat initiation provides a substantially lower-burden route to sustained operation and may represent one of the most operationally sophisticated demonstrations of continuous MP6 host maintenance to date. Nonetheless, mutagenesis plasmid variability remains a broader constraint of continuous evolution systems, and molecular engineering to stabilize mutagenesis strains would improve long-term robustness.

Although deploying TurboPRANCE requires mechanical assembly, fluidics setup, and software installation, replication has been straightforward in practice and validated across multiple hardware builds. Additionally, protocols, a bill of materials, and documentation for configuring fluidics and other hardware will be released alongside the full source code upon final publication to standardize setup and reduce site-to-site ambiguity. The ease of adoption is purposefully built into the method, as TurboPRANCE separates method logic from hardware-specific parameters through local configuration files, in order to accommodate alternative equipment and site-specific calibrations without modifying core code. More broadly, as laboratory automation continues to diversify rapidly, the modular software interfaces used here are designed to carry TurboPRANCE beyond a single instrument, enabling porting to additional platforms via PyLabRobot^46^ and to emerging open, purpose-built robotic systems^52^. With minor modifications, the platform can also be paired with lower-footprint, lower-cost plate readers as recently described. Nonetheless, the current implementation runs on a Hamilton STARlet, a mechanically sophisticated, relatively low-footprint workhorse that remains an industry gold standard and is widely available— including in many core facilities and as refurbished, routinely serviceable systems.

Ultimately, TurboPRANCE reframes throughput in directed evolution as a means of fine-grained sampling of the fitness landscape surface rather than simply increasing scale. Obtaining this structure permits comparative analysis across unrelated selective pressures and supports experiments in which pressures are alternated, combined, or temporally staged within the same evolving population. The resulting trajectories are not single endpoints but time-resolved records of adaptation across distinct conditions. The advantages of the platform are highly complementary to limitations of generative protein design pipelines. Existing AI models for protein design are primarily trained with supervised learning on labeled datasets of existing sequences annotated with folding structure or other physical properties^53–56^, entailing challenges of data sparsity and bias. TurboPRANCE overcomes these limitations in a manner similar to reinforcement learning with a dense reward function, due to its capacity for self-guided exploration of a tunable fitness landscape. As AI-guided design continues to increase in capabilities, the principal limitations will become the dimensionality and resolution of available datasets. In the future, multivariate evolutionary time series that report how mutations jointly arise across distinct pressures will represent a fundamentally different class of training data than static screens. In this view, the platform presented herein is not only for evolving improved molecules, but a framework for generating structured, multi-objective datasets that enable quantitative interrogation of how selective pressures shape sequence space and ultimately function.

## Acknowledgements

This work was supported through generous funding to the Chory Lab by Duke University as well as an award from the Hypothesis Fund to E.J.C. Additionally, work was made possible with support through the National Institute of General Medical Sciences including the Ruth L. Kirschstein NRSA fellowship awarded to R.M.B. (F32-GM157893) as well as funds from the Biomolecular and Tissue Engineering Training grant awarded to S.M.G. (T32-GM144291).

## Availability of Data and Reagents

Raw and processed sequencing data generated during this study will be deposited GEO upon publication. A bill of materials and links to 3D part files will be provided upon final publication. Any phage and plasmids generated and data not included in the final version of this study are available by reasonable request. For status updates or early access to in-progress documentation, please contact.

## Ethics Declarations

E.J.C., S.M.G., and R.M.B. are listed as co-inventors on a pending patent application for TurboPRANCE. E.J.C. is a co-founder of and holds equity in Bullseye Biosciences, Inc. S.M.G. is a consultant and also holds equity in Bullseye Biosciences, In

## Supplemental Figures

**Supplemental Figure 1.**
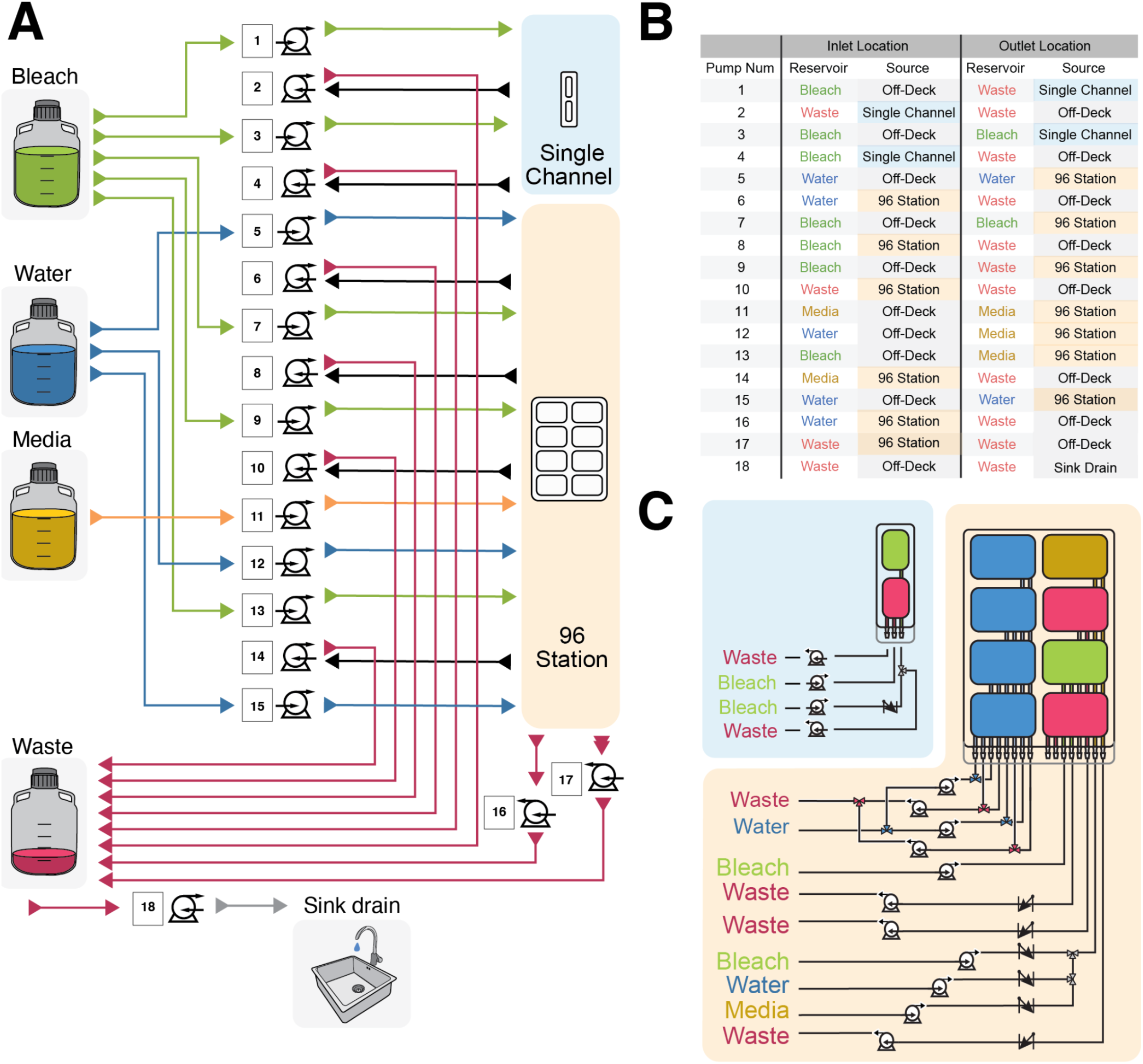
Fluidics and pump architecture. **(A)** Schematic of the peristaltic pump network linking off-deck bleach, water, media, and waste reservoirs to on-deck components. Pumps 1-18 drive directed flow to the single-channel washer, 96-channel washer station, media staging basin, and waste lines. Green lines indicate bleach, blue indicate water, gold indicate media, and red indicate waste. The sink drain provides final off-deck disposal. **(B)** Pump routing table defining inlet and outlet assignments for each pump channel. For each pump number, the source reservoir and destination reservoir are specified, enabling programmable routing between off-deck carboys, on-deck washer reservoirs, media stations, and waste containers. **(C)** On-deck washer modules and internal channelization. The single-channel washer (top) and 96-channel station (bottom) contain segregated bleach, water, media, and waste compartments with dedicated inlet and outlet lines. Independent pump control allows asynchronous refilling, draining, and bleach dosing during operation while maintaining fluid separation and minimizing cross-contamination.

**Supplemental Figure 2.**
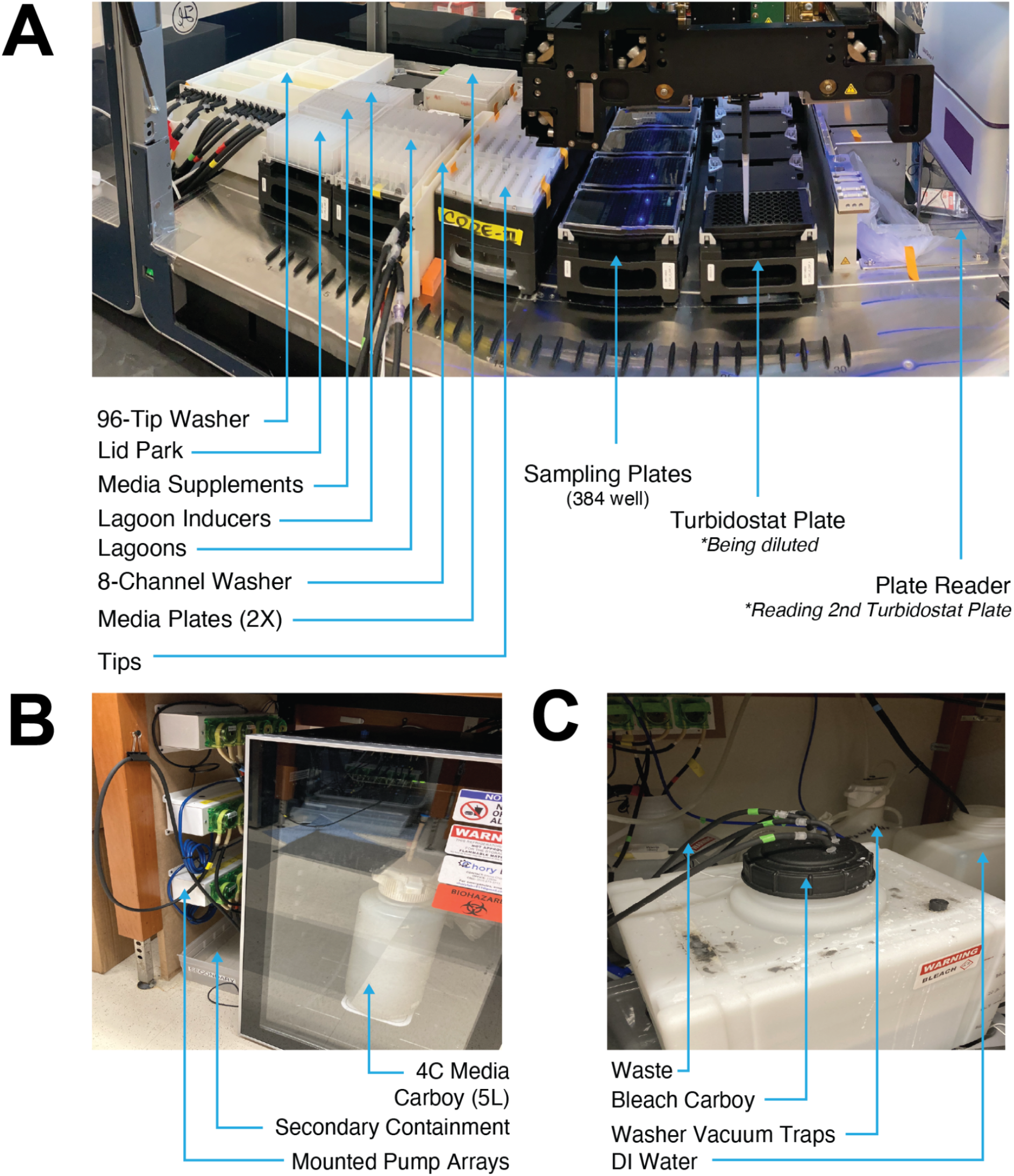
Physical TurboPRANCE platform and fluidic infrastructure. **(A) On-deck layout of the Hamilton STARlet-based TurboPRANCE system**. Shown are the 96-tip washer, lid park, media supplements, lagoon inducers, lagoon plate, 8-channel washer, dual media staging plates, tip racks, 384-well sampling plates, active turbidostat plate (being diluted), and plate reader (reading the alternate turbidostat plate). The layout enables alternating plate read and dilution steps while maintaining physical separation of media, lagoons, and wash stations to reduce cross-contamination. **(B) Under-bench media delivery system**. A 4 C media carboy (5 L) supplies fresh media to the deck via mounted peristaltic pump arrays. Secondary containment surrounds fluid reservoirs and pump hardware for spill protection. **(C) Waste and wash infrastructure**. Separate carboys for waste, bleach, and DI water feed the washer modules and collect effluent. Vacuum traps and dedicated tubing isolate bleach, water, and waste streams, supporting automated refilling, draining, and sterilization during multi-day operation.

**Supplemental Figure 3.**
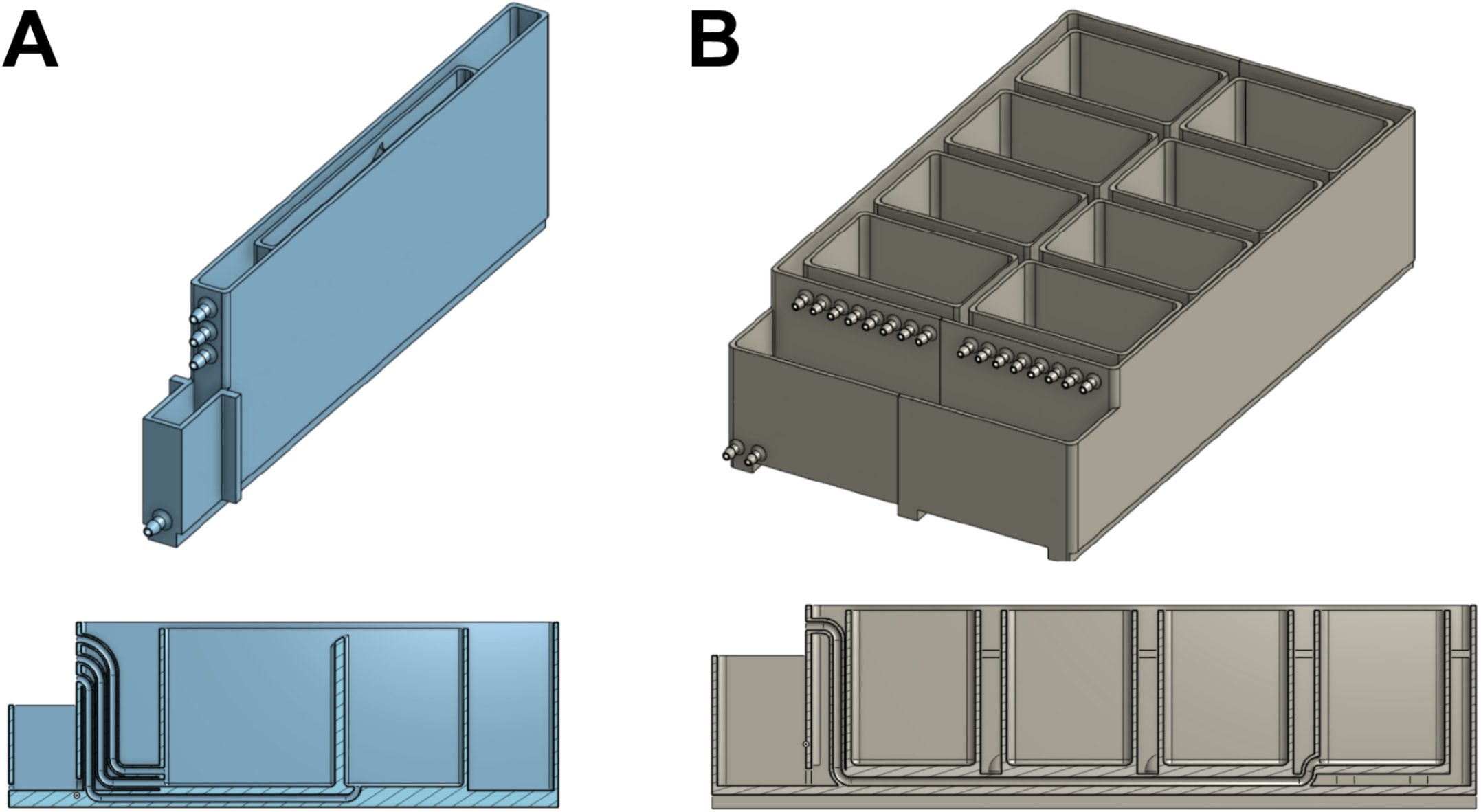
(A) Single-channel washer. (B) Eight-reservoir washer. Both washers are monolithic components 3D printed by SLA and incorporate multiple fluid reservoirs with dedicated inlets and outlets for bleach, water, media, and waste handling. Internal channels (1 mm wall thickness) route each reservoir to standard 3/16 in barbed fittings for flexible tubing. Each washer includes built-in secondary containment that captures reservoir overflow and directs it to a dedicated waste outlet; all secondary containment basins are also equipped with a conductivity-triggered overflow sensor that activates a backup sump pump to drain directly to the sink via an independent outlet in the event of overflow or pump failure.

**Supplemental Figure 4.**
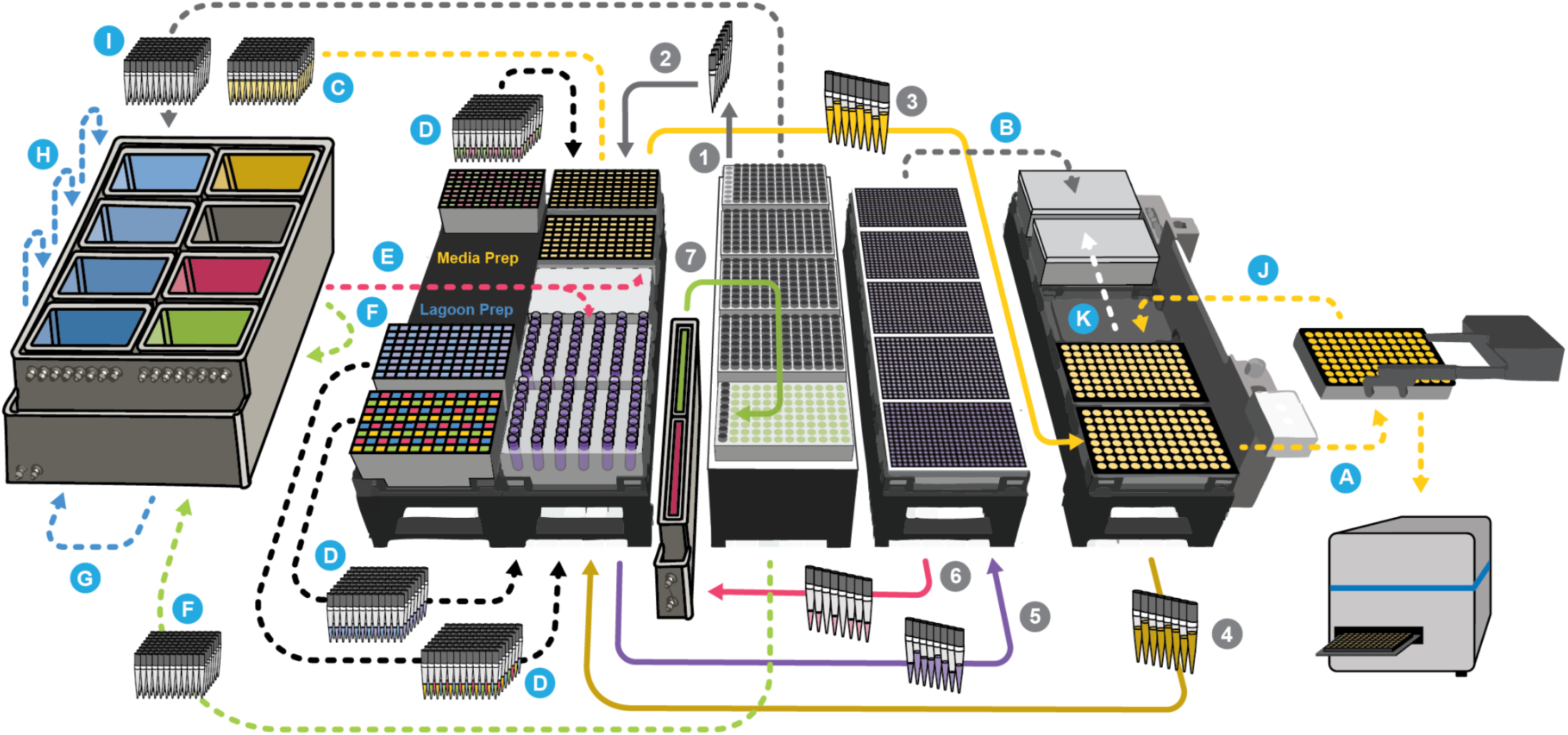
TurboPRANCE full method overview with two asynchronous processes. Gray numbers (1-7) indicate the repeating per-cycle actions executed in order: (1) move the active turbidostat plate to the reader and acquire OD600 (and reporter, if used), (2) plate and lid moves to enable alternating read vs service, (3) dispense fresh media to turbidostat wells, (4) aspirate excess culture to hold constant volume and generate outflow, (5) route defined outflow volumes to mapped lagoons, (6) sample lagoons into 384-well plates for measurement, and (7) update control outputs (replacement volumes and projected Δ t) for the next cycle. Blue letters (A-K) denote operations that run asynchronously, in parallel with the numbered cycle when possible: (E) on-deck media formulation/refill, (F) lagoon reagent prep and staging, (C,D) tip handling and sterilization workflows (clean, dirty staging, reuse), (G,H) washer and waste-line maintenance (drain, refill, bleach dosing), and (A,B,J,K) sample-plate handling and plate-reader workflows (staging, reads, and returns) coordinated with deck plate/lid logistics.

**Supplemental Figure 5.**
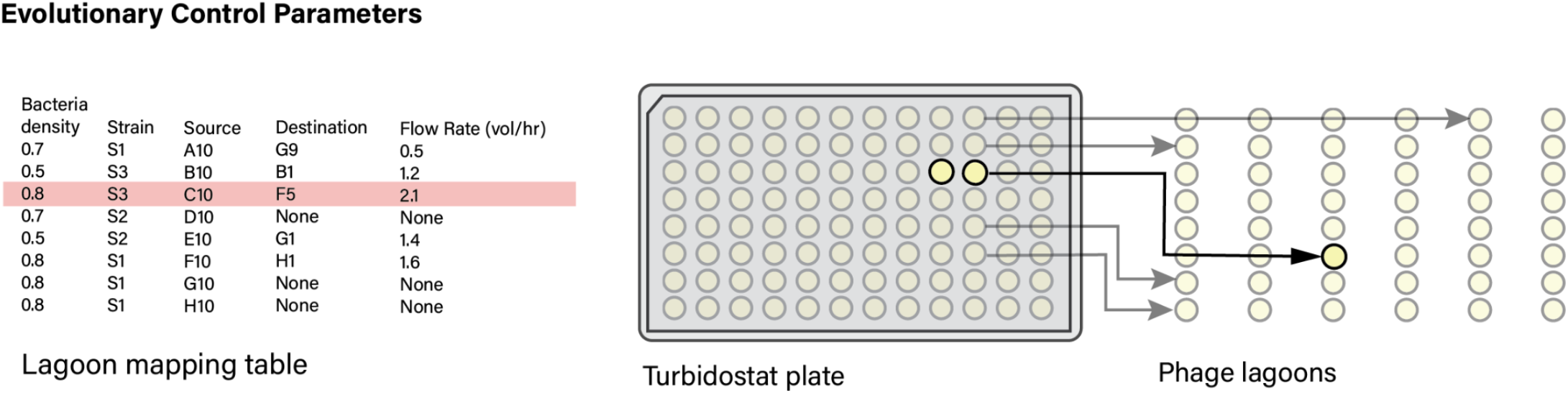
Turbidostat-to-lagoon mapping via manifest-defined routing. **Left:** Example lagoon mapping table specifying per-well control parameters, including OD setpoint (bacteria density), strain identity, turbidostat source well, lagoon destination well, and target flow rate (volumes per hour). Each row defines one turbidostat well and its assigned lagoon behavior. **Right:** Schematic of routing logic. Turbidostat wells on the plate are dynamically mapped to specific phage lagoons based on the manifest. During each cycle, defined fractions of excess culture are transferred from source wells to their designated lagoon wells at the specified flow rates. Wells with “None” as destination are maintained as standalone turbidostats. In batched column configurations, controller parameters are applied per batch, and pooled outflow from grouped wells is routed to the same lagoon, allowing flexible reassignment of sources, destinations, and stringency without modifying the underlying control code.

**Supplemental Figure 6.**
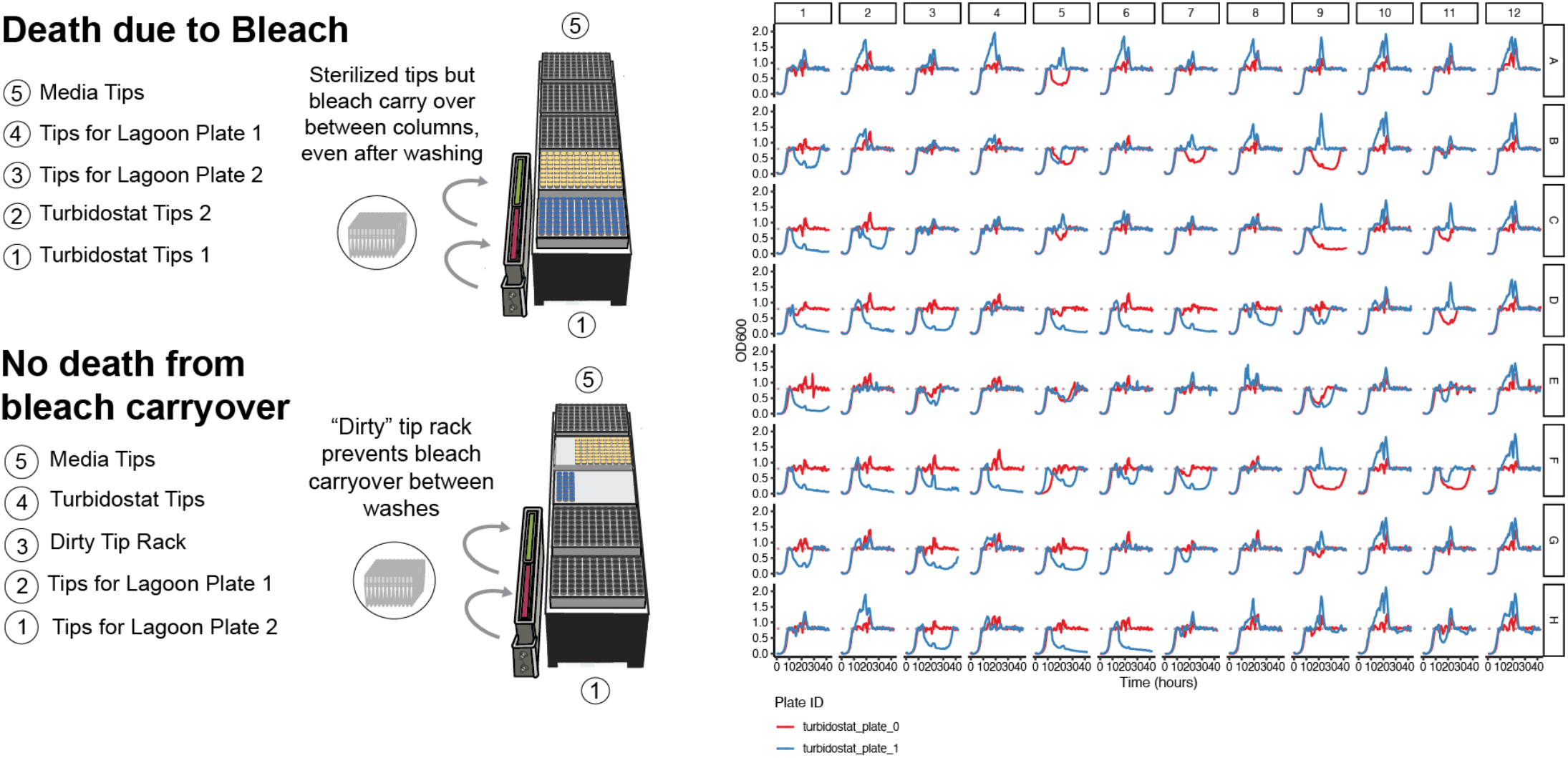
Bleach carryover and implementation of a segregated dirty tip workflow. **Top:** Stochastic turbidostat death caused by bleach carryover. When large-volume tips (p1000) are repeatedly bleached and ejected, increased ejection force is required over time to prevent tip sticking and cling to pipette channels. This mechanical force generates residual splatter that coats tips, the tip rack, and the tip rack housing which allows trace bleach to transfer between columns of tips after the tip cleaning process and intermittently kill bacteria. Representative OD600 traces (right) show sudden culture collapse consistent with bleach exposure. Spikes in measured OD are also often associated with opaque speckles in exposed wells due to the conglomeration of dead bacteria. **Bottom:** Dirty tip rack strategy eliminates bleach carryover. To prevent bleach contamination, all recently bleached tips are first transferred to a designated “dirty” tip rack that never contacts clean staging areas. The 96-head subsequently re-picks these tips, fully sterilizes them including water rinses after bleaching, and returns them to their original housing rack. This segregation prevents bleach splashback from entering clean tip positions while preserving intermittent bleaching cycles. Considering intermittent bleaching is essential for eliminating both turbidostat back-contamination and phage cross-contamination; thus, physical segregation of post-bleach tips from clean storage was required to maintain sterility without inducing culture death.

**Supplemental Figure 7.**
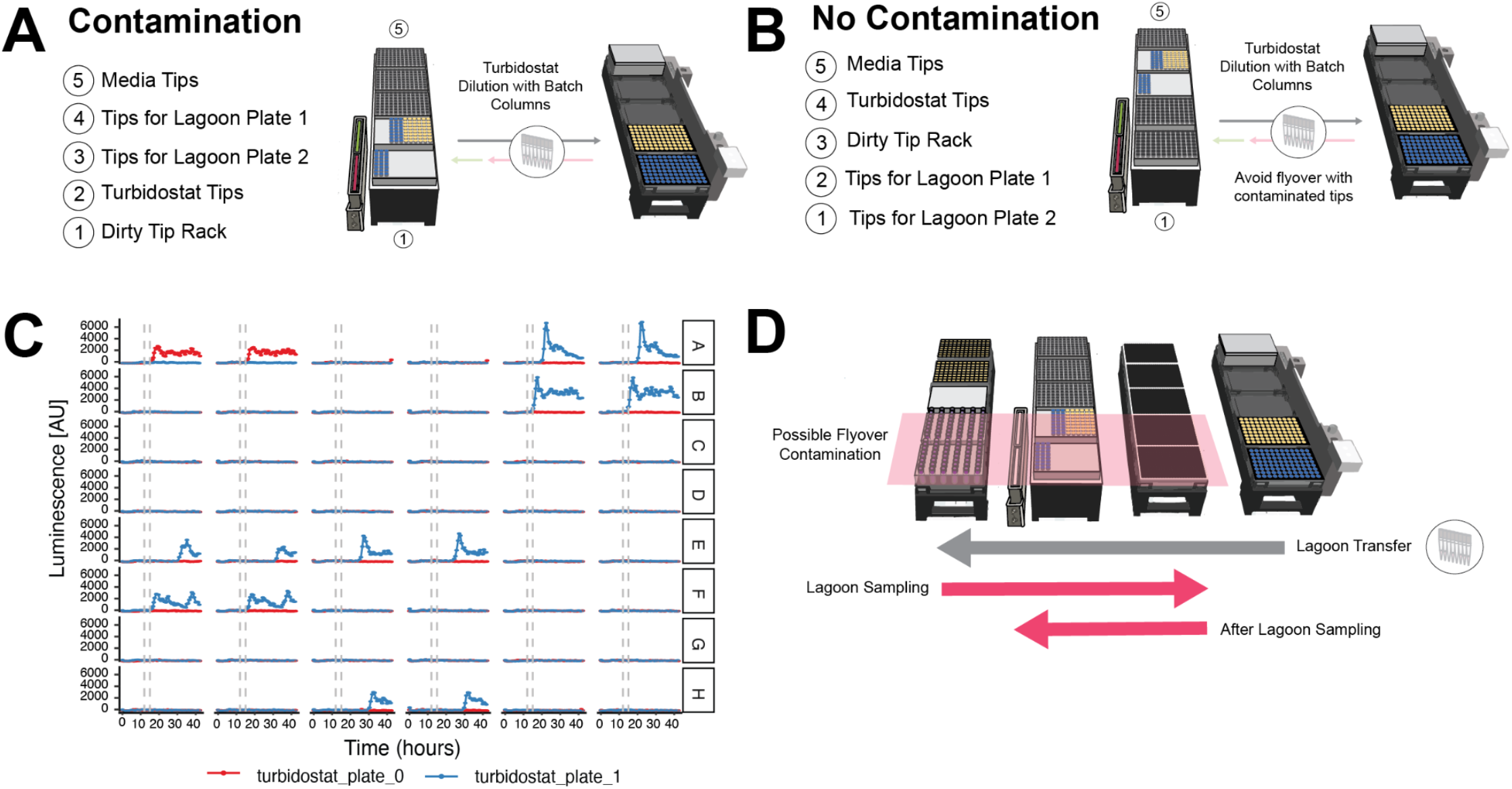
Eliminating turbidostat cross-contamination from lagoon-associated flyover. **(A)** Contamination-prone deck configuration. Clean tip racks positioned near lagoon sampling operations are exposed to potential droplet flyover from phage-contaminated tips during lagoon transfer and sampling. In this layout, tips designated for turbidostat dilution can be secondarily contaminated. **(B)** Revised deck configuration preventing contamination. Clean turbidostat tips are repositioned away from lagoon sampling paths, and a segregated dirty tip rack is introduced. This layout minimizes flyover exposure of clean tips while maintaining batching-based turbidostat dilution. **(C)** Representative turbidostat luminescence traces demonstrating back-contamination originally from lagoons. Wells from two independent turbidostat plates (red and blue) show delayed luminescence increases following lagoon inoculation (gray dashed lines), consistent with unintended phage transfer back into source turbidostats. After tip rack relocation, sporadic gains in luminescence, indicative of infection, were no longer observed across subsequent experiments. **(D)** Schematic of possible flyover contamination during lagoon sampling and transfer. During lagoon sampling, phage-contaminated tips traverse the deck; if clean tip racks are positioned within this path, aerosolized or splashed droplets can deposit phage either from flyover or from proximity to the single channel washer unit when phage contaminated media waste is expelled. Repositioning clean tips outside this corridor eliminated cross-contamination while preserving lagoon transfer functionality.

**Supplemental Figure 8.**
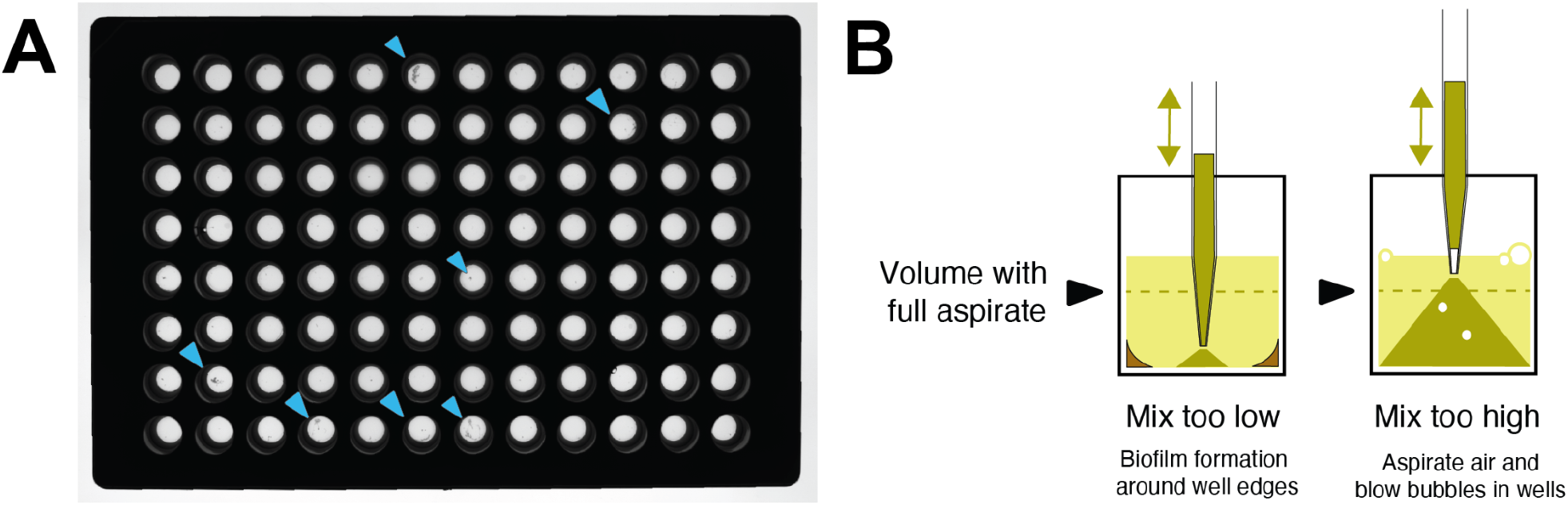
Biofilm emergence in turbidostat wells and mixing-height tradeoffs. **(A)** Representative turbidostat plate showing well-to-well biofilm accumulation after extended operation. Biofilm frequently appears in repeated patterns across batched wells (examples marked by blue arrows), consistent with transfer and nucleation during grouped processing. **(B)** Mixing-height constraints during automated dilutions. Mixing too low leaves material near the well perimeter and promotes edge-associated biofilm, while mixing too high increases air aspiration and bubble formation, degrading OD600 readouts and culture handling. To reduce biofilm in TurboPRANCE, turbidostat dilutions include repeated aspiration/dispense mixing steps and plates are orbitally shaken in the plate reader prior to measurement. Biofilm can still accumulate over 16-24 hours, so soluble culture is periodically transferred to a fresh 96-well plate (and severe wells can be manually cleaned). Mixing parameters were tuned to balance mechanical disruption of biofilm against excessive shear that can reduce F-pilus integrity and phage infectability.

**Supplemental Figure 9.**
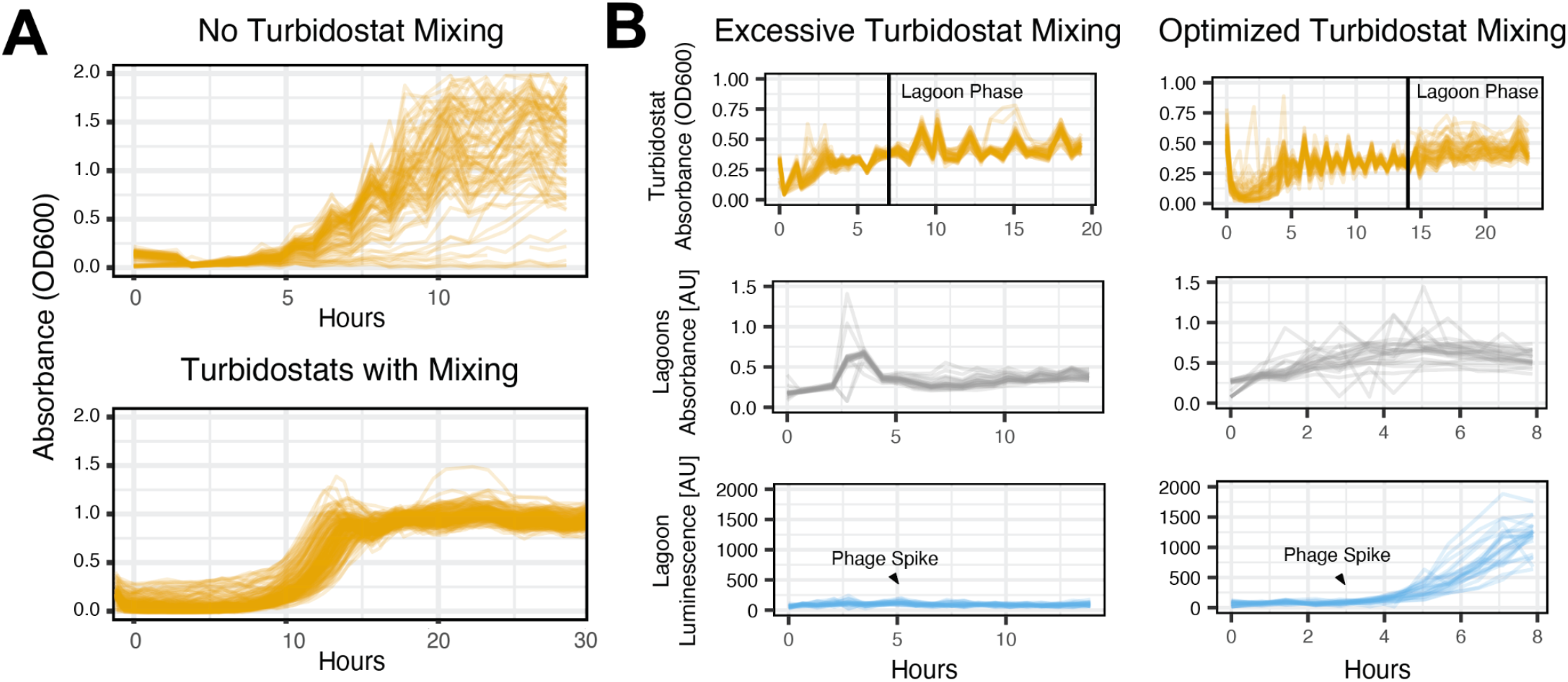
Mixing is required for stable OD control but must be tuned to preserve phage infectivity. **(A)** Turbidostat OD600 trajectories with and without pipette mixing. Without mixing, wells show large apparent OD600 noise and divergence consistent with spatial heterogeneity and biofilm; adding periodic mixing yields smoother, more stable OD control across wells. **(B)** Tradeoff between mixing aggressiveness and downstream phage performance. Top row: turbidostat absorbance during equilibration and after transition into lagoon phase (vertical line). Middle row: lagoon absorbance. Bottom row: lagoon luminescence after a defined phage spike (arrow). Excessive turbidostat mixing and/or high aspiration speed during transfers suppresses phage activity (likely due to F-pillus shearing) whereas optimized mixing (less frequent mixing with gentler aspiration) maintains turbidostat stability while enabling robust infection and signal amplification in lagoons.

**Supplemental Figure 10.**
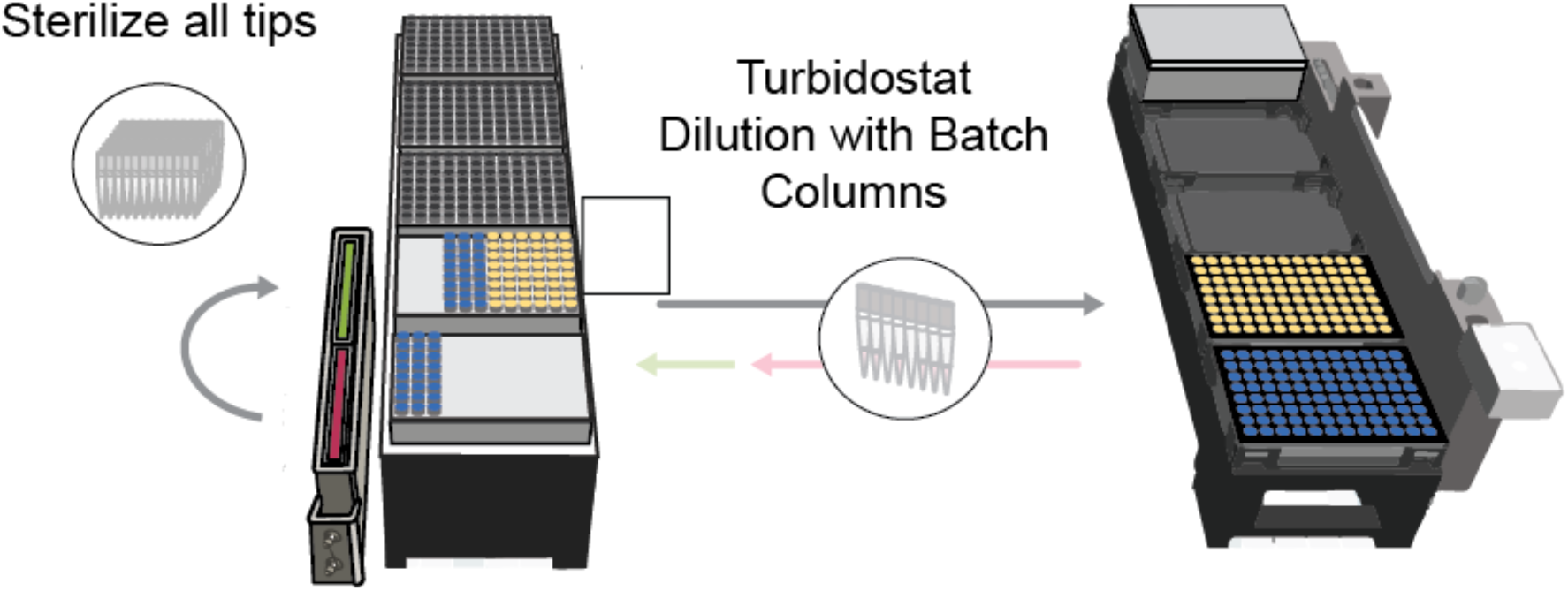
Column batching to increase available host outflow for slow-growing turbidostats. Half a rack of clean tips is capable of servicing each turbidostat plate in the case where two columns of a turbidostat plate are batched diluted together in the same step. TurboPRANCE can batch multiple turbidostat source wells per condition by splitting a single strain across adjacent turbidostat columns and servicing them as one unit during a dilution cycle. In this mode, tips are sterilized and reused, then a single media aspiration is used to dilute the full batch, and excess culture from the batched wells is pooled and routed to the same lagoon destination. Pooling increases the total available volume of bacteria per cycle, enabling higher lagoon flow-through rates even when individual cultures grow too slowly to sustain the desired demand from a single well. Batching is configured by changing only the batch size (number of columns per condition), without changing control logic or per-well setpoints. The tradeoff is reduced strain capacity per plate (fewer distinct conditions), but batching enables robust operation for burdensome or slow-growing strains when high continuous outflow is required.

**Supplemental Figure 11.**
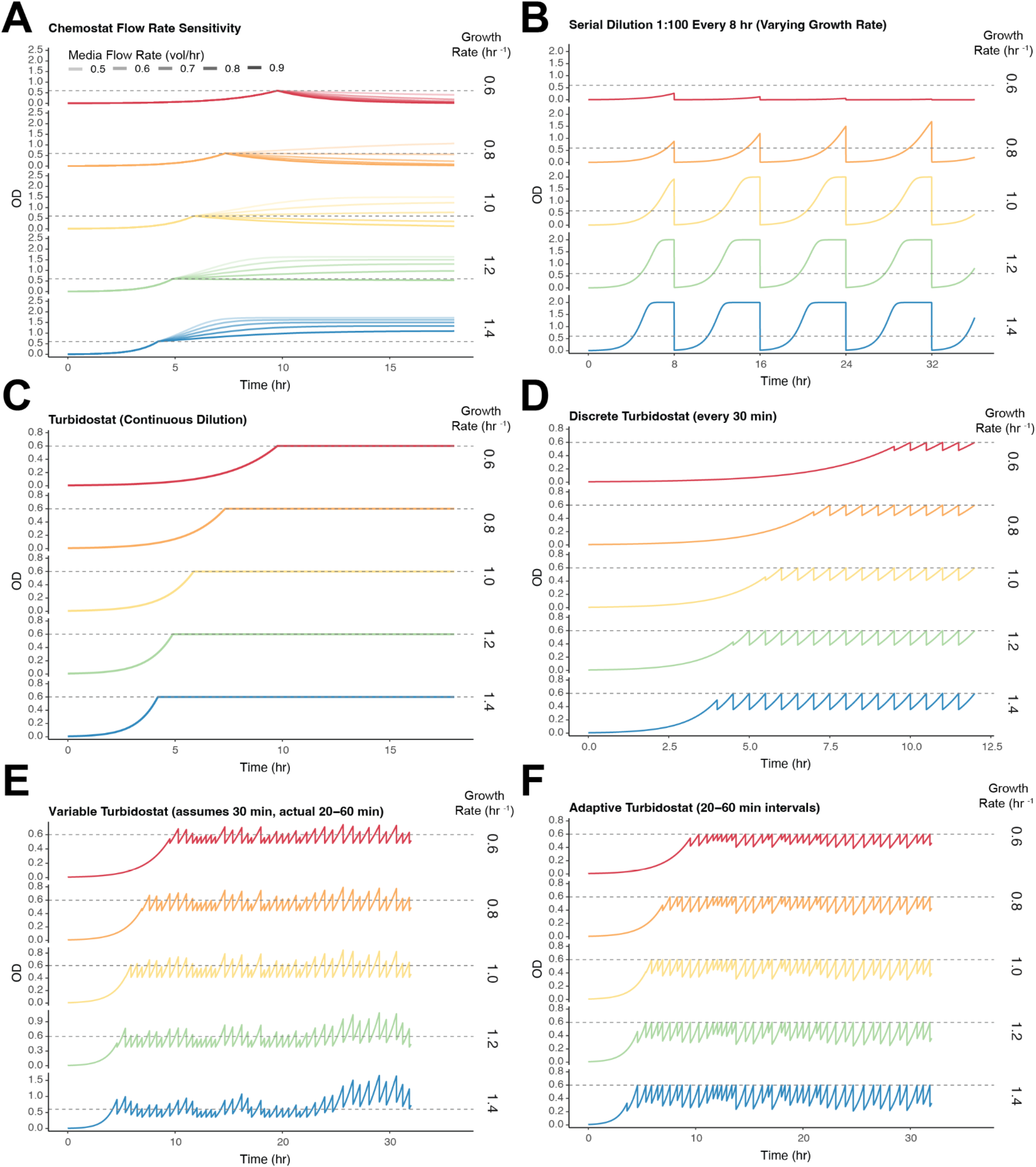
Simulated growth dynamics under six culture control strategies. All panels show optical density (OD) over time using Monod kinetics with a target setpoint of OD 0.6 (dashed line). Rows within each panel correspond to bacterial growth rates of 0.6, 0.8, 1.0, 1.2, 1.4 hr^-1^. **(A)** Chemostat at constant dilution rates of 0.5 - 0.9 vol/hr (light to dark lines). **(B)** Serial dilution (1:100 every 8 hr). **(C)** Continuous turbidostat. **(D)** Discrete turbidostat with fixed 30-minute measurement intervals. **(E)** Non-adaptive turbidostat using random 20-60 minute measurement intervals but assuming a fixed 30-minute interval for dilution calculations, with a maximum two-fold dilution constraint. **(F)** Adaptive turbidostat with random 20-60 minute intervals, where the system uses knowledge of growth rate and actual upcoming interval to calculate precise dilution depth.

**Supplemental Figure 12.**
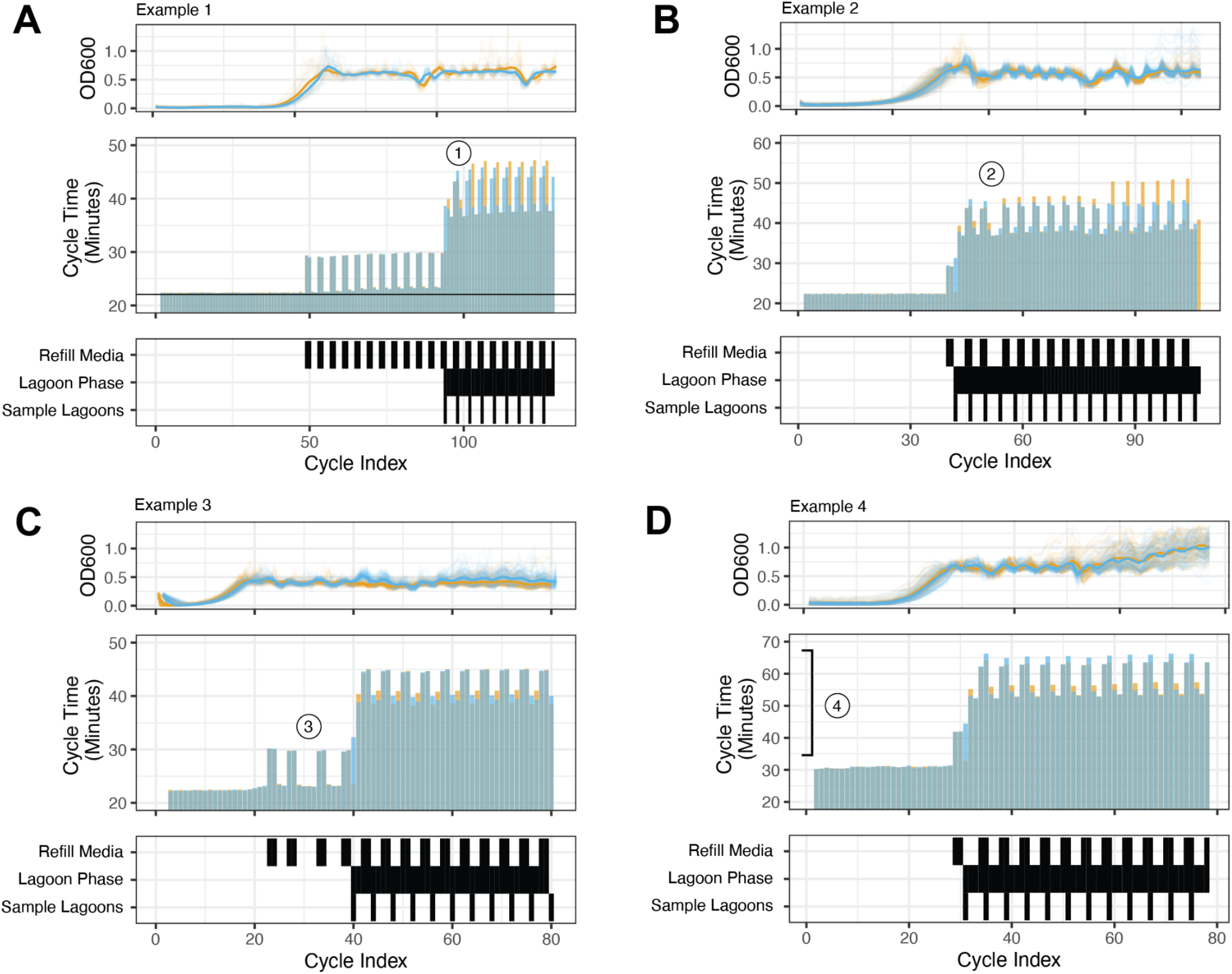
Cycle-time unpredictability across experiments with variable operational load. Each panel shows OD600 trajectories (top), measured cycle times (middle), and the timing of refill, lagoon phase, and sampling events (bottom) for representative runs. **(A)** Turbidostat plates are not necessarily synchronized. Even under similar starting conditions, the two plates diverge in cycle duration and must be projected independently rather than assuming a shared fixed Δt. **(B)** Media refill events occur irregularly and are influenced by culture number and growth rate. As active wells increase and growth accelerates, refill frequency shifts, introducing non-periodic extensions to cycle time. **(C)** Refill timing can change abruptly during a run, including before lagoon activation. Transitions in workload alter cycle duration independent of lagoon phase status. **(D)** Hardware-imposed limits constrain achievable cycle times, as shown here from a TurboPRANCE run using a slower liquid handler than in A,B,C. As lagoon phase begins and demand increases, cycle durations step upward and plateau at values dictated by pipetting throughput, defining the maximum growth rates that can be supported without loss of control. Together, these examples illustrate that cycle duration is an emergent property of biological growth and robotic workload, and cannot be reliably predicted using static or globally fixed timing assumptions.

**Supplemental Figure 13.**
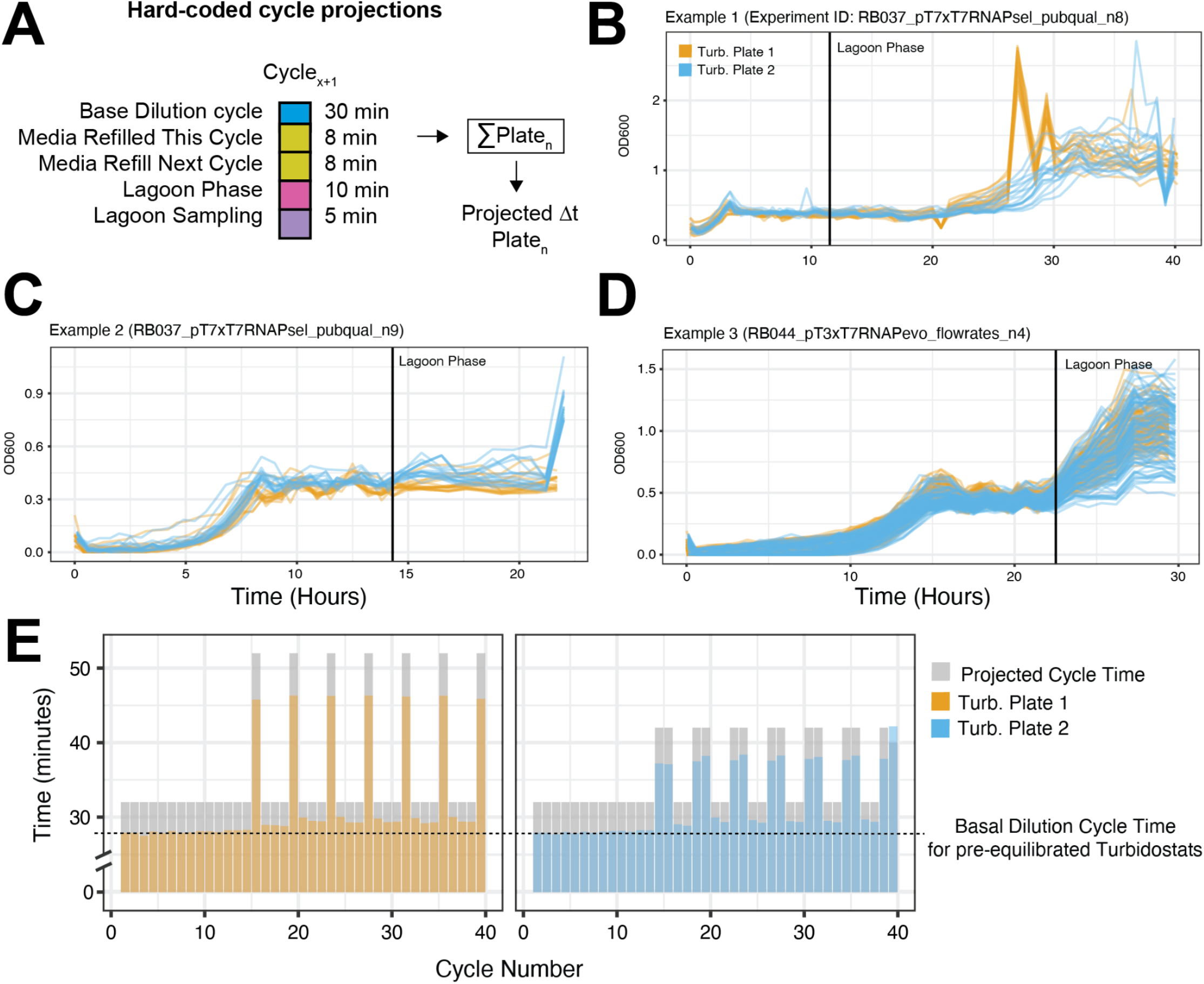
Hard-coded cycle time projections fail under variable operational demand. **(A)** Schematic of hard-coded cycle projection. Total projected Δ t for the next cycle is calculated as the sum of fixed contributions (base dilution time plus optional media refill, lagoon phase, and sampling increments). This assumes deterministic timing across cycles. **(B-D)** Representative experiments demonstrating breakdown of fixed projections. Vertical line marks lagoon phase initiation. Across independent runs, OD600 trajectories diverge when operational load changes, particularly after lagoon activation, indicating that fixed timing assumptions cannot accommodate dynamic pipetting volumes, refill events, and sampling operations. Observations akin to examples B and C often occurred in hard-coded projection schemes when the periodicity of media refill and sampling change in the middle of an experiment. Gradual loss of turbidostat maintenance as seen in D is commonly seen when turbidostats fail to adequately dilute one or more cycles and a positive feedback loop occurs where turbidostats continue to be underdiluted as hard-coded projections are unable to account for extra time needed when pipetting larger volumes. **(E)** Measured cycle times (colored bars) compared to projected times (gray). Actual cycle durations vary substantially across cycles and between plates relative to the nominal basal dilution time (dashed line). Media refills, lagoon sampling, and variable dilution volumes introduce irregular timing shifts and can synergistically add cycle time that is not captured by static projections, motivating the need for adaptive cycle-time estimation.

**Supplemental Figure 14.**
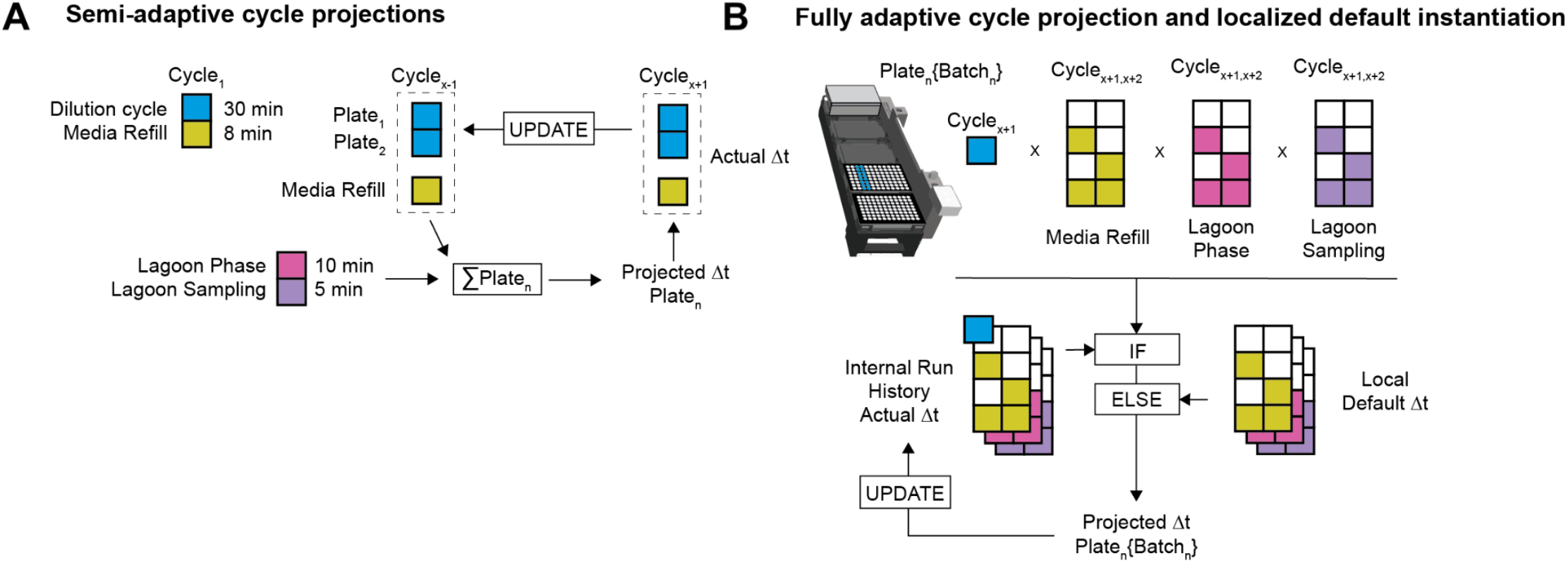
Semi-adaptive cycle projection improves stability but fails under mixed operational states. **A) Semi-adaptive projection.** Each plate initializes with a default cycle time that is updated using the most recent observed lap time. Media refill is recalculated dynamically, but lagoon phase and sampling increments remain fixed. This approach initially stabilizes turbidostats but breaks down once lagoon operations introduce heterogeneous and overlapping workloads across plates. **B) Fully adaptive projection**. Cycle time is instantiated per plate and per batch using the specific combination of active operations. If matching historical data exist, the most recent actual Δ t is reused; otherwise a localized default is applied. This batch-resolved adaptation accounts for mixed media refill, lagoon phase, and sampling states across adjacent cycles and maintains stable OD control where semi-adaptive timing diverges.

**Supplemental Figure 15.**
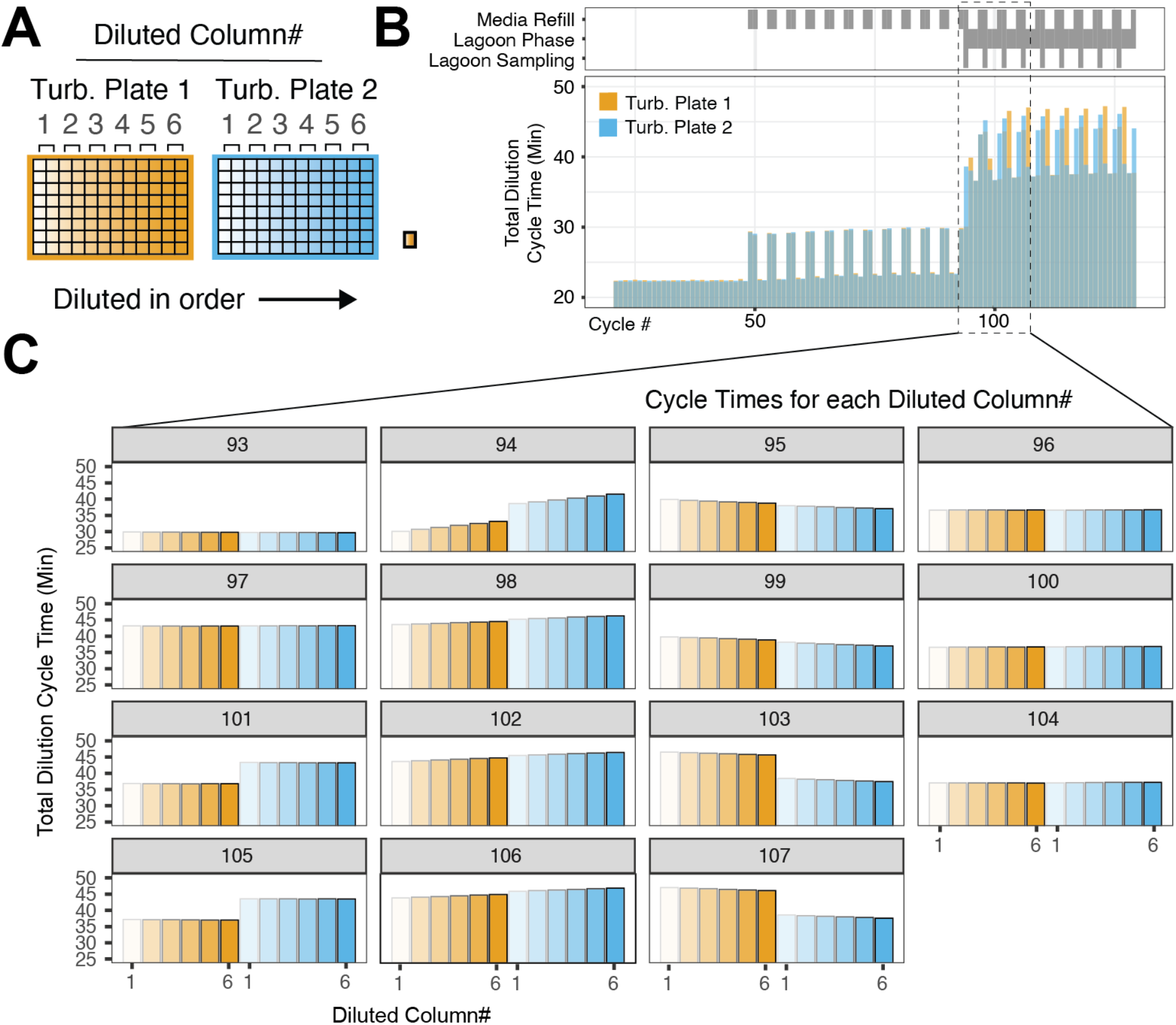
Lagoon sampling induces column-wise cycle-time skew within and across plates. **(A)** Turbidostat plates are diluted sequentially by column (1–6) for each plate. Each column is serviced in order, so any added workload during a cycle propagates to later columns in that cycle. **(B)** Total dilution cycle times for both plates across cycles. Media refill, lagoon phase, and lagoon sampling events (top) introduce abrupt increases in total cycle duration. The dashed region highlights cycles with lagoon sampling. **(C)** Per-column cycle times for representative cycles (93–107). During sampling cycles (e.g., 94, 98, 102, 106), later columns exhibit progressively longer service times within the same cycle. This skew persists into the subsequent cycle because the extended workload delays the next plate’s start time.

**Supplemental Figure 16.**
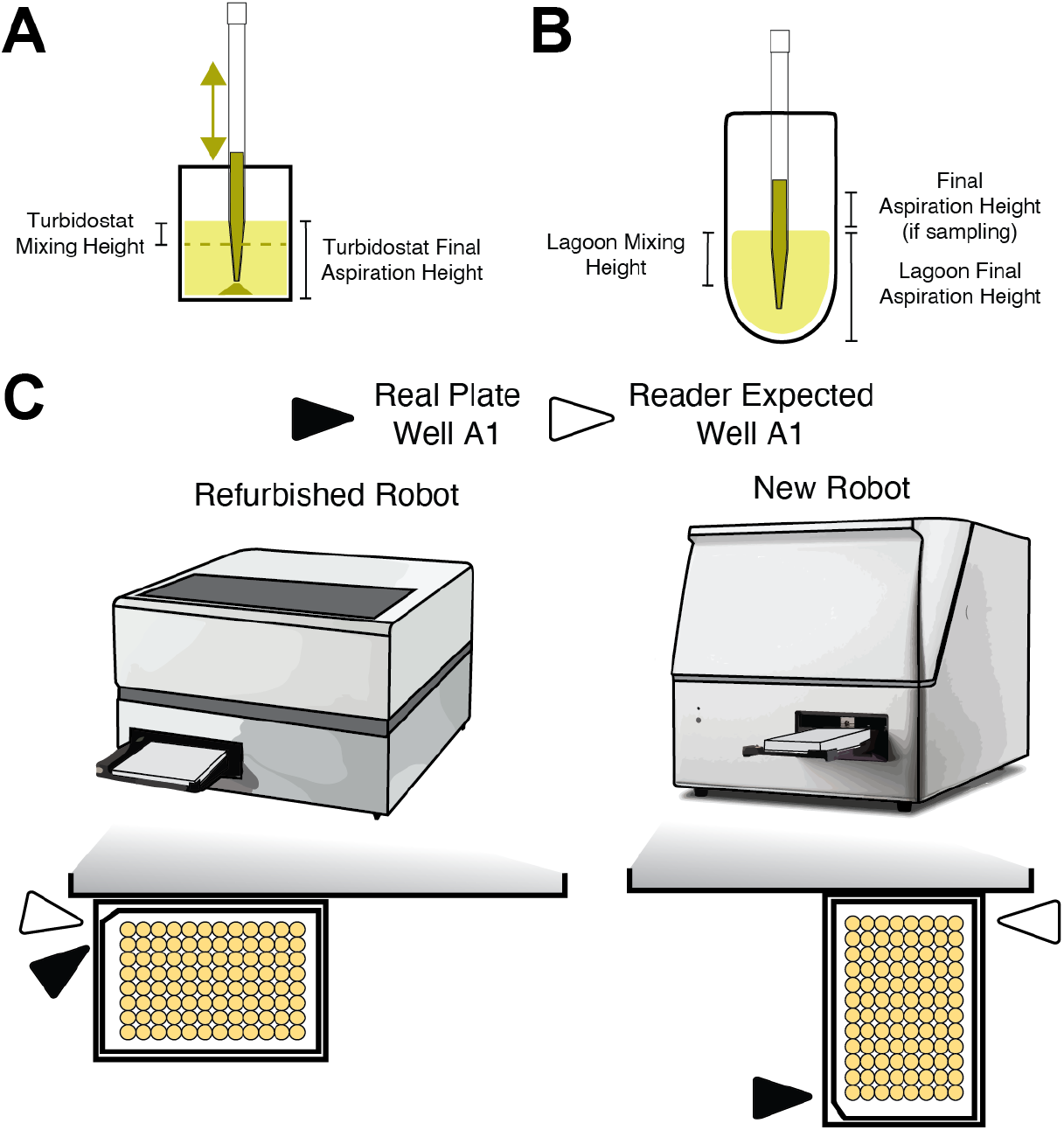
Cross-platform calibration of pipetting heights and plate-reader orientation. **(A)** Turbidostat height calibration. Mixing height and final aspiration height were calibrated on each platform to maintain constant culture volume and achieve sufficient resuspension without overflow or bubble formation. **(B)** Lagoon height calibration. Lagoon mixing height, final aspiration height, and sampling height were independently tuned to preserve steady-state lagoon volume and consistent sampling across platforms. **(C) Reader and deck orientation differences**. Differences in internal X, Y, and Z offsets and in plate-reader tray orientation (landscape vs portrait loading) result in reflected plate layouts relative to the reader’s expected well indexing (e.g., real plate well A1 vs reader-expected A1). These differences were resolved through platform-specific layout files, calibration parameters, and optional data reflection, enabling the same TurboPRANCE codebase to run reproducibly across distinct hardware generations.

**Supplemental Figure 17.**
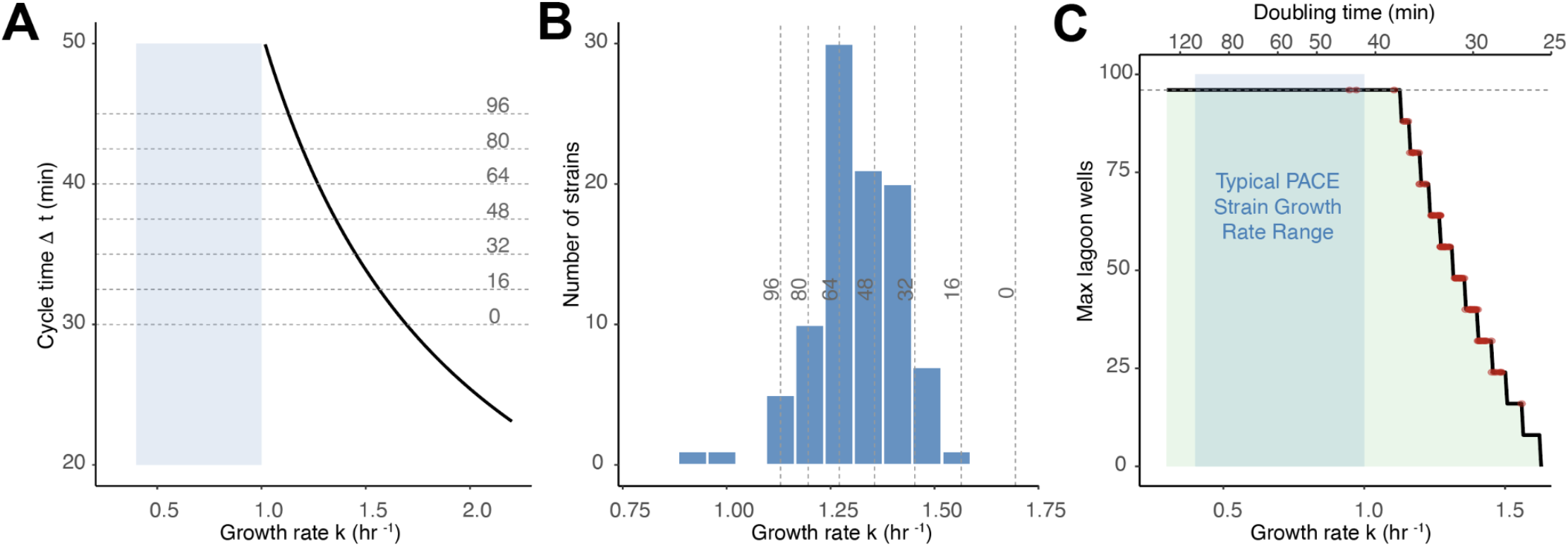
TurboPRANCE growth rate adjustments. **(A)** Maximum allowable cycle time as a function of growth rate (black curve), with horizontal dashed lines indicating cycle times for discrete lagoon configurations in increments of 16 (2 columns). Blue shading indicates typical PACE strain range; **(B)** Histogram of Keio collection growth rates with vertical dashed lines indicating the maximum supportable growth rate at each lagoon configuration (increments of 16). At full capacity (96 lagoons), 93 of 96 strains (97%) exceed the limit; reducing to 24 lagoons accommodates 95 of 96 strains. **(C)** Maximum number of lagoon wells supportable as a function of bacterial growth rate, with 96 Keio strains (DRM, no AP) overlaid as red points. Blue shading indicates typical PACE strain growth rate range (k = 0.4-1.0 hr^-1). red shading shows the Keio strain growth rate range.

**Supplemental Figure 18.**
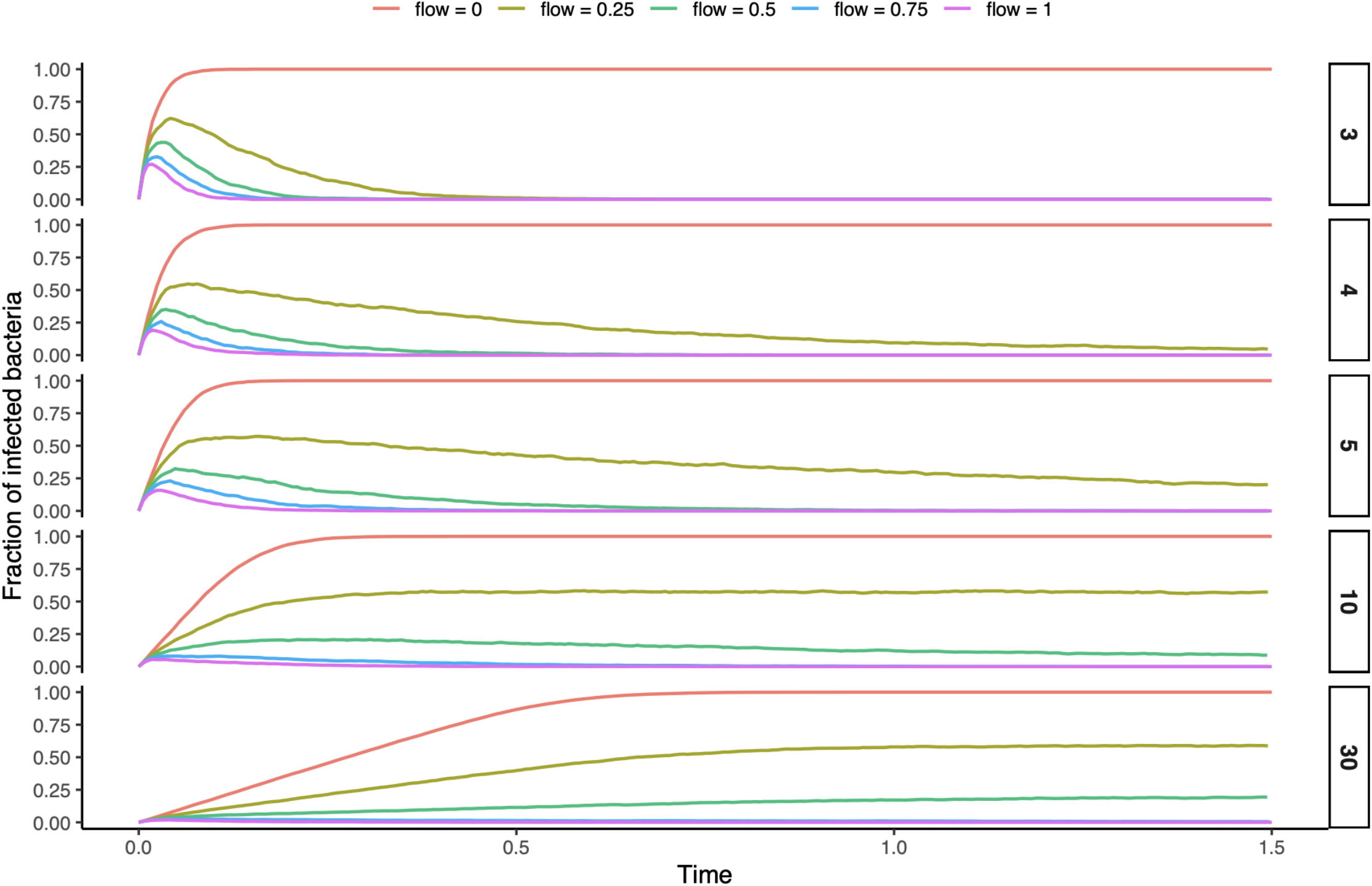
Monte Carlo simulation of M13 phage infection dynamics. Simulation across MOIs of 3, 4, 5, 10, and 30 and flow rates of 0-1 vol/hr. Each trace shows the fraction of infected bacteria over time. At low MOI, infection rapidly saturates the lagoon then declines as continuous flow of uninfected bacteria dilutes the infected population. At higher MOI, infection spreads more slowly and never fully saturates, with higher flow rates suppressing infection spread. With no flow, infection reaches fixation regardless of MOI.

**Supplemental Figure 19.**
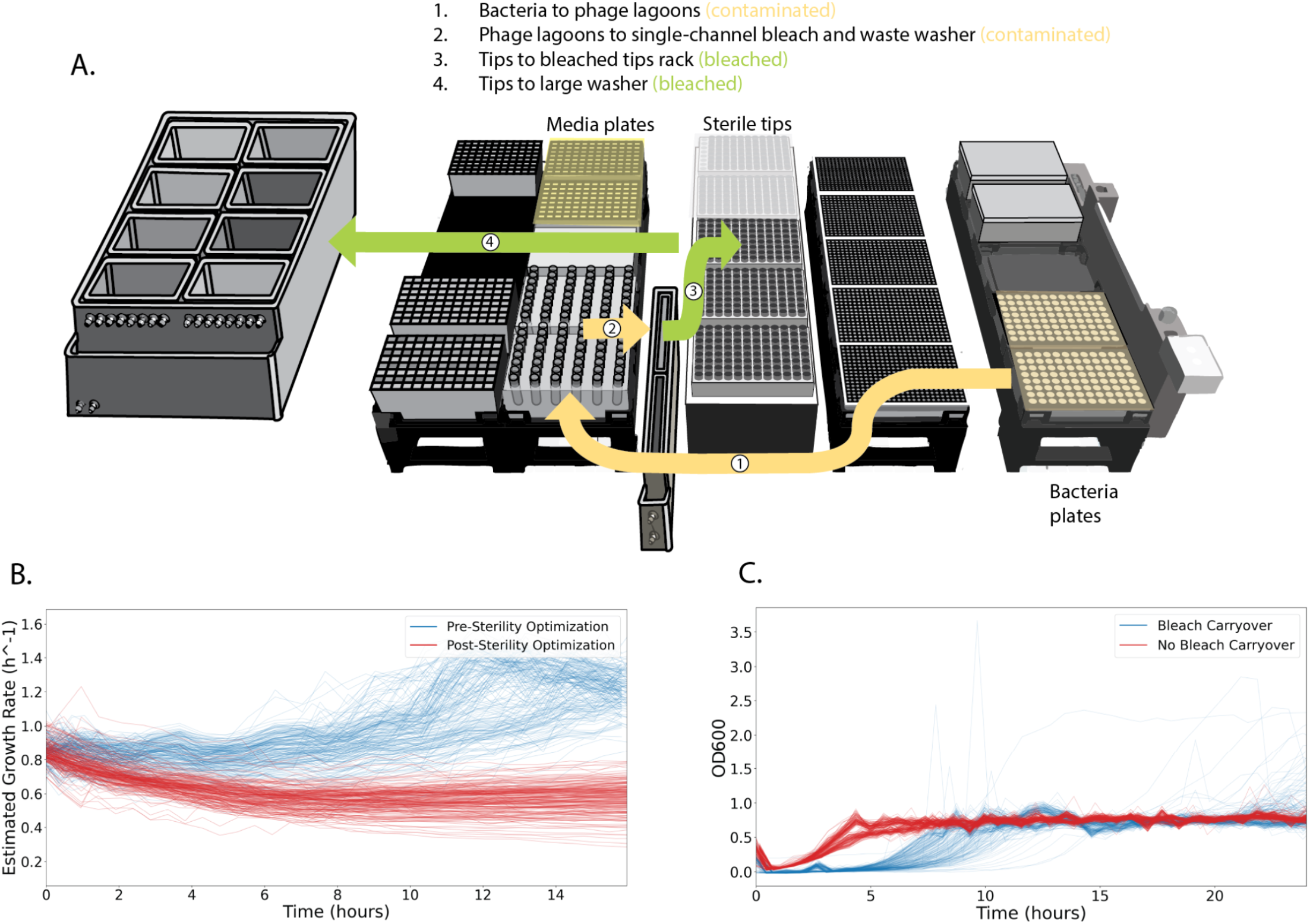
Sterility-optimized deck layout and validation after bleach carryover mitigation. **(A)** Deck layout and fluid path segregation following bleach carryover correction. Pipetting paths are arranged to minimize flyover of contaminated or recently bleached tips across media plates and phage lagoons. Bacteria plates and media plates are physically separated. The single-channel washer is positioned forward to increase distance from media plates. Recently bleached tips are isolated before reintegration, and waste is dispensed with tips positioned inside the washer reservoir to limit aerosolization. **(B)** Media contamination signature. Contamination of media staging plates produces abnormally high and non-convergent estimated growth rates due to inadvertent bacterial introduction into dilution media (pre-optimization, blue). After sterility optimization (red), growth-rate estimates converge and remain stable. **(C)** Bleach carryover signature. Residual bleach exposure results in prolonged lag phases and abnormal OD600 spikes consistent with dead-cell aggregates (blue). Following deck reconfiguration and tip segregation (red), cultures exhibit normal growth kinetics without bleach-associated collapse.

**Supplemental Figure 20.**
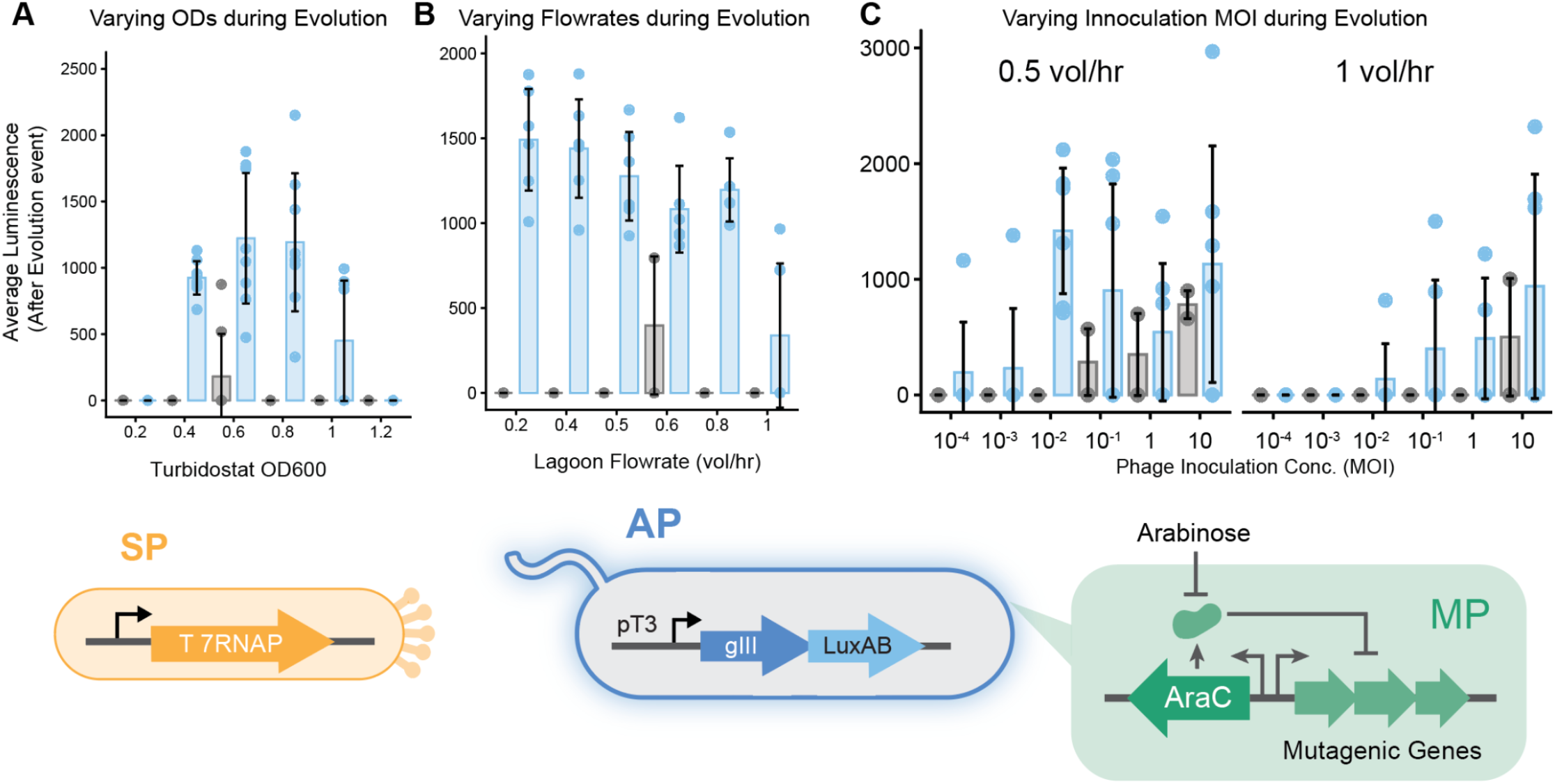
Quantification of T7RNAP evolution from pT7 to pT3 across operational parameters. Continuous evolution experiments were performed in which phage-encoded T7RNAP variants were selected for activity on a host accessory plasmid encoding pT3–gIII–LuxAB under mutagenesis induction. Arabinose-induced MP6 mutagenesis enabled diversification during selection. **(A)** Average post-evolution luminescence across turbidostat OD600 setpoints (0.2–1.2). Intermediate OD setpoints supported the strongest evolutionary signal, while extreme low or high ODs reduced output. **(B)** Average post-evolution luminescence across lagoon flowrates (0.2–1.0 vol/hr). Intermediate flowrates yielded the highest signal, consistent with balanced host supply and selection stringency. **(C)** Average post-evolution luminescence across initial phage inoculation concentrations (MOI 10^-4 to 10) at 0.5 and 1.0 vol/hr. Very low MOI delayed or reduced signal, while intermediate to high MOI conditions accelerated detectable evolution. Points represent independent lagoons; bars denote mean ± s.d. Schematic (bottom) illustrates the selection architecture: phage encode T7RNAP (SP), host cells carry a pT3-driven gIII–LuxAB accessory plasmid (AP), and mutagenesis plasmid (MP) expression is induced by arabinose.

**Supplemental Figure 21.**
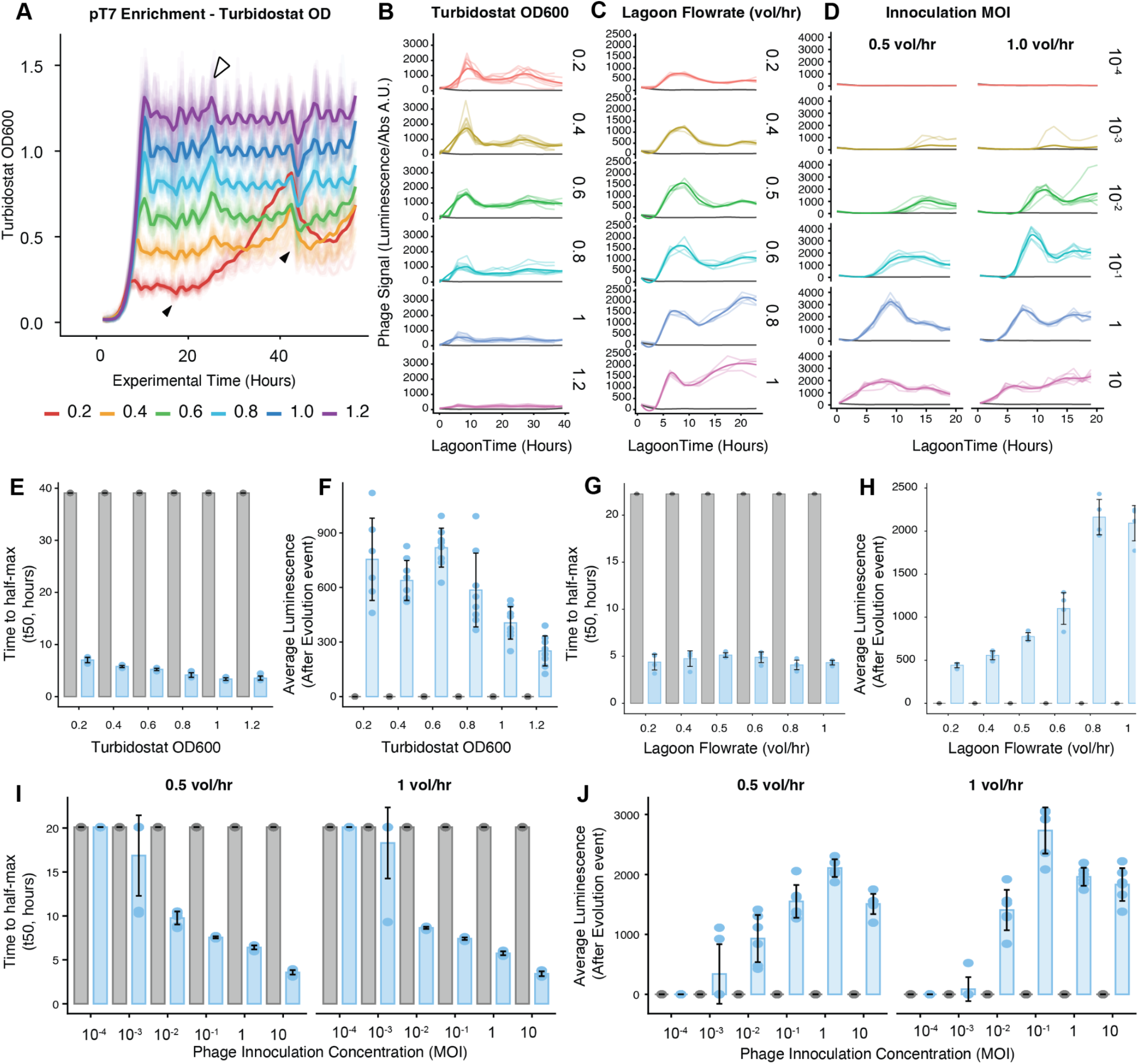
Quantification of T7RNAP pT7 selection across turbidostat and lagoon parameters. Selection-only experiments were performed without mutagenesis to quantify phage propagation as a function of turbidostat OD600 setpoint, lagoon flowrate, and initial multiplicity of infection (MOI). In all conditions, phage constitutively encoded T7RNAP, which activates pIII expression from a host pT7 promoter, enabling direct measurement of propagation without evolutionary change. **(A)** Representative turbidostat OD600 trajectories at indicated setpoints (0.2–1.2). Cultures were equilibrated prior to lagoon transfer. **(B)** Lagoon luminescence traces for turbidostats operated at different OD600 setpoints, reporting phage propagation over time. **(C)** Lagoon luminescence traces across lagoon flowrates (0.2–1.0 vol/hr) at fixed turbidostat OD. **(D)** Lagoon luminescence traces across initial MOIs spanning 10^-4 to 10 at two lagoon flowrates (0.5 and 1.0 vol/hr). **(E–F)** Quantification of selection outcomes across OD600 setpoints: time to half-maximal luminescence (t50) and average post-activation luminescence, respectively. **(G–H)** Quantification across lagoon flowrates. **(I–J)** Quantification across MOI conditions at 0.5 and 1.0 vol/hr.

**Supplemental Figure 22.**
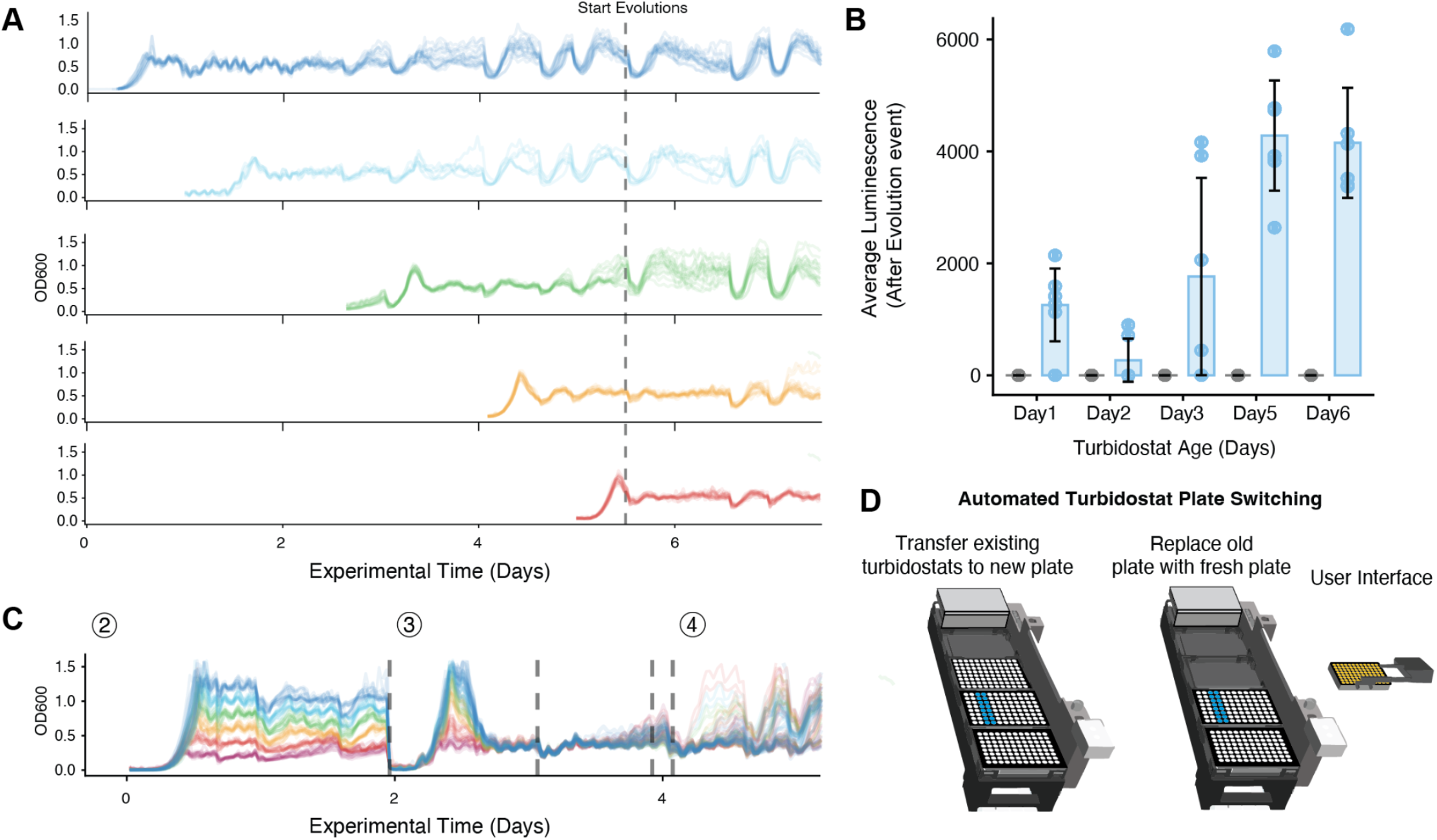
OD600 traces for turbidostats maintained during staggered, multi-day evolutions with automated plate switching. (A) Top, staggered MP6-containing turbidostats initiated and equilibrated at different timepoints prior to lagoon activation (vertical dashed line). Individual wells are shown as faint lines with mean trajectories overlaid. Dips and increasing instability in later timepoints reflect progressive difficulty maintaining homogeneous cultures during extended MP6 growth, consistent with biofilm accumulation and plasmid instability. (B) Quantification of post-activation luminescence as a function of turbidostat age at lagoon initiation. (C) Bottom, alternating plate-transfer strategy implemented in parallel experiments. Periodic bulk transfer of soluble culture to a fresh 96-well plate restores stable OD control and reduces biofilm-associated drift during long runs. (D) Schematic of automated turbidostat plate switching workflow used to replace aged plates without interrupting ongoing lagoon experiments. Together, these data show that while long-term MP6 maintenance leads to biofilm-associated OD excursions, staggered initiation and automated alternating plate transfer preserve stable operation across extended campaigns.

**Supplemental Figure 23.**
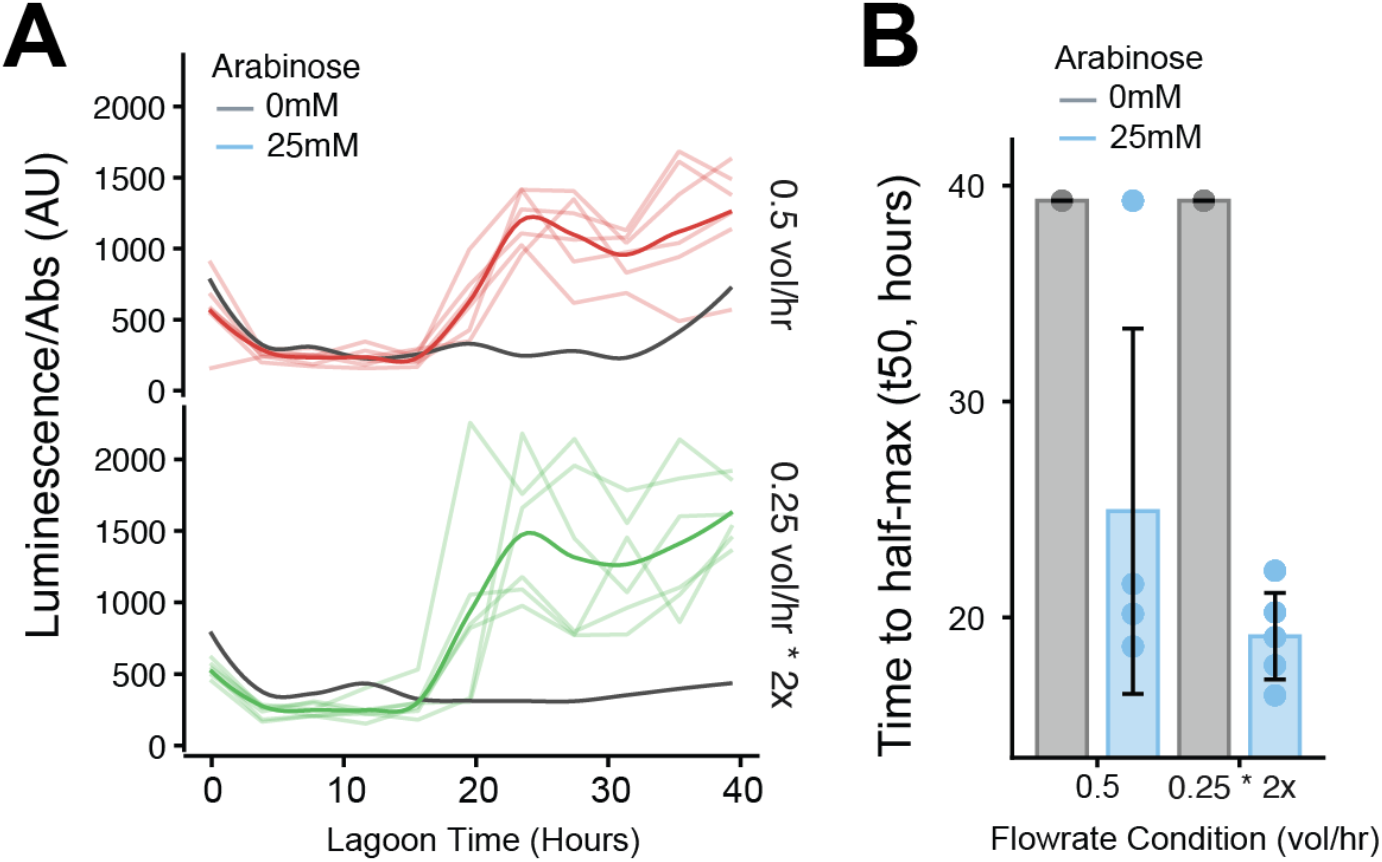
Programmable composition and scheduling of selection pressures during continuous evolution. **(A)** To test whether TurboPRANCE can implement mixed selection pressures within a single lagoon, T7RNAP-to-pT3 evolutions were performed under two pressure-delivery programs with the same total dilution rate (0.5 vol/hr). In the single-pressure program, lagoons were supplied by one pressure stream each cycle. In the mixed-pressure program, the same total inflow was partitioned across multiple independently maintained pressure streams within each cycle, creating a defined per-cycle mixture while keeping each stream under separate OD control. Evolution trajectories were comparable between programs, indicating that composing pressures does not compromise evolution while enabling mix-and-match selection schedules. **(B)** Quantification of evolution kinetics from A using time to half-maximal luminescence (t50). Bars show mean; error bars indicate s.d.

**Supplemental Figure 24.**
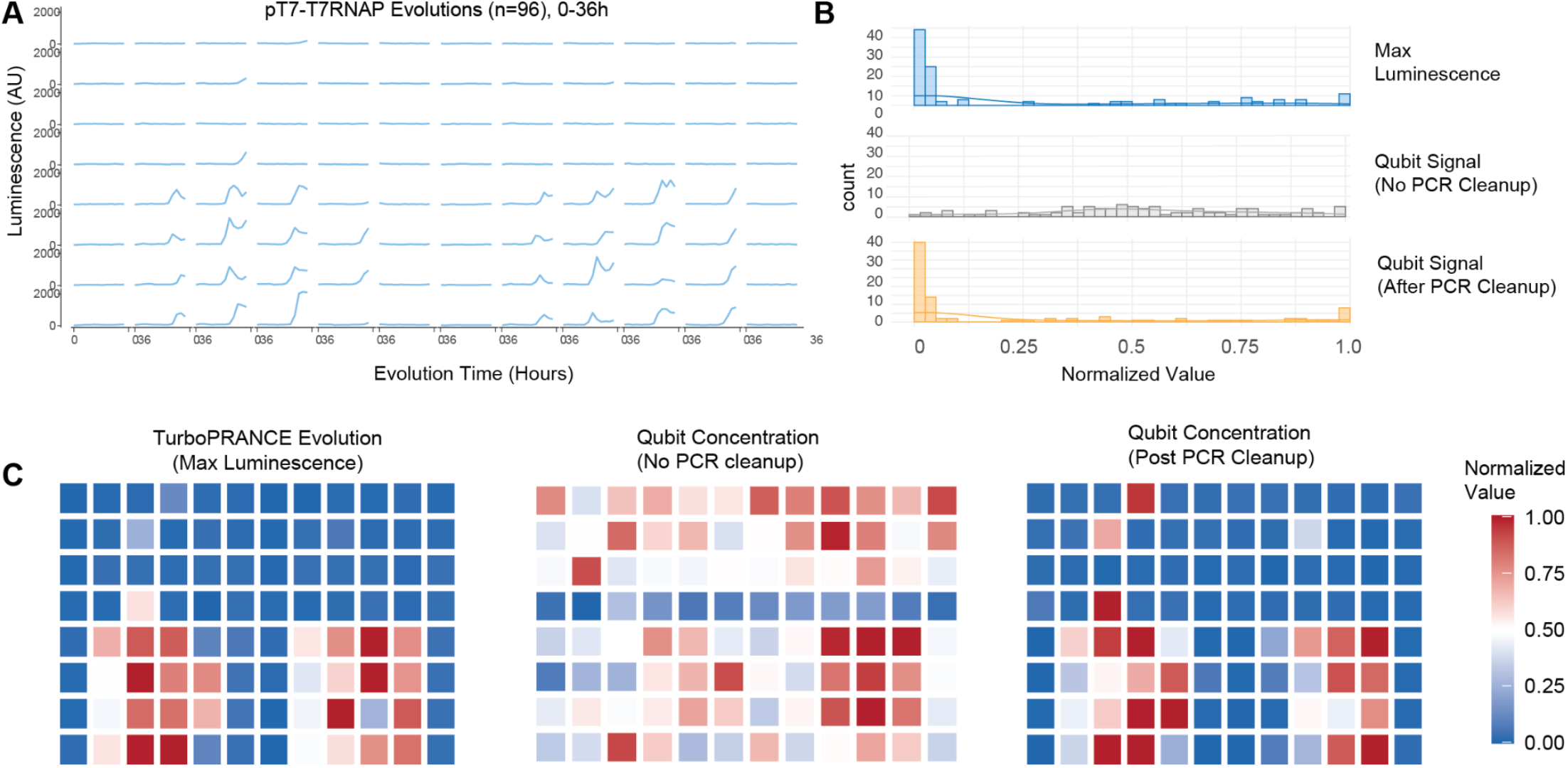
PCR cleanup enables quantitative pooling of TurboPRANCE Nanopore libraries. **(A)** Luminescence trajectories for 96 parallel pT7-T7RNAP evolutions (0-36 h); maximum luminescence per lagoon was used as an orthogonal proxy for phage propagation. **(B)** Distributions of normalized values for maximum luminescence (top), Qubit dsDNA signal measured directly from PCR products without cleanup (middle), and Qubit signal after bead-based PCR cleanup (bottom). Without cleanup, Qubit measurements were artificially compressed across wells, consistent with background fluorescence from residual primers and short products. After cleanup, Qubit measurements spanned a broader dynamic range. **(C)** Well-wise heatmaps of maximum luminescence (left), Qubit concentration without cleanup (middle), and Qubit concentration after cleanup (right), each normalized to the same 0 to 1 scale. Post-cleanup Qubit concentrations more closely recapitulate the well-to-well heterogeneity in evolutionary outcomes, supporting more accurate equimolar pooling prior to sequencing.

**Supplemental Figure 25.**
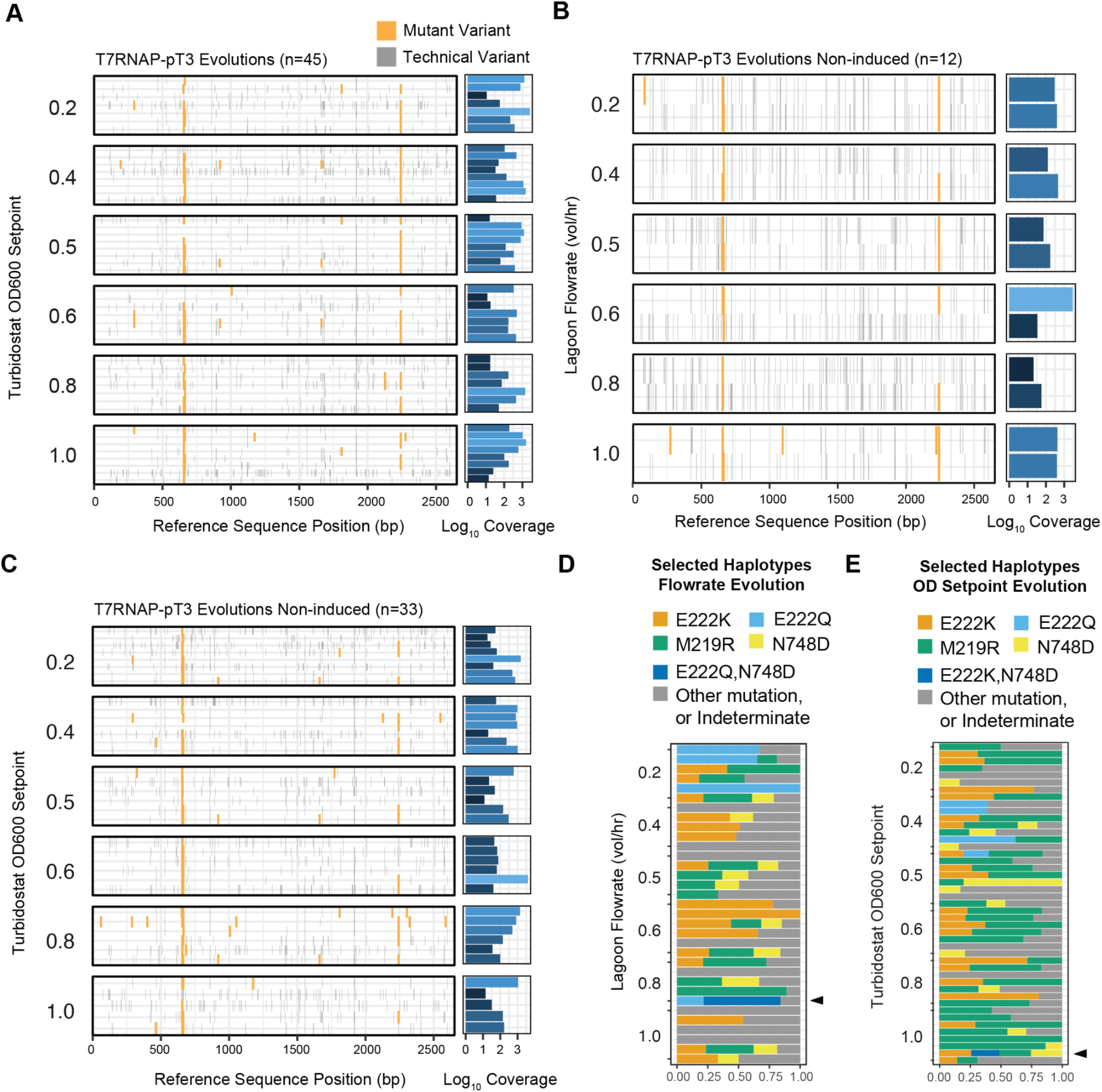
Variant maps for 83 independent evolutions. **A)** All called variants for lagoons evolving T7RNAP for pT3 activity across a variety of turbidostat OD setpoints. Coverage represents the number of mapped reads for each sample **(B**,**C)** Called variants for lagoons that were not induced with arabinose. Commonly evolved mutations are still observed due to natural phage mutation in the absence of induced mutagenesis even though in many cases luminescence is not yet detected. **(D**,**E)** Haplotype phasing for common mutations of all induced evolution samples from setpoint and flowrate pT3-T7RNAP experiments. Black arrows indicate lagoons with detected double mutants.

**Supplemental Figure 26.**
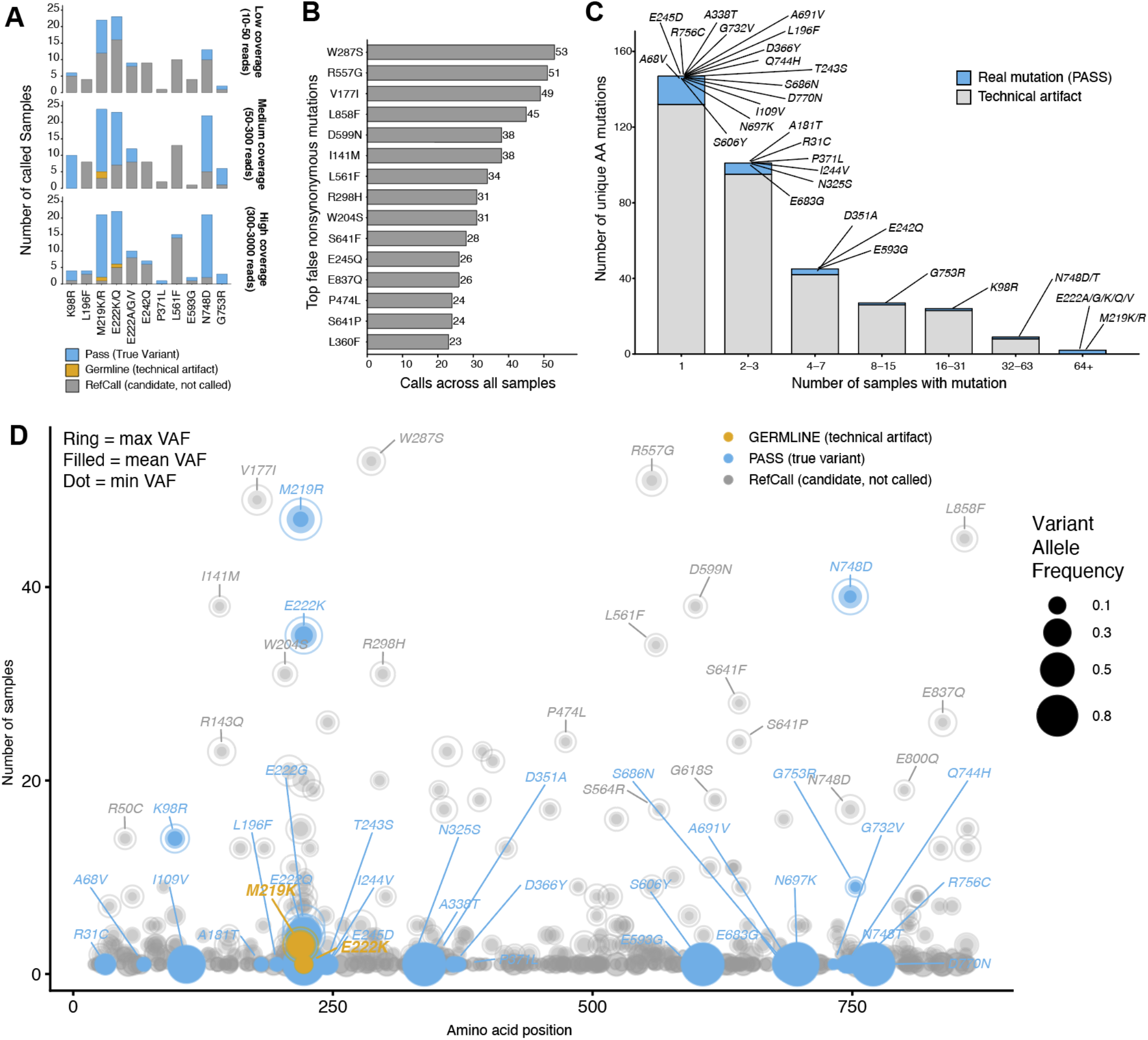
Variant classification, coverage dependence, and recurrence analysis. **(A)** Stacked bar plot showing the number of variant calls classified as PASS (true variant), GERMLINE (technical artifact), or RefCall (candidate not called) at each PASS-validated nonsynonymous position, faceted by sequencing coverage tercile (Low, Medium, High based on mapped read counts). Only positions detected in all three coverage levels are shown. Labels indicate the amino acid change(s) at each position. Higher RefCall counts in low-coverage samples reflect missed real mutations due to insufficient read depth. **(B)** Top 15 false nonsynonymous mutations from the RefCall category, ranked by the number of calls across all samples. These represent recurrent technical artifacts that would be reported as real mutations without DeepSomatic filtering. **(C)** Distribution of unique nonsynonymous mutations by the number of samples in which they were detected. Mutations are classified as real (PASS in at least one sample are shown in blue) or technical artifact (never classified as PASS, grey, mutation labels not shown). Individual PASS mutations are labeled. **(D)** Scatter plot of all recurrent nonsynonymous variants by amino acid position and number of samples, colored by DeepSomatic classification (PASS, GERMLINE, RefCall). For each variant, the open ring shows the maximum VAF across samples, transparent circle corresponds to the mean VAF, and the filled dot shows the minimum VAF between samples.

## Supplemental Methods

### Bacterial Propagation and Preparation

All experiments performed in this study were conducted using S2060 cells (Addgene, Cat #105064). Cell line maintenance, building, and competent cell generation was performed using 2XYT media (US Biological, Cat# T9200) with the appropriate antibiotics including Carbenecillin, Kanamycin, Chloramphenicol, Spectinomycin, and or Streptomycin. All chemically competent cells were generated using the Mix N’ Go Chemically Competent Transformation Kit with cells grown in 2XYT (Zymogen Cat# T3001).

### Cloning Phage

New M13 phage constructs used in this study were generated using In-Fusion assembly (Takara #Cat 638949) with fragments obtained from high-fidelity PCR (either PrimeSTAR Max v2, Takara #Cat R047A or PrimeSTAR GXL, Takara #Cat R050A). After gel purification, all amplicons were treated with DpnI (NEB, Cat# R0176) and purified again via column purification before being used in In-Fusion half reactions with <200ng total DNA. Reactions were transformed into chemically competent S2060 cells containing pED12x0 (A kanamycin-resistant derivative of SC101 psp-pIII, Addgene #79219) expressing pIII under a phage shock promoter. We used a standard transformation protocol, except instead of plating cells at the end of a 2hr recovery step in 2XYT we added 3ml 2XYT media containing Kan and incubated the liquid culture overnight, shaking at 37C. The following day the liquid culture was pelleted at 3000g and the supernatant was filtered to purify polyclonal phage. To isolate clonal phage we performed phage plaque assays as previously described^3^. In brief, overnight, confluent S2060 cells containing pED12x0 were diluted 1:100 in fresh 2XYT containing Kan and grown at 37C to reach an OD of 0.4. Then, cells were combined with a series of 100-fold dilutions of polyclonal phage and 2XYT + 0.5% Agar supplemented with 1% Bluo-gal (Goldbio, Cat# B-673-250). Dilutions were then evenly plated on 2XYT+2% Agar quadrant petri dishes and incubated face up overnight at 37C. Distinct blue plaques representing clonal phage were picked using a pipette tip and used to directly inoculate liquid 2XYT containing kanamycin which was grown overnight. Filtered supernatant of clones was obtained and then submitted for rolling circle amplification and sanger sequencing.

### Phage Titering by qPCR

To quantify phage, cultures were first pelleted at 3000g and the supernatant was filtered to remove bacterial genomic and infected M13 genome DNA (mdi Membrane Technologies, Cat# CFPL2101XXXX203). To remove background M13 genome DNA from polyphage a thermocycler was used to incubate supernatant at 80C for 30min and then samples were treated with DNAse I (GoldBio, Cat# D-303) for 30min at 37C before denaturing the sample at 95C for 5min. Samples were then diluted 1:5 in ultrapure water (CAT#) and then 2ul was used as input into absolute quantification by qPCR with Luna 2X qPCR Master Mix in 10ul total reactions (NEB, Cat# M3003). The following primers were used for titration: 5’-TGTTCTCGATGAGTGCGGTA-3’ and 5’-TTCTGTCCAGACGACGACAA-3’. We then converted Ct values to phage counts using a calibration curve generated between our protocol and phage plaque counting assays. Any samples with Ct values below the quantitative range were rediluted and requantified to obtain accurate quantities.

### Media preparation for automated experiments

All TurboPRANCE and automated experiments used DRM based media prepared in carboys and pumped on deck. To prepare 4L of complete DRM, 79.5g of DRM-A (US Biological, Cat# CS050H-001) and 1150ml of Tween 20% (Sigma-Aldrich, Cat# P1379) was added to MilliQ water and autoclaved using a 1hr liquid cycle. The liquid was allowed to cool overnight at 4C or in a room-temp water bath for several hours before 100ml DRM-C was added. To prepare DRM-C, 115g of DRM-C (US Biological, Cat# CS050H-003) was mixed with 400ml of MilliQ water and 120ul of Trace Metals (Final: 30nM (NH_4_)_6_Mo_7_O_24_·4H_2_O, 4uM H_3_BO_3_, 550nM CoCl_2_, 160nM CuSO_4_, 1.26uM MnCl_2_, and 100nM ZnSO_4_·7H_2_O) and 10ul of 1M CaCl was added to create 500ml of DRM-C. Sterile filtration using a 0.45um filter was then performed before addition to cooled DRM-A. For all experiments carboys contained 50mg/L of Streptomycin (Fisher Scientific, Cat# AAJ6129922). To simplify preparation and to avoid potential degradation of stock antibiotics across long experiments on the 37C heated robot deck, antibiotics were included in the media carboys if possible. Additionally, for experiments with MP6 containing bacteria, half the typical working concentrations of antibiotics were used to stabilize MP6, consistent with previous PACE studies.

### Statistical analysis of lagoons

To generate t50 and mean normalized luminescence values, luminescence readings from sample plates were first normalized to absorbance values. An emerging functional variant propagating in a lagoon (evolution or enrichment) is assumed to increase exponentially before plateauing and decreasing over time consistent with random-walk modeling. Therefore, binomial regression was used to quantify the kinetics of normalized luminescence. The following processing and filter criteria were used to minimize false positives and to ensure appropriate binomial fitting. First, a threshold was calculated using the mean of normalized luminescence from all control lagoons of an experiment multiplied by 2. For every individual lagoon, at least 2 data points were required to pass the threshold, else t50 was set to the maximum experimental time. This approach effectively filtered out lagoons where no or highly delayed luminescence were observed. Then, for each experiment, an early timepoint cutoff was used to avoid the impact of early measurement noise which otherwise resulted in premature binomial fits. It was challenging to fit binomial regressions for some samples because their luminescence trajectories showed early and immediate spikes, or strong spikes later on during evolution. This prevented fitting due to lack of an early baseline and or weighted t50 values too heavily towards later luminescence values in evolution respectively. Therefore, next, 3 additional “padding” data points were imputed both before the earliest timepoint and after the latest timepoint with values corresponding to the minimum and maximum luminescence values detected over the course of evolution. After processing and binomial fitting, any t50 values over the maximum experimental time were capped at the maximum experimental time. It is noted that this approach fails in cases where luminescence increases monotonically with non-exponential slopes, often seen when multiple variants are co-evolving or a single variant has acquired additional advantageous mutations before dominating a lagoon. This artificially inflates t50 when a fit is attempted using binomial regression. As these values are typically much higher than experimental time they generally default to the maximum experimental time. Lastly, using t50 values, the mean normalized luminescence was calculated by averaging the normalized luminescence values for all available time points within 12 hours after the t50 for that lagoon.

### Robot platform and hardware configuration

The liquid-handling robot we used was a Hamilton STAR with an eight-channel dispensing head, a 96-channel dispensing head, and a plate gripper robot arm. Placed within reach of the robot arm was a BMG VantaSTAR plate reader capable of absorbance and luminescence measurements. Inside the robot enclosure was a custom 3D printed eight-compartment washer module **(Supp Fig. 3)** with two independent flow channels on each compartment. Additionally inside the robot enclosure was a 3D printed single-channel washer with two compartments. All consumable labware such as 96-well plates were placed on plate carriers inside the robot enclosure. A complete list of parts including models for 3D printed parts is included in the supplementary information.

### TurboPRANCE Protocol Overview

TurboPRANCE is run on a Hamilton STARlet liquid-handling robot using a Python script interfacing with the PyHamilton software package and other libraries for controlling auxiliary equipment. The process starts in the turbidostat phase, in which the robot maintains up to 192 bacteria cultures at a density setpoint in parallel by periodically reading the culture density and diluting with fresh media. During the turbidostat phase, excess culture media is disposed of by dispensing into the waste compartment of the single-channel washer. At the beginning of an experiment, the bacteria cultures are seeded at a low density (e.g. 1:500 dilution) and media dilutions are minimal. Once the turbidostat densities have equilibrated at the setpoint, the user manually triggers a transition to the lagoon phase, in which the robot transfers culture from the turbidostats to specified phage lagoons. During the lagoon phase, the robot periodically samples from phage lagoons into 384-well plates to track lagoon culture density and luminescence. Once the lagoon cultures have equilibrated to the desired culture volume, the user manually infects lagoons with bacteriophage to begin the evolution process.

### Turbidostat culture maintenance

Up to 192 independent cultures of bacteria can be maintained in parallel as part of the turbidostat module of TurboPRANCE. The objective of the turbidostat module is to transfer the optimal volume of media into a culture to maintain the culture at the density setpoint. The calculation for determining the optimal transfer volume uses an exponential growth model *P* = *P*_0_*e*^*kt*^ where *k* is a growth-rate constant determined from historical data. Future population density values are estimated based on the projected time until the next measurement step, and media transfer volume is calculated as the volume needed such that the next density measurement is at the setpoint. Since density measurements are noisy, we use damping and weighted averages to ensure that estimates of *k* are not overly perturbed by single measurements. The equation for the media transfer volume that dilutes the culture to the density setpoint at the next measurement time is *V*_*medi*_ = *(OD*_600, *next*_/*OD*_600, *now*_−1)∗*V*_*culture*_. Media is aspirated from a deep-well plate and dispensed into turbidostat cultures using the robotic eight-channel pipetting arm of the Hamilton STAR. Then, the same volume of culture is aspirated in order to maintain the cultures at constant volume. In the case of a pure turbidostat experiment, all of the excess culture is dispensed into a 3D-printed waste trough and pumped off-deck. During the lagoon phase of a TurboPRANCE experiment, a specified portion of the excess turbidostat volume is dispensed into the phage lagoons in order to supply constant density culture.

### Lagoon transfers and sampling

TurboPRANCE enables real-time user control over the allocation of bacteria cultures from the turbidostat culture wells to the phage evolution lagoons. During every cycle iteration, the TurboPRANCE Python script reads a spreadsheet that specifies the lagoon destination and volumetric flow rate of every turbidostat culture (including null destinations). As the script loops over turbidostat cultures, it dynamically constructs a dictionary mapping from turbidostat source wells to destination lagoons. This spreadsheet can be manually or automatically updated in real-time during the TurboPRANCE experiment, providing a high degree of control for fine-tuning the selection stringency of experiments. The Python script also dynamically constructs a mapping from phage lagoons to sampling wells, so that the lagoon origin and time point for each sample can be easily determined for downstream processing and analysis. After dispensing fresh media and aspirating excess culture, the robot dispenses the difference between the aspirated volume and the culture volume specified in the destination mapping to waste. The robot then dispenses the requested volume into the lagoons, and if sampling, aspirates a sample from the lagoon and dispenses these to the sampling plate. The cycle continues until all of the turbidostat wells in the current plate have been serviced. The robot then moves the sampling plate to the plate reader and the absorbance and luminescence for each of the sampled wells is measured and recorded in a database. Finally, the robot uses the 96-channel head to dispense an arabinose solution into the bacteriophage lagoons, which induces the mutagenesis plasmid. A dynamic sampling system is implemented in the Python script in order to allocate samples to appropriate positions in 384-well plates and record these positions for downstream processing.

### Turbidostat culture batching

The upper limit of lagoon flowthrough rate is governed by the doubling time of each host strain and the steady-state biomass production of its supplying turbidostat well. For burdensome strains (for example, those carrying mutagenesis plasmids with effective growth rates around ∼0.6 hr^-1), a single turbidostat well limits the sustained volumetric outflow that can be delivered to a 350 uL lagoon at higher target dilution rates (for example, >0.5 vol/hr). To increase available host supply, columnwise batching is used, where a single strain is distributed across paired (n>1) turbidostat wells and their excess outflow is pooled to service the same lagoon. During batched processing, the eight-channel pipetting head picks up one set of tips and aspirates the total media volume required to dilute all wells in the batch. The channels then dispense the appropriate volume of media into each batched well and aspirate excess culture from each well using the same tips, without intermediate sterilization between wells within the batch. The pooled aspirated culture from the batch is then directed to the specified lagoon. Batches are arranged along rows so the columnwise head traverses horizontally during processing. The controller manifest specifies control parameters at the batch level, applying a shared setpoint and transfer configuration across all wells in that batch.

### Sterility-optimized component separation

Deck layout and fluid routing are designed to spatially separate contaminated, bleached, and sterile components to reduce cross-contamination between turbidostats, media plates, and phage lagoons **(Supp Fig. 19), (Supp Fig. 6)**. Media staging plates and sterile tip racks are positioned distal to lagoon plates and bleach-containing washer modules. The single-channel washer is positioned forward on the deck to increase physical distance between bleach reservoirs and media resources. Tip roles are segregated by function. Media formulation tips, turbidostat dilution tips, lagoon induction tips, and dirty staging tips are maintained in distinct racks. Recently bleached tips are first transferred to a designated “dirty” staging rack before re-sterilization with the 96-channel head and return to a clean rack, preventing bleach splashback into sterile tip positions. Pipetting paths are constrained to avoid flyover of contaminated tips above sterile labware. During lagoon sampling and transfer, clean tip racks are positioned outside the trajectory corridor of phage-contacting tips. Waste dispensing is performed with tips positioned inside washer reservoirs rather than above them to limit aerosolization. Bleach is replenished in waste compartments immediately after dispensing to maintain antimicrobial activity.

### Media Preparation & Media Line Flushing/ Sterilization

All automated experiments are supplied with bulk DRM-based media prepared in 4 L carboys and delivered on deck via a dedicated peristaltic pump array. DRM-A is prepared by dissolving 79.5 g DRM-A powder and 1150 uL Tween-20 in 4 L Milli-Q water and autoclaving on a 1 hr liquid cycle. The media is allowed to cool completely before addition of 100 mL sterile-filtered DRM-C (prepared separately and filtered through a 0.2 to 0.45 um filter). Antibiotics are added either directly to the carboy or into DRM-C prior to addition, depending on the experimental configuration. To prevent biological or bleach carryover between runs, media lines are flushed immediately before connecting sterile media. The media intake line is disconnected from any existing carboy and submerged to the bottom of a graduated cylinder containing 1 L of 5% bleach. Using the pump control utility, the line is flushed twice for 30 s each. The line is then flushed twice using 1 L sterile water with the same timing. After flushing, the tubing end is sprayed with ethanol and capped with a sterile luer lock until connection to a freshly prepared sterile media carboy . On-deck reservoirs used for media delivery and washing are drained and refilled as needed at the start of an experiment. The bleach carboy is filled to 20 L and adjusted to 5% final bleach by adding concentrated bleach during the fill. The water inlet valve is closed after filling to prevent continuous dilution. If required, washer and waste reservoirs are emptied using the pump control interface prior to method initiation

### Evolution feedback control

Evolutionary stringency in TurboPRANCE is governed entirely through the controller manifest, which is reloaded at the beginning of every cycle. The manifest defines turbidostat OD setpoints, lagoon destinations, lagoon flow rates, strain identity, and batching configuration. Any of these parameters can be modified during a run to implement feedback control. No changes to the core robot method are required; updated manifest values are applied automatically on the next cycle. Lagoon flow rate and supplying host strain are the primary levers for modulating selection pressure, consistent with classical PACE implementations. However, the framework is agnostic to how control decisions are generated. Experimental data are written continuously to a database and can be processed externally by any analysis workflow. The resulting control decisions are implemented by programmatically updating the manifest file. Feedback strategies can range from simple rule-based thresholds to advanced model-driven controllers. For example, lagoon flow rates can be increased once luminescence exceeds a defined cutoff, reduced after prolonged absence of signal, or stepped according to elapsed time. Strains can be reassigned based on growth-rate stability, OD deviations, reporter intensity, or predefined stringency schedules. More advanced implementations can incorporate rolling averages, adaptive cutoffs, sequencing-derived genotype frequencies, fitness estimates, or manual review checkpoints. Because control logic is decoupled from the robotic execution layer and mediated only through manifest updates, feedback controllers of arbitrary complexity can be implemented without modifying the underlying method.

### Asynchronous parallelization

Individual cultures can be initiated asynchronously while a TurboPRANCE process is underway by adding cultures to turbidostat plate wells and updating the manifest to include the relevant control parameters and experimental metadata for each new turbidostat. In each cycle, the Python script reads the manifest, and if new rows have been added, the script generates new controller objects to account for the new turbidostat cultures, and likewise terminates objects whose rows have been deleted.

### Media refilling

In order to continuously supply fresh media to the bacteria wells, a media refilling system is used to transfer media from a 5 liter vessel to deep-well plates on the robot deck, from where it can be used to dilute turbidostat cultures. The process for refilling media involves pumping fresh media from a 5 liter vessel into a dedicated media compartment in the eight-compartment washer. Media refilling is triggered based on volume trackers that account for the total amount of media added to and withdrawn from wells in the media plates during turbidostat maintenance.

### Tip washing

Due to the continuous and long-term nature of TurboPRANCE protocols, it would be unrealistic to use a new set of pipette tips for each volume transfer (>200 pipetting steps per hour). An automated system for sterilizing and washing pipette tips was developed in order to reuse tips continuously over the course of an experiment. This tip washing system consists of two 3D printed washers with separate compartments for bleach, water, or waste, a set of automated peristaltic pumps for loading and unloading washer fluids from the system, and a function within the TurboPRANCE script for submerging used pipette tips with the 96-channel pipetting head into the washer compartments. Washer fluid is into the washer from 20L carboys positioned below the robot deck. Detailed schematics for the pumping system and 3D printed washer are provided in the supplementary information.

### High-throughput library preparation of evolution samples

#### PCR amplification of lagoon samples

To analyze evolution endpoints or trajectories, individual lagoon samples were collected into 384-well plates and retrieved from the robot deck, centrifuged for 5 min at 3000g, and then stored at 4C sealed with foil covers until library preparation. All subsequent liquid handling steps were performed either manually using multichannel pipettes or using an Opentrons OT-2 liquid handler. During aspiration steps, care was taken to aspirate at heights that avoided disturbing pelleted bacteria. For each sample, 10 uL of phage supernatant was diluted 1:10 in ultrapure water to 100uL total and filtered in 96-well 0.2uM filter plates (Pall Corporation, Cat# 8019) to remove residual bacteria. PCR reactions were assembled in 384-well PCR plates (Thermo Fisher, Cat #AB-1384). Each reaction contained 8.5 uL Q5 polymerase master mix (NEB, Cat #0491), 1.5 uL of 10 uM premixed forward and reverse primers, and 5 uL of filtered phage supernatant, for a total reaction volume of 15 uL. Reaction components were combined according to the manufacturer’s recommendations. Plates were foil sealed and PCR was performed in a 384-well thermocycler using the following program: 98C for 30 seconds, followed by 10 to 12 cycles of 98C for 10 seconds, 60C for 30 seconds, and 72C for 3 minutes. Following PCR, samples were transferred to a 96 well plate and amplicons using a homemade SPRI bead suspension (see below) as previously described^57^. Briefly, SPRI beads were mixed thoroughly with DNA samples at a 0.5x ratio.and incubated at room temperature for 5 minutes to allow DNA binding. Plates were placed on a 96 well magnetic rack until the solution cleared and beads were fully collected. The supernatant was removed, and beads were washed twice for 30 seconds with freshly prepared 80% ethanol without disturbing the pellet. After the second wash, residual ethanol was removed and beads were air-dried for 3 to 5 minutes until dry. DNA was eluted in nuclease-free water, incubated at 37C for 3 minutes, and placed back on the magnet. The supernatant containing purified, size-selected DNA was transferred to a new plate for downstream quantification.

#### Post-PCR dilution and 384-well Qubit quantification

Following amplification and purification, PCR products were diluted 1:10 by transferring 10 uL of purified PCR product into 95 uL of ultrapure water in a 384w plate. These diluted products were quantified using a Qubit High Sensitivity dsDNA assay adapted to a 384-well format. For quantification, 18 uL of Qubit HS dsDNA buffer and reagent mix was dispensed into each assay well, including wells designated for samples, two provided standards (0ng/ul and 20ng/ul). If a plate contained 384 samples, a second 384-well plate was used exclusively to measure standards and controls. To each assay well, 2 uL of diluted PCR product or control was added. Plates were read on a BMG VANTAstar plate reader without a lid using the following fluorescence settings and filters: Bottom optic reading, point-scan, max precision, Ex: F:482-16, Dichroic F:LP504, Em: F:530-40, at ∼800 gain. . Fluorescence measurements were used to generate a linear standard curve, from which molarity for each sample was calculated. If a sample exceeded the upper range of the standard curve, an additional 1:10 dilution was performed beginning from the first diluted plate, resulting in an effective 1:100 dilution relative to the original PCR reaction. Quantification was repeated using this further dilution.

#### Sample Pooling and SPRI purification

Samples were pooled in equimolar amounts into one or more 1.5 mL Eppendorf microcentrifuge tubes using an Opentrons OT-2 liquid handler controlled by a PyLabRobot script. Pooling volumes were calculated based on measured molarity, and transfers were performed from whichever dilution plate provided reliable concentration measurements. Reference phage samples were added at this stage if required. Pooled libraries were concentrated using a combination of homemade SPRI beads (see below) and sample evaporation on a 60C thermoshaker. After bead purification, libraries were assessed either by Qubit or by an Agilent TapeStation 2100 using D5000 ScreenTape and associated reagents (Agilent, Cat# 5067-5588) to evaluate fragment size distribution and to determine a distribution-based molarity for input into the Nanopore library preparation workflow.

#### Nanopore library preparation and sequencing

Sequencing libraries were prepared using the Nanopore Sequencing Ligation Kit v14 (Oxford Nanopore Technologies, Cat #SQK-LSK114), with modifications to the manufacturer’s protocol. For end-preparation, 24.5 uL of pooled input DNA, corresponding to up to 100 fmol total DNA, was combined in a 0.2 mL PCR tube with 0.5 uL DCS, 3.5 uL End-prep Reaction Buffer, and 1.5 uL Ultra II End-prep Enzyme Mix (NEB, Cat #E7546). Reactions were mixed by gentle pipetting and incubated in a thermocycler at 20C for 5 minutes, followed by 65C for 5 minutes. End-prepped DNA was purified using AMPure XP beads supplied with the kit at a 1.0x bead-to-sample ratio. Beads were washed with 80% ethanol according to the kit protocol. DNA was resuspended in 31 uL and transferred to a new PCR tube for adapter ligation. For ligation, the following reagents were added in order: 15 uL Ligation Buffer (LNB), 5 uL NEBNext Quick T4 DNA Ligase (NEB, Cat #M2200), and 2.5 uL Ligation Adapter (LA). Reactions were incubated for 10 minutes at room temperature. A second bead cleanup was performed using AMPure XP beads at a 0.6x bead-to-sample ratio. Instead of 80% ethanol, beads were washed with 125 uL Long Fragment Buffer (LFB). After drying for 5 minutes, DNA was eluted in 7 uL Elution Buffer (EB). Final library concentration was quantified using a tube-based Qubit HS dsDNA assay. At most, 10 fmol of the library was loaded onto a Nanopore Flongle Flow Cell (R10.4.1) according to the manufacturer’s instructions. Sequencing was performed for approximately two days or until no active pores remained.

### Nanopore data processing and analysis

Basecalled Nanopore reads were first filtered for reads that were equal to less than a 20% difference in nucleotide length from the expected amplicon size. Reads were then demultiplexed by dual-barcode identity using Cutadapt^58^ (reverse complement aware, max of 3 barcode mismatch errors tolerated). Demultiplexed reads were aligned to the reference amplicon sequence using minimap2^47^, (map-ont preset, no secondary alignments). Resulting alignments were sorted, indexed, and summarized using SAMtools^59^ and MultiQC^60^. Only samples with at least 10 mapped reads (“coverage”) were carried forward in analyses to prevent bias in variant calling due to low depth. A reference (unevolved inoculant) phage sample, barcoded and amplified as with experimental samples, served as the matched normal control for variant calling. For the T7 RNA polymerase evolutions a separate sequencing run for a control sample was used, though It is likely that variabilities seen between Nanopore runs may limit this approach in some cases.Prior to variant calling, the reference sample was downsampled to the same number of mapped reads as each evolution. Somatic variants were called using DeepSomatic v1.9.0^48^ in tumor-normal mode with the ONT model type. DeepSomatic produced per-sample variant calls in VCF format, which were then aggregated across all samples into a single table with sample identity appended for downstream analysis, and per-read sequence calling. For every variant called as a mutant by DeepSomatic within each sample, a custom script was used to analyze the frequencies of the called variant, non-mutant, or alignment errors in aggregate across all called variant positions.

### SPRI Bead Preparation

Homemade SPRI beads, analogous to standard Ampure XP beads, were prepared following the Serapure v2.2 protocol, as previously described^57^. Briefly, SPRI beads were prepared ￼ using Sera-Mag SpeedBeads (Fisher Scientific, Cat #09-981-123) suspended in a PEG-8000/NaCl binding buffer and stored at 4C protected from light. A TE solution (10 mM Tris-HCl, 1 mM EDTA, pH 8.0) was prepared in a sterile 50 mL conical tube by combining 500 uL 1 M Tris-HCl, pH 8.0, 100 uL 0.5 M EDTA, pH 8.0, and nuclease-free water to a final volume of 50 mL. SpeedBeads were washed three times prior to buffer formulation. The SpeedBead stock was mixed thoroughly by shaking and vortexing, and 1 mL was transferred to a 1.5 mL microcentrifuge tube. The tube was placed on a magnet stand until beads were fully collected, and the supernatant was removed. One milliliter of TE was added, beads were removed from the magnet and fully resuspended by vortexing, then returned to the magnet and the supernatant was removed. This wash step was repeated for a total of three washes. After the final wash, beads were resuspended in 1 mL TE and retained off the magnet. To prepare the SPRI binding buffer, 9 g PEG-8000 (Fisher Scientific, Cat #BP233) was weighed into a sterile 50 mL conical tube. Ten milliliters 5 M NaCl was added, followed by 500 uL 1 M Tris-HCland 100 uL 0.5 M EDTA. Nuclease-free water was added to approximately 49 mL total volume. The tube was mixed for 3 to 5 minutes until the PEG fully dissolved and the solution was clear upon settling. Tween 20 (27.5 uL) was then added and mixed gently. The washed SpeedBeads in TE were mixed thoroughly and transferred into the PEG/NaCl buffer. The volume was brought to 50 mL with nuclease-free water if necessary, and the suspension was mixed gently until uniformly brown. The final bead suspension was wrapped in foil and stored at 4C.

### Monte Carlo Simulation of Phage Infection Dynamics

A Monte Carlo simulation models the stochastic dynamics of M13 phage infection in a continuous-flow lagoon. The lagoon contains a population of n_molecules_ bacteria (e.g., 1000), each represented as a node on a one-dimensional lattice. Each bacterium exists in one of two states: uninfected (0) or infected (1). The lattice is indexed by position, and infection spreads between neighboring bacteria via a random walk. At each time step, three stochastic events are evaluated independently for every bacterium:

- **Infection spread**. An infected bacterium at position k can spread infection to its neighbors. A uniform random number is drawn and compared against two transition probabilities: 1) with probability p_pos_ = 1 - exp(-k_pos_ * dt), infection spreads to position k - 1 (leftward), and 2) with probability p_neg_ = 1 - exp(-k_neg_ * dt), infection spreads to position k + 1 (rightward). Here k_pos_ = k_neg_ = k1 * turn_scale, where k1 = 50 is an arbitrary base rate constant and turn_scale = 1. Interior positions (2 through n_phage_ - 1) are evaluated for spread; boundary positions do not propagate beyond the lattice edges. This represents the random-walk-like nature of phage propagation between neighboring bacteria in the lagoon.
- **Continuous flow (dilution)**. The continuous influx of fresh, uninfected bacteria into the lagoon is modeled by independently reverting each bacterium to the uninfected state with probability p_flow_ = 1 - exp(-k_flow_ * dt), where k_flow_ = k_pos_ * flow_scale. The flow_scale parameter (typically 0.25-1.0) sets the dilution rate relative to the infection rate, corresponding to lagoon volume turnovers per unit time.
- **Time step**. The simulation time step is set small relative to the fastest rate: dt = 1 / (k_pos_ + k_neg_ + k_flow_) * 0.01. This ensures that the probability of multiple events occurring in a single step is negligible, preserving the accuracy of the stochastic simulation.

At time zero, a small number of bacteria in the lattice are initialized as infected, simulating the multiplicity of infection (MOI) at inoculation. All remaining bacteria start as uninfected. At each time step, two quantities are recorded: (i) the total number of infected bacteria across the entire lagoon (B_total_), and (ii) the total number of infected bacteria excluding the initially seeded positions (B_spread_), which isolates de novo infection from the seed. Simulations are run across combinations of population size (n_phage_), number of independent Monte Carlo replicates per bacterium (n_molecules_), and flow_scale values to characterize the dependence of infection dynamics on these parameters. Each combination is simulated for a duration of 3 time units. The simulation is implemented in Python using vectorized NumPy operations, and figures are generated in R using ggplot2.

### Turbidostat and Chemostat Simulations

Growth dynamics under six culture control strategies were simulated using Monod kinetics. Biomass concentration X (measured as optical density, OD) and limiting substrate concentration S were modeled as coupled ordinary differential equations:

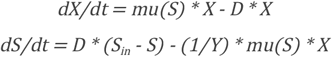

where the specific growth rate follows the Monod equation: *mu*(*S*) = *mu*_*max*_ ∗ *S* / (*K*_*s*_ + *S*). Parameters were set to mu_max_ = 0.8 hr ^-1^, K_s_ = 1.0, yield coefficient Y = 0.5 OD per unit substrate, feed substrate concentration S_in_ = 4.0, and target OD setpoint = 0.6. All cultures were initialized at OD = 0.006 (equivalent to a 1:100 dilution from setpoint) with substrate at S_in_. These equations were integrated using Euler’s method with a time step of 0.01 hr for continuous models and 0.001 hr for batch growth segments in discrete-dilution models. The simulations assume a single limiting substrate, well-mixed culture volume, no cell death or maintenance energy, no lag phase, and instantaneous dilution events. For the continuous turbidostat, D was set to zero during initial growth and engaged once OD first reached the setpoint. At each subsequent time step where OD exceeded the setpoint, a fraction of the culture volume was instantaneously removed and replaced with fresh medium to restore OD to exactly 0.6, with substrate concentration adjusted proportionally:

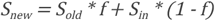

where f = OD_setpoint_ / OD_current_ is the fraction of culture volume retained. This was repeated across five growth rates (mu_max_ = 0.6, 0.8, 1.0, 1.2, and 1.4 hr^-1^). For the chemostat, a constant dilution rate D was applied continuously once the culture reached the setpoint. This was simulated across five dilution rates (D = 0.5, 0.6, 0.7, 0.8, and 0.9 vol/hr) at each of the five growth rates to illustrate the interaction between growth rate and dilution rate on steady-state OD. The Monod steady-state biomass concentration is given by: X* = Y * (S_in_ - K_s_ * D / (mu_max_ - D)), which predicts washout (X* = 0) when D >= mu_max_ and lower equilibrium OD when D is suboptimal. Serial dilution was modeled as unrestricted batch growth punctuated by 1:100 dilutions into fresh medium every 8 hours (f = 0.01), with both OD and substrate reset proportionally at each transfer. The batch maximum OD is bounded by X_max_ = Y * S_in_ = 2.0, representing complete substrate consumption. Three semi-continuous turbidostat strategies were compared, all using the same proactive dilution framework. At each measurement time (t_i_), the system projects growth forward by simulating Monod batch kinetics over the interval dt to predict OD at t_i_ + dt. If the projected OD would exceed the setpoint, the system dilutes immediately to a target OD_low_ such that subsequent growth over dt returns OD to exactly the setpoint. The value of OD_low_ is found by binary search (50 bisection iterations) over the function:

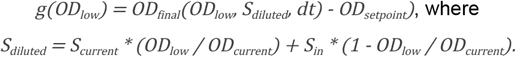

This approach begins dilution before OD reaches the setpoint, preventing overshoot. A physical constraint of the 150 uL culture volume limits the maximum dilution to two-fold across all semi-continuous modes (minimum fraction retained f >= 0.5). In practice, only the non-adaptive mode encounters this limit, as the adaptive and fixed-interval modes calculate dilution depths that remain within this range. In the fixed-interval (discrete) turbidostat, measurements occur at regular 30-minute intervals and the system knows the true growth rate, enabling exact calculation of dilution depth at each cycle. The adaptive turbidostat uses measurement intervals drawn uniformly at random from U(20, 60) minutes. Because the system knows both the growth rate and the actual duration of the upcoming interval, it solves for the correct dilution depth at each cycle despite the variable timing. The non-adaptive turbidostat experiences the same random measurement intervals as the adaptive mode (using an identical random seed for direct comparison) but calculates dilution depth assuming a fixed 30-minute interval. When the actual interval exceeds 30 minutes, the culture overgrows the setpoint; when it is shorter, the culture undershoots. Because this mode routinely encounters the two-fold maximum dilution constraint, it cannot compensate for large mismatches between assumed and actual interval duration, leading to sustained overshoot at higher growth rates. All simulations were run in R.

## Notes

### Summary of Updates

Updated the COI and and funding statements which had previously been included in the manuscript file and did not auto-populate (as in previous BioRxiv submissions). Two in-text figure references were corrected, but no changes were made to the manuscript file (which already contained these declarations).

